# PepPre: Promote Peptide Identification Using Accurate and Comprehensive Precursors

**DOI:** 10.1101/2023.05.13.540645

**Authors:** Ching Tarn, Yu-Zhuo Wu, Kai-Fei Wang

## Abstract

Accurate and comprehensive peptide precursor ions are crucial to tandem mass spectrometry-based peptide identification. An identification engine can greatly benefit from the search space reduction hinted by credible and detailed precursors. Additionally, both the number of identifications and the spectrum explainability can be increased by considering multiple precursors per spectrum. Here, we propose PepPre, which detects precursors by decomposing peaks into multiple isotope clusters using linear programming methods. The detected precursors are scored and ranked, and the high-scoring ones are used for the following peptide identification. PepPre is evaluated both on regular and cross-linked peptides datasets, and compared with 11 methods in this paper. The experimental results show that PepPre achieves 203% more PSM and 68% more peptide identifications than instrument software for regular peptides, and 99% more PSM and 27% more peptide pair identifications for cross-linked peptides, which also outperforms all other evaluated methods. In addition to the increased identification numbers, further credibility evaluation evidence that the identifications are credible. Moreover, by widening the isolation window of data acquisition from 2 Th to 8 Th, the engine is able to identify at least 64% more PSMs with PepPre, demonstrating the potential advantages of large isolation windows.

**Graphical TOC Entry:** 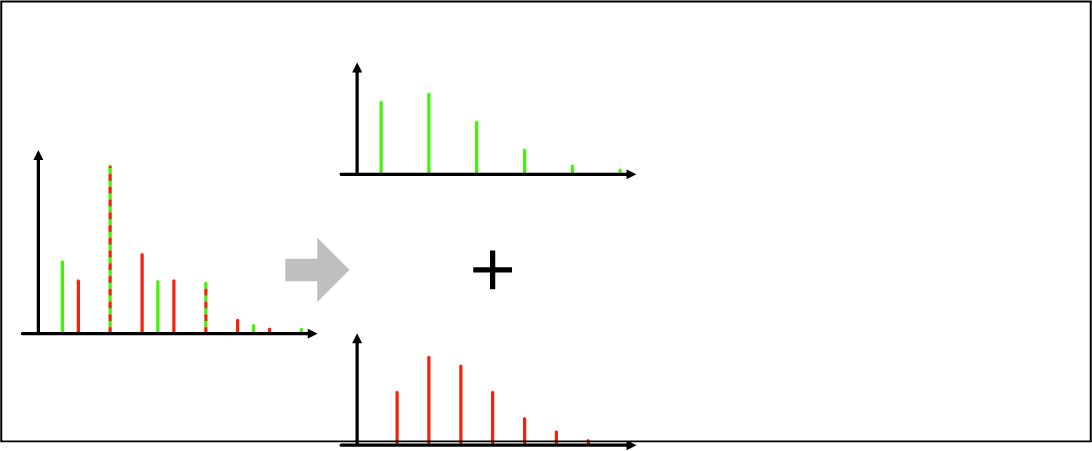

## Introduction

Mass spectrometry (MS)-based peptide identification has long been of vital importance in proteomics and related fields,^1^ and has also spawned a large number of techniques and tools. Taking a conventional bottom-up protein identification workflow as an example, proteins are first digested into peptides and then separated by liquid chromatography. The peptides are charged and analyzed by a mass spectrometer, which measures mass-to-charge ratios (m/z) and abundance values of the peptides and generates MS1 scans. Since a peptide molecule of the same amino acid sequence can be composed of different isotopes or be differently charged, one peptide may be observed as several isotope clusters, where a cluster consists of molecules of the same charge state but of various isotopic compositions, and also corresponds to a peptide precursor ion.

As the peptides are eluted sequentially, they are further selected by isolation windows and fragmented into MS/MS (MS2) scans separately. For a typical data-dependent acquisition (DDA), a mass spectrometer usually prefers to target abundant peaks as the activation center, and to include and fragment the whole isotope cluster, it will isolate all peaks within a certain radius, say 2 Th. Notably, the mechanism leads to the possibility that one MS2 consists of multiple peptides. Besides the peptide targeted as the isolation center, there can exist other co-eluted peptides whose m/z values are close to the targeted one, and consequently, the peptides are co-fragmented into the same MS2 scan. The co-fragmentation is common, especially for complex samples,^2^ and can increase the complexity of spectra but also makes it possible to identify extra peptides.

Peptide identification can greatly benefit from precursors.^3^ In many applications, engines search against huge databases, various modifications, and complex proteolysis, and thus it is not feasible to compare a spectrum with all candidate peptides.^4,5^ Therefore, additional information is required to reduce the search space and improve identification efficiency. Such information includes the mass and charge state of precursor ions, retention time, etc., of which the mass and charge state are usually the most accurate and effective. Additionally, the properties can also play crucial roles in verifying identification results.^6,7^

The importance and challenges of peptide precursor detection make it attract the attention of many researchers. Most of the existing methods can be grouped into two categories, detecting precursors directly and (2) detecting peptide features based on the liquid chromatography with mass spectrometry (LC-MS) map.

The first type of precursor detection method mainly focuses on analyzing the isotopic pattern. For example, some instrument software tries to estimate the charge state and shift m/z of the activation center to the monoisotopic mass based on the averagine model.^8^ The correction may be rough and the co-eluted precursor ions are ignored. Some engines, such as MS-GF+^9^ and xiSearch,^10,11^ make further corrections, which can subtract 1, 2, or 3 Da from the original masses respectively to ensure that the monoisotopic masses are included. The precursors obtained by enumerating are searched separately, and isotopic patterns of them are not further evaluated, which can introduce potential mistakes, even though generally only the best one of each spectrum is preserved. THRASH^12^ is a standalone precursor detection algorithm, which compares a peak cluster with theoretical isotopic patterns using least-squares fitting and removes matched peaks from the spectrum recursively. Unfortunately, least-squares fitting can not effectively distinguish two precursors with a mass difference of about 1 Da. Park et al.^13^ purposed a method based on the intensity ratio of adjacent peaks, which is able to deisotope overlapping clusters,^14,15^ but it is still not robust enough to handle complex cases like Figure 1. RawConverter^16^ corrects precursors using linear programming, which is able to deisotope highly overlapping clusters. However, the scoring method of RawConverter still assumes that one peak is only composed of one peptide, which limits its performance. Besides, for DDA data, RawConverter does not consider co-fragmented precursors and only exports one precursor per MS2. Differently, pParse^17^ estimates all possible combinations of candidate peaks and charge states, and is able to detect co-eluted precursor ions. However, since the candidate precursors are estimated individually, pParse can not effectively handle overlapping isotope clusters. It also requires a large amount of labelled data to train its model, resulting in a lack of robustness and interpretability.

**Figure 1:**
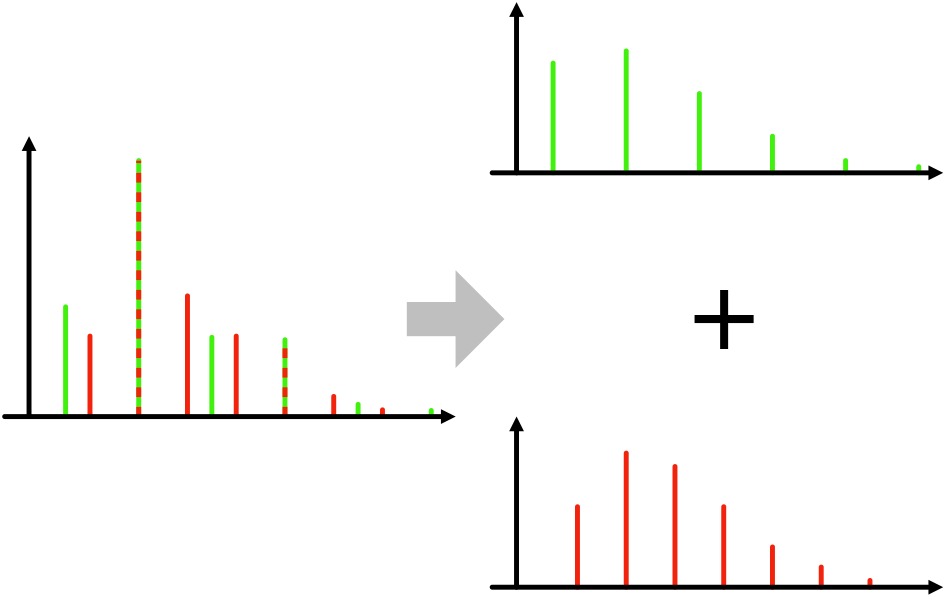
A set of peaks that consists of two isotope clusters. Since some peaks of the clusters are merged, the isotope clusters are weirdly shaped and thus resemble implausible, even though the peaks are free of noise.

Instead of detecting peptide precursor ions of an MS2 scan directly, many methods^18–26^ try to extract all peptide features of the LC-MS map. The methods report the mass, charge state, and retention time of each peptide feature. Specifically, MaxQuant^22^ assembles peaks of each scan into three-dimensional (3D) peaks, constructs a graph with the 3D peaks based on the correlation between the masses and retention time ranges of them, and then splits the graph into connected subgraphs. Compared to MaxQuant, Dinosaur^24^ uses the averagine model to determine monoisotopic masses and to discard incredible features. DeepIso^25^ and PointIso^26^ apply recent artificial intelligence technologies. DeepIso builds modules based on neural networks to detect peptide features. PointIso converts mass spectra into point clouds and achieves better performance than DeepIso.

The feature-based methods can be infeasible if a continuous LC-MS map is not available. More importantly, the feature detection methods may suffer from serious performance issues since they aim to find all features from the whole LC-MS map.^24,25^ The computing costs can be unaffordable in some applications, and, in fact, only a small number of detected features are targeted for MS2.^27^ And thus, it is a good choice to focus on deisotoping the spectrum slice where the peptides are fragmented.

Researchers also explored wide window acquisition (WWA) in many works,^28–30^ including Chimerys. In this work, we mainly focus on precursor detection instead of comparing different acquisition or analysis methods, and thus related topics are not further discussed.

Here we propose a new peptide precursor detection method PepPre, which does not rely on peptide features. It assumes that a spectrum is a weighted linear combination of candidate isotope clusters, where the candidates can be generated by enumerating each possible peak and charge state. Unlike most existing approaches, PepPre does not evaluate candidate isotope clusters individually or recursively but instead attempts to interpret the spectrum, which makes PepPre can handle challenging cases such as heavily overlapped ones. In practice, a linear optimization algorithm can be applied to calculate the weight of each candidate, so that the theoretical combination of candidates can fit the experimental spectrum. The isotopic pattern of each candidate can be further evaluated even if some peaks are shared by multiple precursors. Naturally, candidate precursors with significant weights and suitable isotopic patterns are considered very credible and more likely to be identified. Moreover, we also release a visualization tool called PepPreView to help with data analysis.

PepPre is implemented with Julia Language,^31^ which is designed for high performance and is easy to use as a scripting language. JuMP^32^ and GLPK are used to solve the problem. The source code is open, and we also provide executables with a graphic user interface and command line tools for multiple platforms including Linux, macOS, and Windows. Users are able to use the software directly or modify the source code for specific purposes. The resources can be found at http://peppre.ctarn.io. We look forward to seeing the software serve the community in the long term.

## Methods

### Candidate Selection

In the natural environment, for a digested peptide, although the relative abundance of monoisotopic peak decreases as the number of atoms increases, the peak is usually detectable by mass spectrometers. For example, for a peptide with a mass of 4000 Da, the theoretical relative abundance of the monoisotopic peak is about 9%, which is about 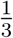 of the most abundant peak (see http://averagine.ctarn.io/?mass=4000). Therefore, a candidate precursor ion list can be constructed by enumerating possible monoisotopic peaks and charge states, and the peaks are supposed to be located in or near the isolation window.

It is not enough to only consider peaks in the isolation window. As illustrated in Figure 2a, in addition to the monoisotopic peaks of red clusters, which are located in the isolation window, the monoisotopic peaks of brown clusters are outside the window. However, since the tails of these brown clusters are still inside the window, some of the peptide molecules can also be isolated and fragmented into the MS2 scan. Such peptides are identifiable too, and thus should be considered. Apart from this, no part of the green clusters is inside the isolation window, and the respective peptides are not fragmented into the MS2 scan. Even though, since the green clusters share some peaks with the brown or red clusters, adding them to the candidate list can improve the accuracy, and thus the clusters should also be included.

**Figure 2:**
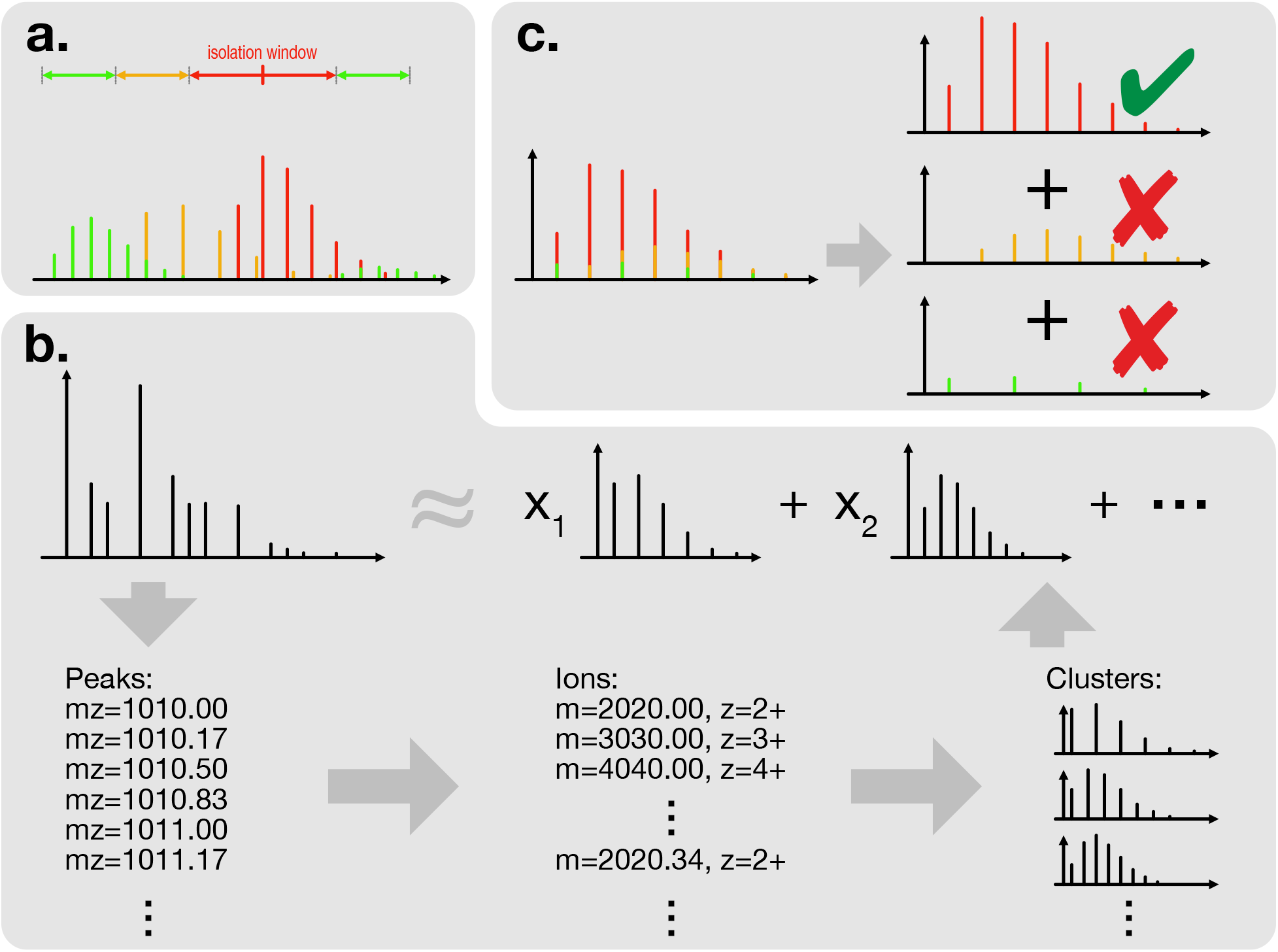
a. Several types of isotope clusters that should be considered when decomposing a spectrum slice. The monoisotopic peak of the red cluster is located in the isolation window, while the monoisotopic peak of the brown cluster is not. However, some isotopic peaks of the brown cluster are also included and fragmented into the respective MS2 scan, making the respective precursor ion identifiable. The green clusters are not fragmented but can share peaks with fragmented ones. Thus, all three types of clusters should be properly considered to improve the accuracy and coverage of the solution. b. The workflow of spectrum decomposing. We first select potential monoisotopic peaks from the spectrum slice, and combine them with possible charges to construct a list of candidate peptide precursor ions. The theoretical isotope cluster of each candidate ion is estimated using methods like averagine model. We assign an unknown weight, or abundance value, to each candidate ion, and add them up. The sum should be as close as possible to the experimental spectrum slice. The unknowns are further solved by methods such as linear programming. c. An isotope cluster split into parts. Due to noise, intensity variations, inaccurate estimation, etc., the theoretical isotope clusters do not exactly match the experimental clusters, and thus some unexpected ions can be reported. They are usually relatively low in abundance, and can be dynamically excluded during decomposition.

In practice, given a isolation window [*mz*_min_, *mz*_max_], all peaks in range [*mz*_min_−2 Th, *mz*_max_+ 1 Th] are considered as candidate monoisotopic peaks, and green clusters are filtered out automatically by the scoring function, while brown clusters generally earn lower scores than red clusters.

Instead of requiring the full chromatograms, PepPre only uses the two nearest MS1 scans of each MS2 scan, that is, the master MS1 scan and the MS1 scan after the master scan. The MS1 scans are decomposed individually, and the results are merged. Notably, PepPre can process the single master MS1 scan only, and the extra MS1 scan is optional.

Additionally, for a candidate cluster, if its 2nd peak, i.e., the isotopic compositions with one extra neutron more than the monoisotopic composition, is not observed in the MS1 scan, the cluster is considered less credible and would be removed directly. The prefiltering strategy reduces the computational cost greatly.

### Spectrum Decomposing

As shown in Figure 2b, considering spectrum *s* and a specified isolation window, we list the peaks in and near the window as {(*mz*_1_, *a*_1_), (*mz*_2_, *a*_2_), …, (*mz*_*m*_, *a*_*m*_)}, where *mz*_*i*_ and *a*_*i*_ are the mass-to-charge ratio (m/z) and abundance value of *i*-th peak respectively, and then the spectrum can be represented as s = [*a*_1_, *a*_2_, …, *a*_*m*_]. Since the monoisotopic peak and charge state can uniquely identify an isotope cluster or a precursor ion, we can easily build a list of all potential precursor ions by enumerating every possible peak and every possible charge state. Using methods like averagine model, for each candidate precursor ion, the mass values and relative abundances of the isotopic peaks can be estimated, and thus, the isotope cluster can also be represented as a vector like a spectrum. Particularly, if the m/z value of an estimated isotopic peak does not appear in spectrum *s*, we add a virtual zero-abundance peak to spectrum *s*. Let’s denote the vector of *i*-th candidate precursor ion *p*_*i*_ as p_*i*_. Since the spectrum is composed of isotope clusters, we have

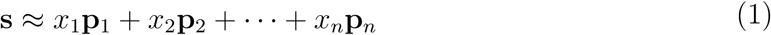

where *n* is the number of candidates, and *x*_*i*_ is the unknown abundance of *i*-the candidate precursor ion. The equation can be rewritten as below,

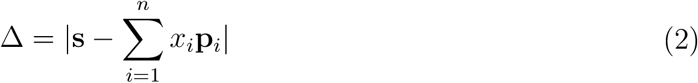

and then we can solve it by minimizing Δ.

Using the L1 metric, that is, minimizing Δ by minimizing the sum of each dimension of Δ, we have

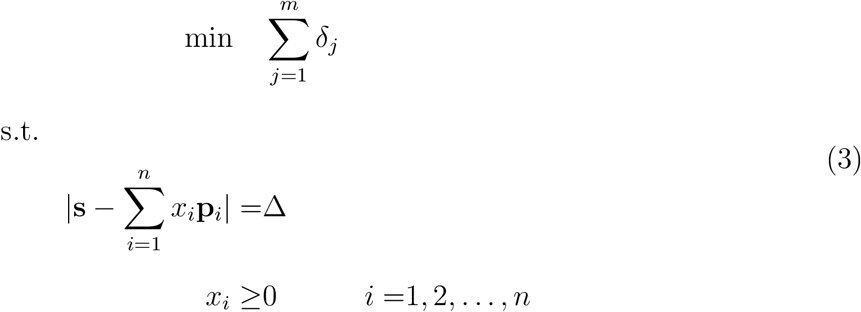

where Δ = [δ_1_, δ_2_, …, δ_*m*_].

The equation is equivalent to

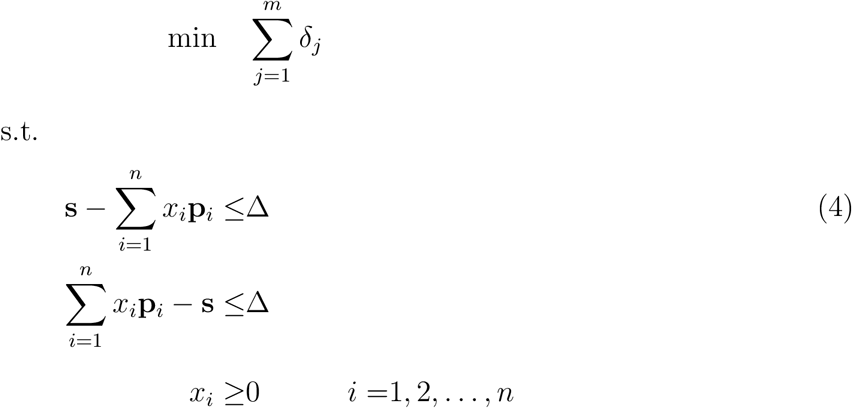

which can be solved with linear programming methods. The candidate precursor ions can be further easily evaluated and compared based on the solution.

### Stepped Dynamic Exclusion

Due to noise, measurement errors, estimating errors of isotope clusters, etc., the theoretical clusters and experimental clusters do not exactly match, which leads to some non-existent precursor ions being erroneously assigned low scores and reported. The incorrect ions usually share many peaks with others. For example, as shown in Figure 2c, the brown cluster, whose mass value is greater than the red one by about 1 Da, and the green cluster, whose charge state is half of the red one, can be reported by mistake. Therefore, during decomposing, a stepped dynamical exclusion strategy is applied to improve the results.

More specifically, a spectrum is iteratively decomposed using current candidates, and after each iteration, some candidates are removed based on the following criteria, until no candidates can be excluded anymore.

For each cluster *a*, it is compared with each other cluster, and if and only if exists cluster *b* satisfies the following conditions, the cluster *a* is excluded.

1. Many peaks of cluster *a* are shared with cluster *b*, that is, the value of charge state of *b* is divisible by that of *a* (e.g., a is 3+ and b is 6+), and the monoisotopic peak of *a* is also an isotopic peak of *b*.
2. Cluster *a* is less abundant than cluster b by threshold τ, that is, *x*_*a*_ *<* τ *x*_*b*_.

Instead of applying an adventurous threshold directly, we increase the threshold step by step to make it more stable. Given a threshold τ, 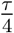 is applied at first, and when the exclusion stops, we increase the threshold to 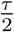, and then increase it to τ at last.

To make the decomposing procedure faster, after each iteration, The less credible candidates, whose weight *x*_*_ or matching scores are less or equal to zero, are also excluded. The matching score is described in detail below.

### Precursor Evaluation

Two types of scores are used to evaluate candidate precursor ions, the abundance score and the matching score. The abundance score is defined as precursor ion fraction (PIF), calculated in the following way.

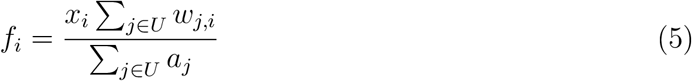

where *U* is the set of indices of all peaks in the isolation window, and *w*_*j,i*_ is the relative abundance of candidate ion *i* corresponding to *j*-th m/z value, that is, **p**_*i*_ = [*w*_1,*i*_, *w*_2,*i*_, …, *w*_*m,i*_]. Specifically, the raw intensities of peaks are used, without further correction based on isolation efficiency profile.

For the matching score of a candidate cluster, we define it as the error rate of corresponding peaks as below. The score considers shared peaks properly.

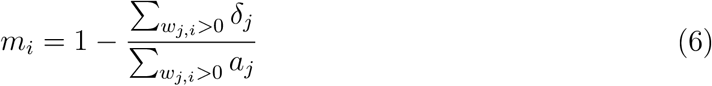

where the selector *w*_*j,i*_ *>* 0 is used to determining which peaks are associated with the candidate cluster *i*.

The product of *f*_*i*_ and *m*_*i*_ is used as the final score of candidate precursor ion *i*. The results of multiple MS1 scans are merged by adding up scores of the same ions. The merged candidate precursor ions are sorted in descending order of score, and the ones with high scores are preserved. Typically, the number of precursor ions to be preserved is controlled by a fold of the number of MS2 scans. For example, 2-fold exporting indicates that two precursor ions per MS2 scan are exported on average.

### Datasets and Settings

The method, PepPre, has been evaluated on both regular peptides and cross-linked peptides, and compared with pParse,^17^ RawConverter,^16^ Monocle,^35^ Decon2LS,^36^ RAPID,^37,38^ MaxQuant,^22^ Dinosaur,^24^ PointIso,^26^ and three enumeration-based methods (EnumInst, EnumIW, EnumEx), as shown in Table 1 and Table 2. The datasets are searched with pFind^4^ and pLink^5^ respectively.

**Table 1:** Datasets

| name | isolation window | description |
| --- | --- | --- |
| Zubarev-Human-2<sup>33</sup> | 2 Th | regular peptide, human |
| Zubarev-Human-4<sup>33</sup> | 4 Th | regular peptide, human |
| Zubarev-Human-6<sup>33</sup> | 6 Th | regular peptide, human |
| Zubarev-Human-8<sup>33</sup> | 8 Th | regular peptide, human |
| Dong-DSS-1.6<sup>34</sup> | 1.6 Th | DSS-linked, synthetic |

**Table 2:** Baseline Methods

| name | version | co-eluted ions? | description |
| --- | --- | --- | --- |
| pParse<sup>17</sup> | 2.2 | Yes | averagine model, trained with labeled data |
| RawConverter<sup>16</sup> | 1.2.0.1 | No | averagine model, linear programming |
| Monocle<sup>35</sup> | 0.3.51 | No | averagine model |
| Decon2LS<sup>36</sup> | 1.1.7787 | Yes | averagine model, recursively subtraction |
| RAPID<sup>37,38</sup> | 1.15 | Yes | intensity ratio, recursively subtraction |
| MaxQuant<sup>22</sup> | 2.0.3.0 | Yes | peptide feature, 3D peak |
| Dinosaur<sup>24</sup> | 1.2.0 | Yes | peptide feature, averagine model |
| PointIso<sup>26</sup> | 28110df | Yes | peptide feature, artificial intelligence |
| EnumInst | - | No | subtract 1, 2, 3 Da from instrument-exported ions |
| EnumIW | - | Yes | enumerate all peaks in isolation window |
| EnumEx | - | Yes | enumerate all peaks in and near isolation window |

For Dong-DSS-1.6, 16 RAW files provided by the authors are used in this work. The dataset is larger than others we know, e.g., the dataset from Mechtler lab,^39^ and thus is more suitable for our evaluation.

For regular peptide identification, Zubarev-Human-2/4/6/8^33^ are applied, which consist of four single shotgun proteomics runs with a two hours gradient. As the names indicate, the isolation window sizes of these runs vary from 2 Th to 8 Th. pFind (version 3.1.6) is applied as the identification engine and set to open search mode. The FDR is set to 1% at the PSM level.

For cross-linked peptide pair identification, Dong-DSS-1.6^34^ is applied. The dataset consists of synthetic peptides cross-linked by DSS, and thus, credibility evaluation based on entrapment databases is additionally available to validate the identifications. pLink (version 2.3.9) is used to identify the cross-linked peptide pairs. Fixed modification cysteine carbamidomethylation and variable modification methionine oxidation are set respectively. The FDR is set to 1% at the PSM level.

Databases of corresponding organisms are downloaded from UniPort on Nov. 19, 2021, and we use the first 100 *E. coli* proteins as the trap database for Dong-DSS-1.6.

Since pParse can not control the export fold directly, we vary the scoring threshold with the step of 0.1. The isolation window is set manually since pParse can not extract it automatically. For MaxQuant, precursors are extracted from the .apl files and converted into .mgf files using scripts. The .mzML files required by Dinosaur are generated using MsConvert (version 3.0.21304-6ca6020). PointIso is downloaded from GitHub, and the signature of the latest commit is 28110df.

For EnumInst, we additionally consider ions whose mass values are less than the original instrument exported precursors by 1 Da, 2 Da, and 3 Da as calibration. Therefore, it generates four types of results, whose number of peptide precursors are 1-fold, 2-fold, 3-fold, and 4-fold respectively. For EnumIW, we consider the charge state range 2+ to 6+, and enumerate all peaks in isolation window as monoisotopic peaks. EnumEx is similar to EnumIW, but all peaks in [*mz*_min_ − 1 Th, *mz*_max_] are included, where [*mz*_min_, *mz*_max_] is the isolation window.

The settings not mentioned above are set as default respectively. For methods based on peptide features, if no matching process is embedded, i.e., Dinosaur and PointIso, the features are matched with MS2 scans as DeMix.^33^ Specifically, DeMix assigns a feature to an MS2 scan if the m/z value of the feature is within the isolation window of the scan and the retention time of the scan is within the range of the feature. Decon2LS and RAPID process the full MS1 scan, and a similar matching process is applied.

## Results

### Identification and Validation

Figure 3 reports the identification numbers at PSM level and peptide (pair) level on Zubarev-Human-2 (3a), Zubarev-Human-8 (3b), and Dong-DSS-1.6 (3c) respectively. Additionally, on Dong-DSS-1.6, credibility evaluation based on the entrapment database is further applied. The results show that PepPre achieves more identifications at both the PSM level and peptide (pair) level by exporting significantly fewer precursor ions than others. The gap is further widened as the isolation window size increases, especially when compared with pParse.

**Figure 3:**
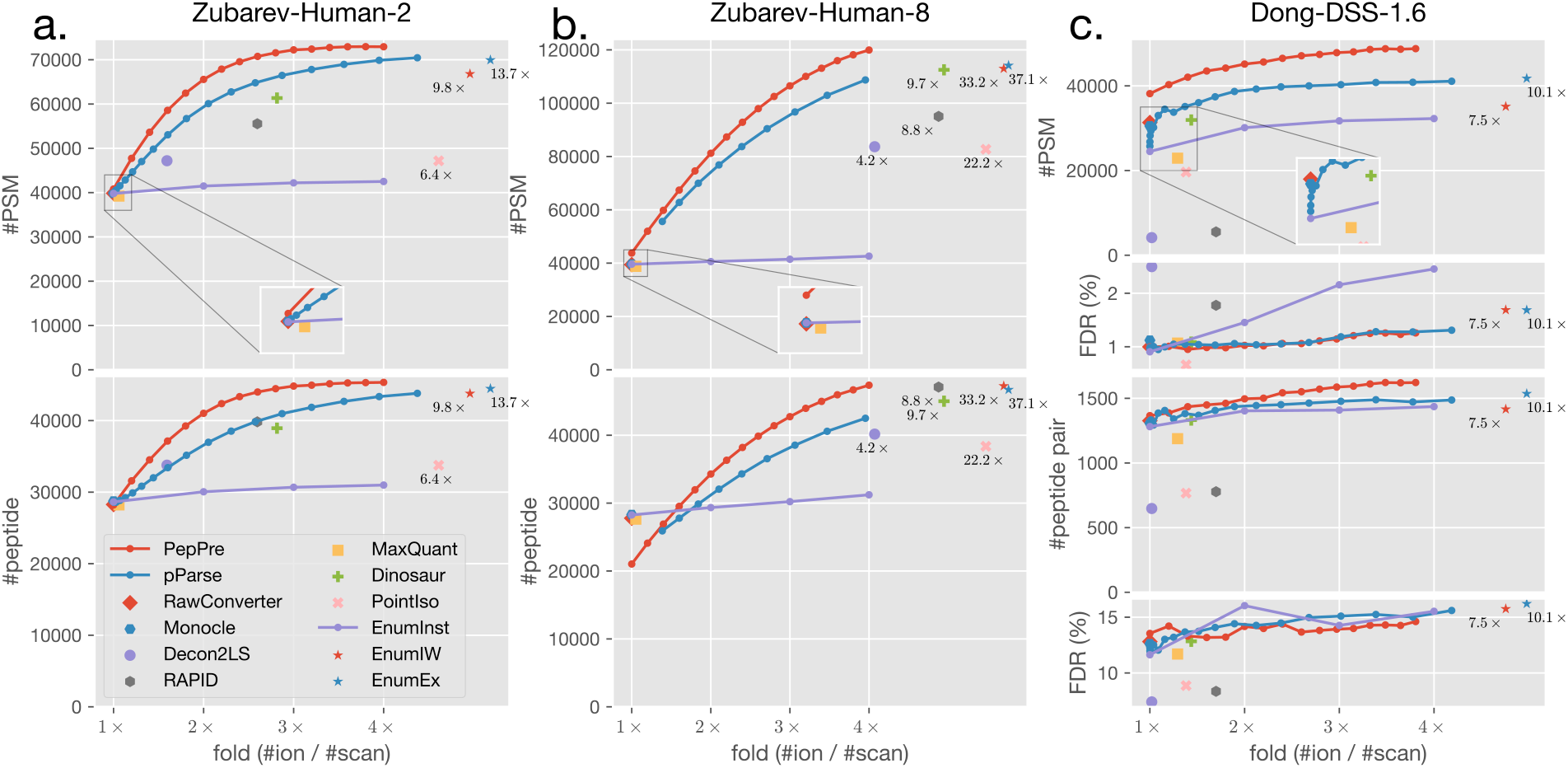
The identified PSMs and peptides or peptide pairs on Zubarev-Human-2 (a), Zubarev-Human-8 (b), and Dong-DSS-1.6 (c). The results show that PepPre achieves more identifications than other methods on both regular peptides and cross-linked peptides datasets. Additionally, on the synthetic cross-linked peptides dataset, the results reported by PepPre are as credible as others.

For example, when the methods (PepPre, pParse, and EnumInst, since others are not configurable) are set to exporting 4-fold precursor ions, on Zubarev-Human-8, whose isolation window size is set to 8 Th, PepPre achieves about 203% more identified PSMs and 68% more identified peptides than the uncorrected instrument exported ions (1-fold of EnumInst), and 10% more PSMs, 11% more peptides than pParse. On Dong-DSS-1.6, whose isolation window size is set to 1.6 Th, PepPre achieves about 99% more identified PSMs and 27% more identified peptide pairs than the uncorrected instrument exported ions, and 19% more PSMs, 9% more peptide pairs than pParse. PepPre outperforms all other baseline methods including EnumEx, which exports 13.7-, 37.1- and 10.1-fold ions on the three datasets respectively.

The calculated FDR values by evaluation using entrapment databases also show that the identifications are equally credible. It seems weird that the calculated FDR of the peptide pair level is pretty high, even if only the original precursors by instrument software are exported. Actually, the high FDR is caused by the fact that only the FDR on the PSM level is controlled, and it is very common in our similar research.

Notably, PepPre achieves fewer peptide identifications at 1-fold on Zubarev-Human-8, compared with many other methods. The reason is that the ranking of precursors is global, and thus many MS2 scans export no precursor, while many methods always export at least one precursor per MS2. The global ranking increases the number of PSMs, but the possibility that two PSMs are of the same peptide also increases. We additionally provide the option to force exporting the original precursor ion to guarantee that at least one precursor per MS2 is exported, named PepPre+. The results are shown in Figure S2.

It is important to note that since pParse uses a threshold of scoring to control the number of precursor ions to be exported, and instead of setting the fold directly, we try different thresholds in steps of 0.1, and thus the folds are not exactly equal to 4. In this case, pParse exports 3.95-fold ions on Zubarev-Human-8 and 4.19-fold ions on Dong-DSS-1.6. PepPre does not export 4-fold ions on Dong-DSS-1.6 either, since only 3.81-fold ions are left after decomposition. The following experiments are the same as this.

We compare PepPre and the baseline methods respectively further, especially for the overlapped identification, and the results are shown in Figure S4-S14. For example, compared with EnumEx, the major part (58.4%, 90.2%, and 44.5% respectively on the three datasets) of the PSMs uniquely identified by PepPre, are also exported by EnumEx. The precursors exported by EnumEx nearly include all possible candidate precursor ions. However, the engines fail to identify the precursor ions, which results in PepPre even outperforming EnumEx, showing the negative impact of too many incorrect precursor ions. Additionally, Table S7-S10 suggest that the co-eluted ions are the major part of increased identifications compared with many methods. The number and sequence coverage of identified protein groups are also shown in Figure S3.

To further evaluate the quality of identification, we calculate the number of observed fragments of each identification. Only the *b*-ions and *y*-ions are considered. For regular peptides, the candidate charge states are {+1, +2}, and for cross-linked peptides, {+1, +2, +3} are considered instead. The results are shown in Figure 4, indicating that PepPre achieves more high-quality identifications.

**Figure 4:**
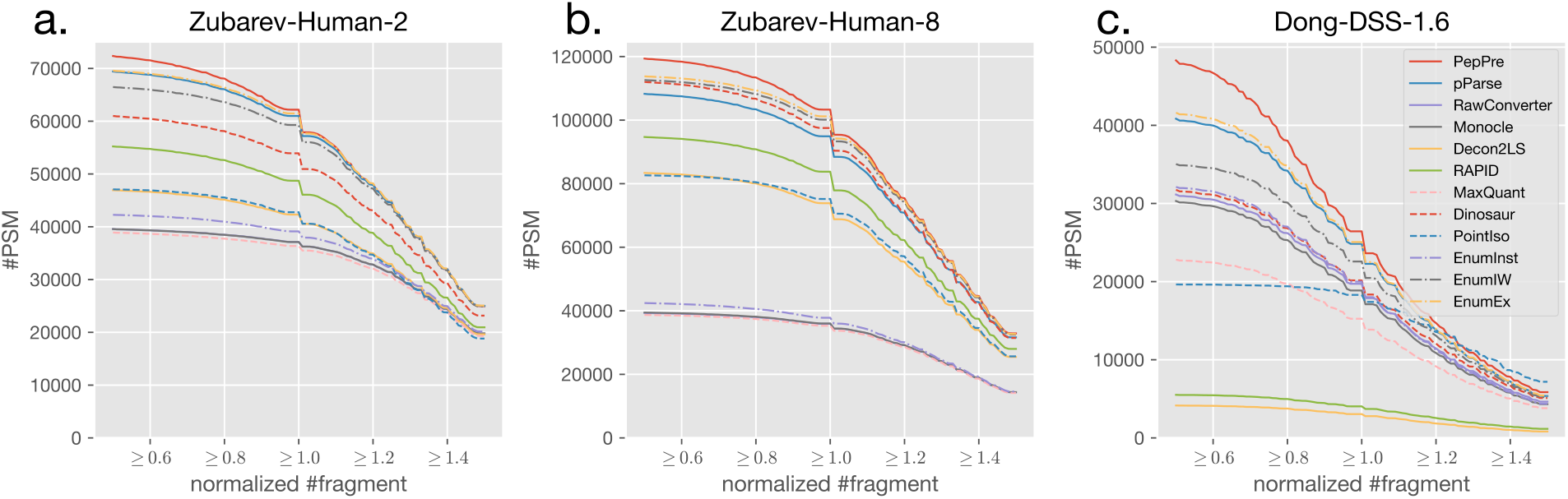
The number of identified PSMs that are well matched to respective spectra. PepPre, pParse, and EnumInst are all set to 4-fold, while the fold can not be specified for other methods. The number of fragments is normalized by dividing the number of amino acids of the respective sequence. For example, with a lower bound of 0.8, PepPre, pParse, and EnumInst enable the engine to identify 38034, 34190, and 27034 PSMs on Dong-DSS-1.6 respectively, of which the number of observed fragments of each PSM is not less than 0.8*n*, where *n* is the number of amino acids of the respective sequence.

### Charge State and Mass Coverage

Figure 5 reports the distributions of the charge states and masses of identified PSMs. It shows that for almost all charge states and mass ranges, the number of identified PSMs of PepPre is more than pParse, EnumInst, EnumIW, and EnumEx, which indicates that PepPre has no weakness in various charge states or masses. Differently, since pParse requires a lot of data to train, its performance also heavily relies on and is limited by the quality and characteristics of the training data. The figure and Table S1-S6 shows that cross-linked peptide pairs carry more charge than regular peptides, and because pParse is trained with regular peptides, PepPre achieves more highly-charged PSMs than pParse. Specifically, on Dong-DSS-1.6 and the fold is set to 4, the 5+ PSMs identified by PepPre are 38% more than pParse, and the 6+ PSMs identified by PepPre are 21% more than pParse, while the average increment is about 19%. The details of other baselines are shown in Table S1-S6.

**Figure 5:**
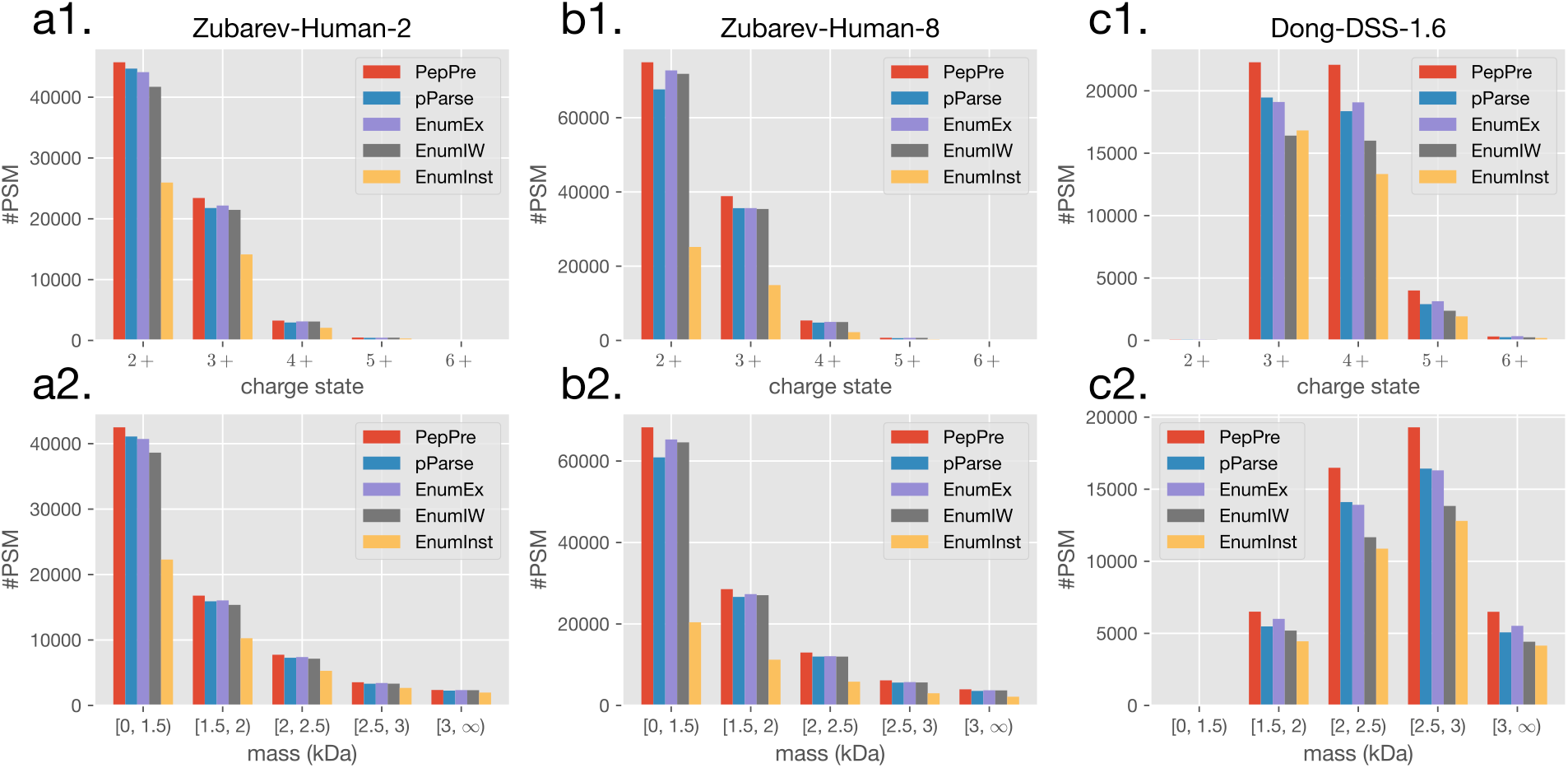
The charge states and masses of identified PSMs on Zubarev-Human-2 (a1, a2), Zubarev-Human-8 (b1, b2), and Dong-DSS-1.6 (c1, c2). The results show that for almost all charge states and mass ranges, PepPre achieves more identifications than pParse and the three enumeration-based methods.

### Cofragmented Precursor with Overlapping Cluster

One of the major advantages of PepPre is that it can decompose complex and heavily overlapping clusters more effectively. For each identified precursor ion, we further check whether its theoretical isotope cluster shares peaks with any others in the same MS2. As illustrated in Figure 6, PepPre enables 8881 identified PSMs with overlapping clusters, which is about 134% more than 3791 identified PSMs reported by pParse. These gains account for 69% of the overall increment, indicating that they should be valued. Additionally, the calculated FDR also shows that heavily mixed precursor ions reported by PepPre are equally credible than other precursor ions, while the FDR value of such ions reported by pParse is significantly higher than expected.

**Figure 6:**
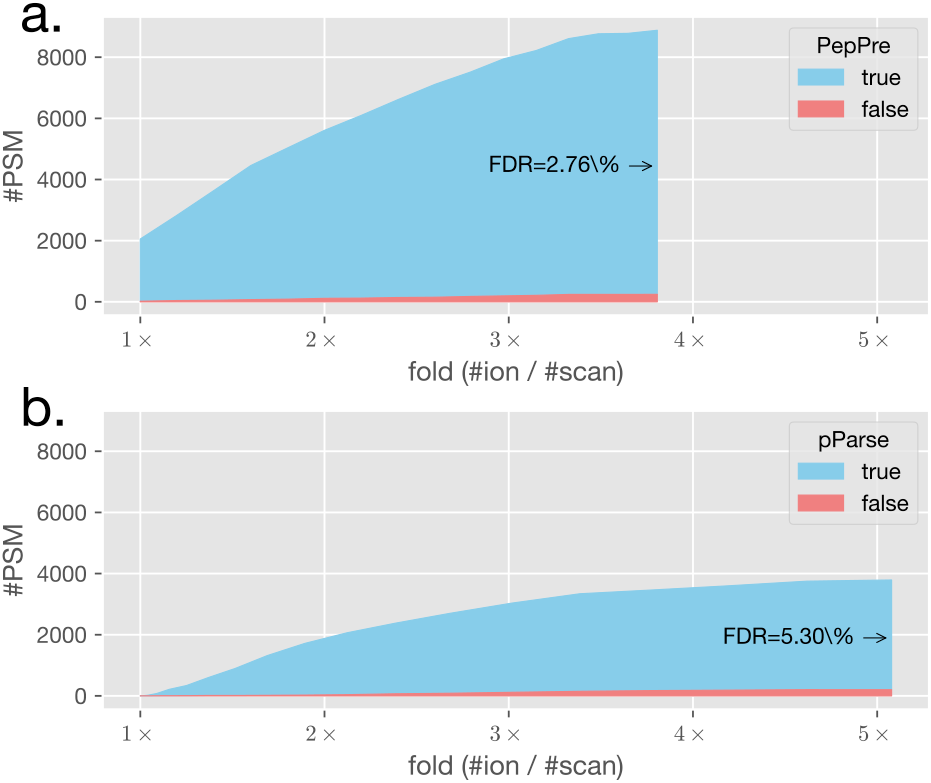
Validation on cofragmented precursors that share peaks with any others using Dong-DSS-1.6. The results show that PepPre is better at detecting more complex and heavily mixed clusters, and the clusters reported are also more credible than pParse.

### Isolation Window Size and Time Costs

In this part of the experiments, we evaluate the performance of PepPre and pParse on datasets with various isolation window sizes. The Zubarev-Human-2/4/6/8 datasets are applied, and their isolation window sizes are 2 Th, 4 Th, 6 Th, and 8 Th respectively. We also try to set methods to different window sizes. Figure 7 reports the numbers of identified PSMs when the fold is set to 4. The results show that both pParse and PepPre achieve the best performance when the window size is set to the same as the instrument. For each setting, the number of identified PSMs of PepPre is always significantly more than pParse, and the increments vary from about 4% to 10%.

**Figure 7:**
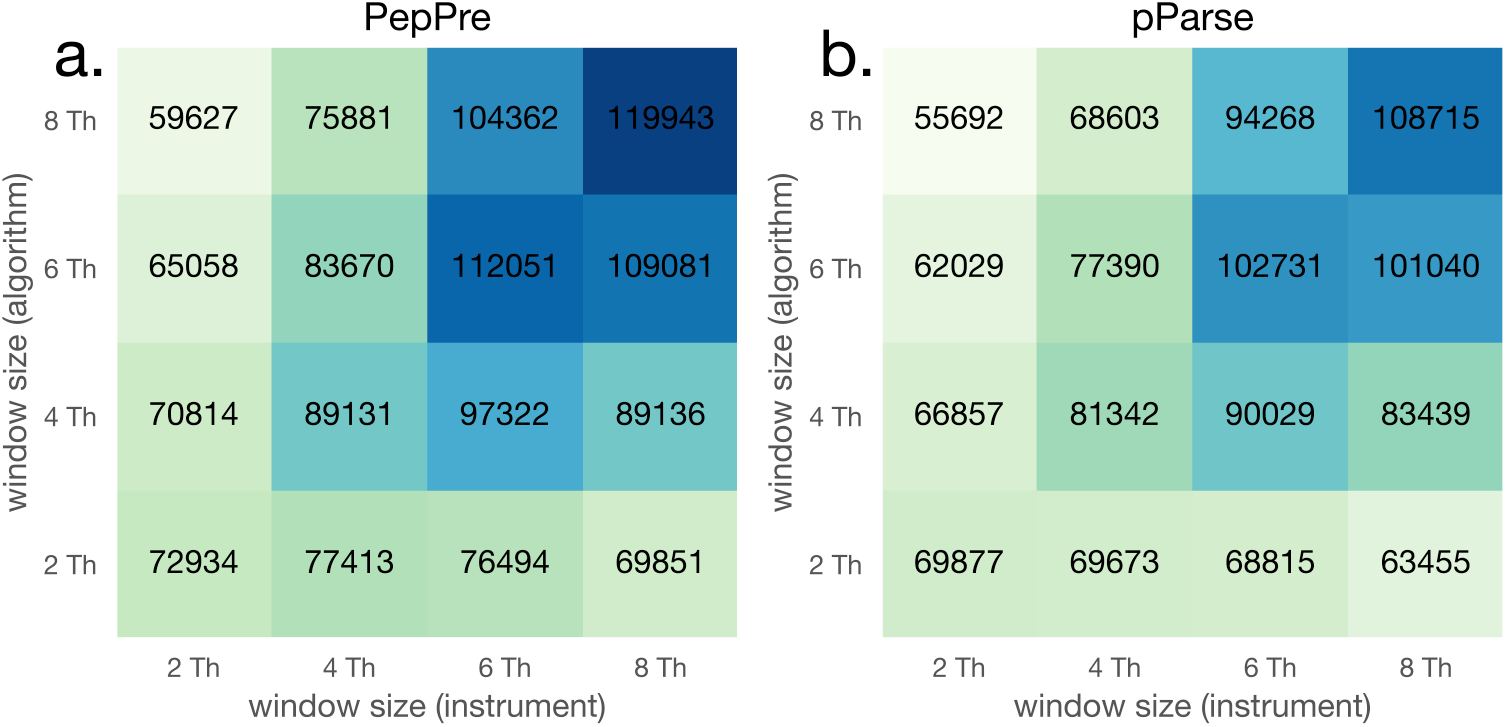
The number of identified PSMs on the datasets with various isolation window sizes, Zubarev-Human-2/4/6/8. The results show that PepPre achieves more identifications than pParse regardless of the size of the isolation window. It is also indicated that widening the isolation window size is a cheap and very effective solution to achieve more identifications.

The number of MS1 and MS2 scans is nearly equal for each run. However, e.g., with the isolation window size expanded from 2 Th to 8 Th, for PepPre, the number of identified PSMs increases from 72934 to 119943 by more than 64%, and for pParse, it increases from 69877 to 108715 by about 56%. The time costs are increased nearly linearly, as shown in Figure S1.

Conversely, increased window size does make the MS2 scans more complex. When the window size of algorithms is set to 2 Th, and as the window size of the instrument increases from 2 Th to 8 Th, the number of identified PSMs decreases by about 9% for pParse and by 4% for PepPre, which also indicates that PepPre is more robust than pParse.

For the dataset with 8 Th isolation window, Figure 8 illustrates a case, where the single one MS2 identifies 9 peptides, which shows the potential of larger isolation windows. More cases and figures of each PSM are included in Figure S15 S53.

**Figure 8:**
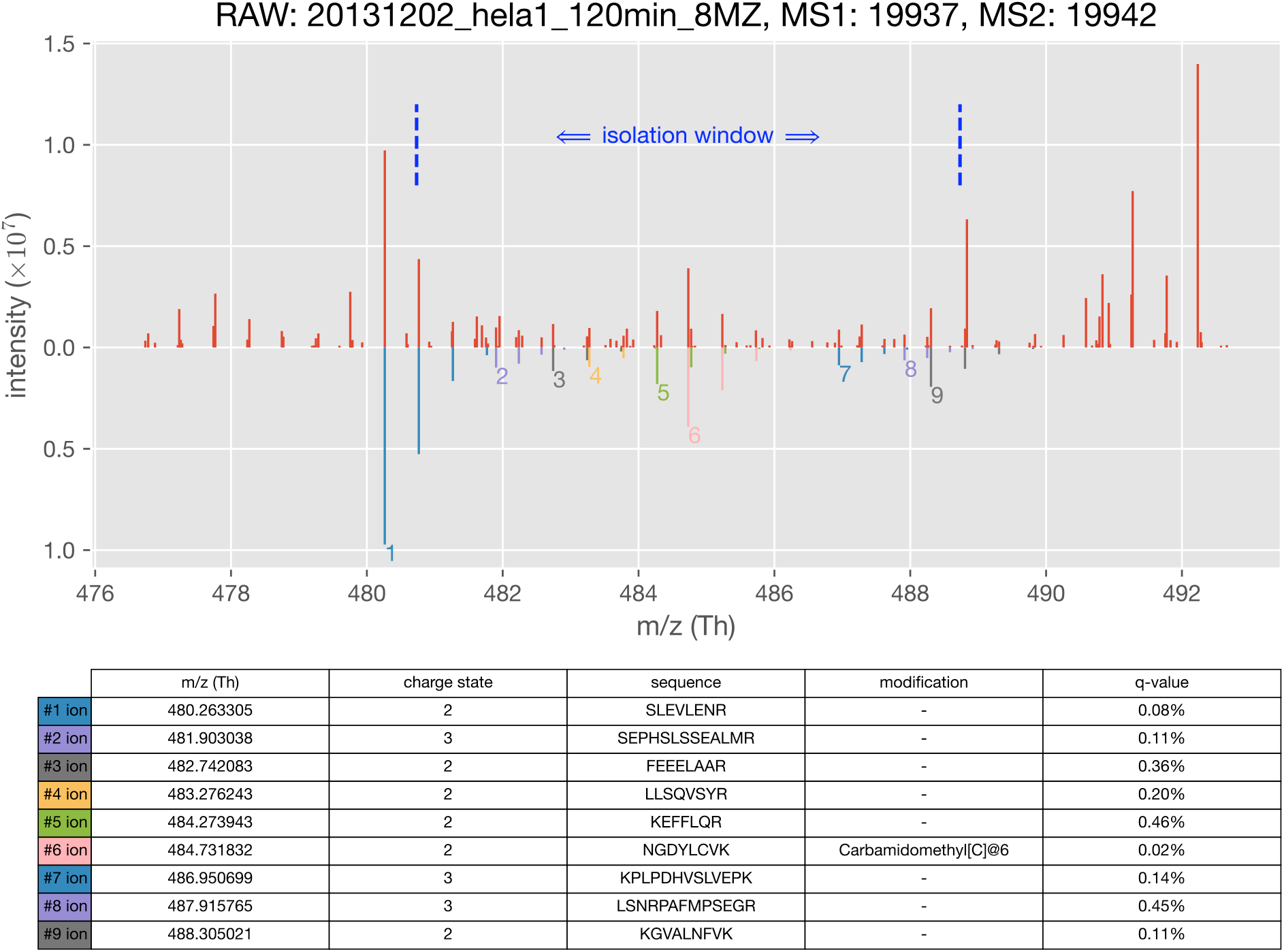
A case that 9 peptides fragmented into a single MS2 scan are successfully identified. The upper part is experimental peaks, and the lower part is identified precursor ions estimated by PepPre.

## Discussion

In this work, we proposed a simple and effective method to detect the peptide precursor ions of MS2 scans. This approach is easy to understand and achieves significant performance improvement. It is also portable and lightweight, and thus easier to use or implement. And importantly, no training is required by PepPre, which frees developers or users from labelling data and also makes it fundamentally more robust. Besides peptide precursor ions detection, this approach can also be extended to many fields, such as peptide feature detection, MS2 deisotoping, and top-down proteomics. We look forward that the method can improve data analyses of MS-based proteomics and related fields, and the software can serve the community in the long term.

## Supporting information

Graphical User Interface of PepPre

Graphical User Interface of PepPreView

## Acknowledgement

We thank the support from other members of the pFind team and especially the invaluable review and discussion from Prof. Si-Min He. We also thank Prof. Meng-Qiu Dong, Dr. Yong Cao, and other members of Dong’s lab for providing the dataset and discussion. Particularly, C. T. thanks his beautiful BB for her love.

## Supporting Information Available

- Supporting Tables and Figures
- Software and Source Code: http://peppre.ctarn.io
- Averagine Model Visualization: http://averagine.ctarn.io
- Graphical User Interface of PepPre: PepPre GUI.pdf
- Graphical User Interface of PepPreView: PepPreView GUI.pdf

## Notes

http://peppre.ctarn.io

### Competing Interest Statement

The authors have declared no competing interest.

