## Supporting Tables and Figures for "PepPre: Promote Peptide Identification Using Accurate and Comprehensive Precursors"

### Contents

Time Costs S4

PepPre+ S5

Identification Including Protein Group Level S6

PepPre vs. pParse S7

PepPre vs. RawConverter S8

PepPre vs. Monocle S9

PepPre vs. Decon2LS S10

PepPre vs. RAPID S11

PepPre vs. MaxQuant S12

PepPre vs. Dinosaur S13

PepPre vs. PointIso S14

PepPre vs. EnumInst S15

PepPre vs. EnumIW S16

PepPre vs. EnumEx S17

Charge State Distribution S18

Mass Distribution S19

Mixture Spectrum S20

**Center Ion vs. Non-center Ion** **S21**

**Case #1** **S22**

**Case #2** **S32**

**Case #3** **S42**

**Case #4** **S52**

### Time Costs

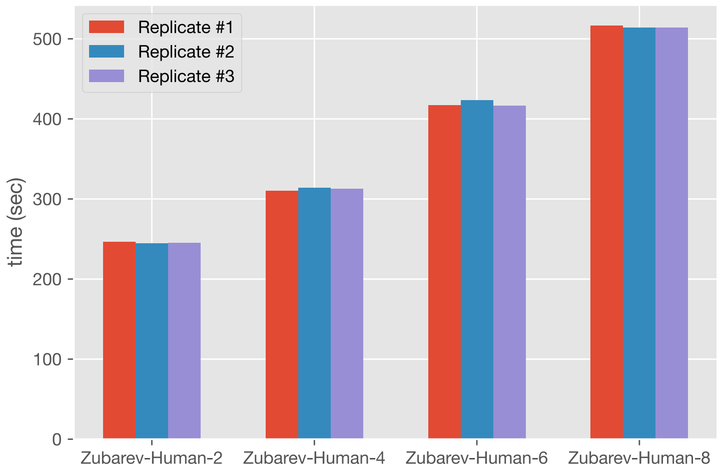


Figure S1: Time Costs. The experiments are applied on an iMac with a 3.6 GHz 8-Core Intel Core i9 processor and 64 GB 2667 MHz DDR4 memory. The software is not parallelized and only one core is used.

### PepPre+

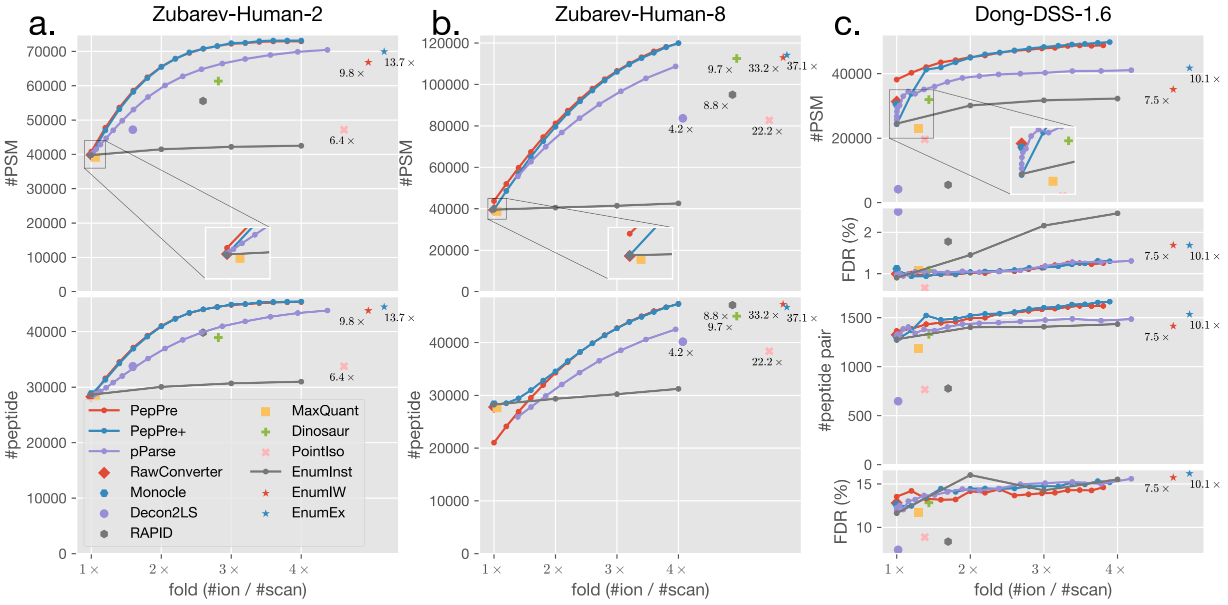


Figure S2: Identification and validation results with PepPre+ added. The figure shows that PepPre identifies more PSMs but fewer peptides compared with PepPre+ on Zubarev-Human-8 with small folds.

### Identification Including Protein Group Level

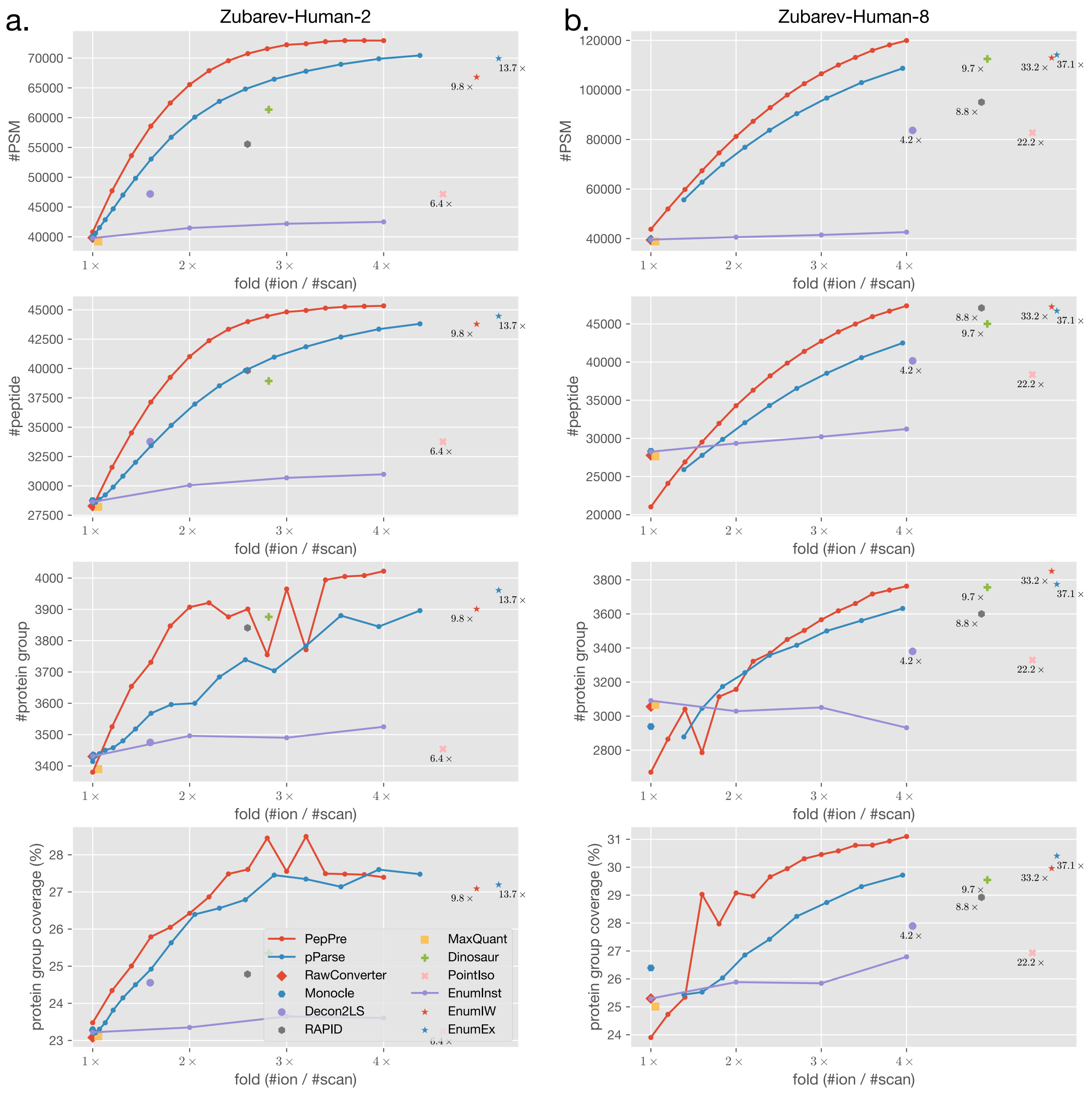


Figure S3: Identification results including the number and sequence coverage of protein groups.

### PepPre vs. pParse

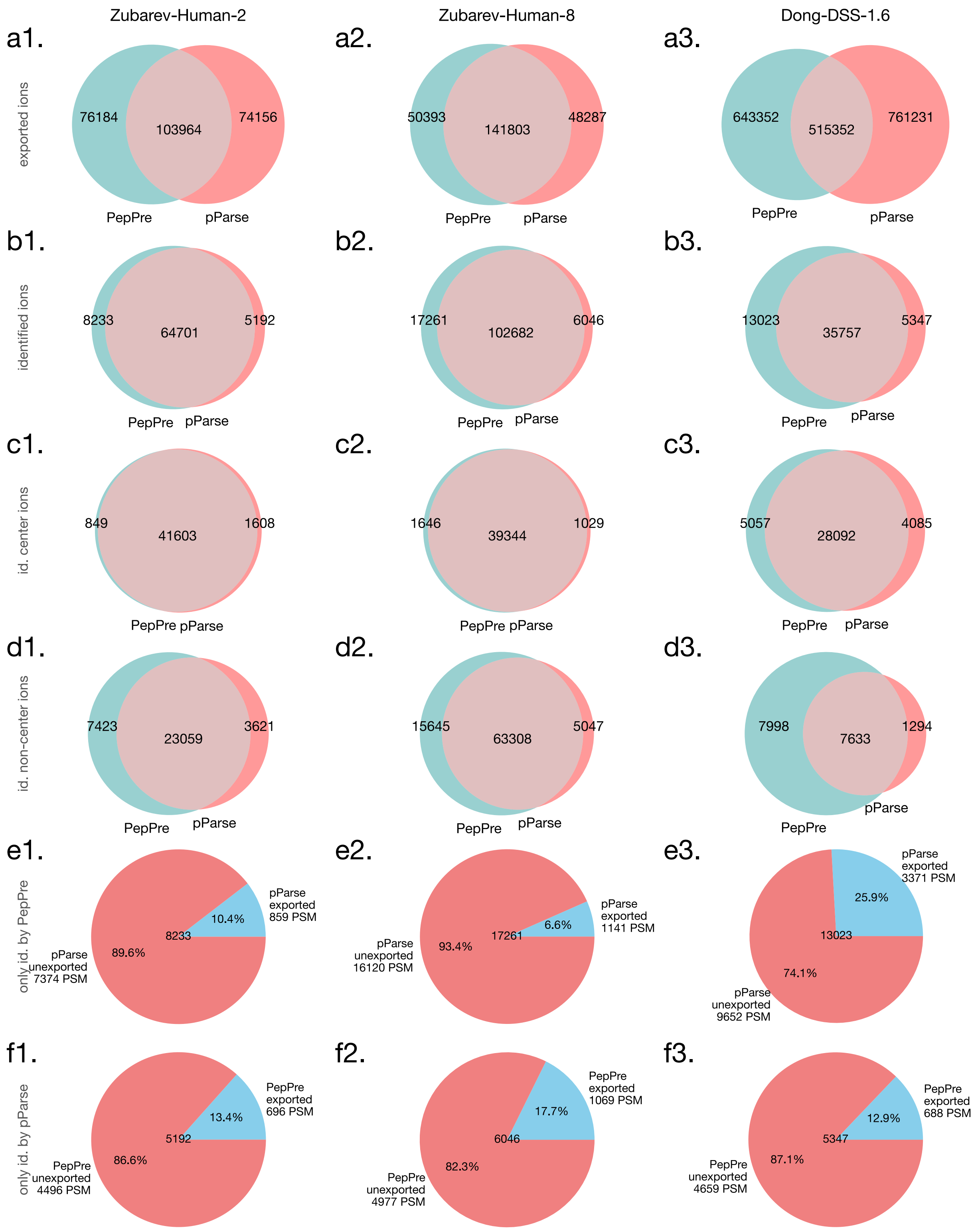


Figure S4: Comparison between PepPre and pParse.

### PepPre vs. RawConverter

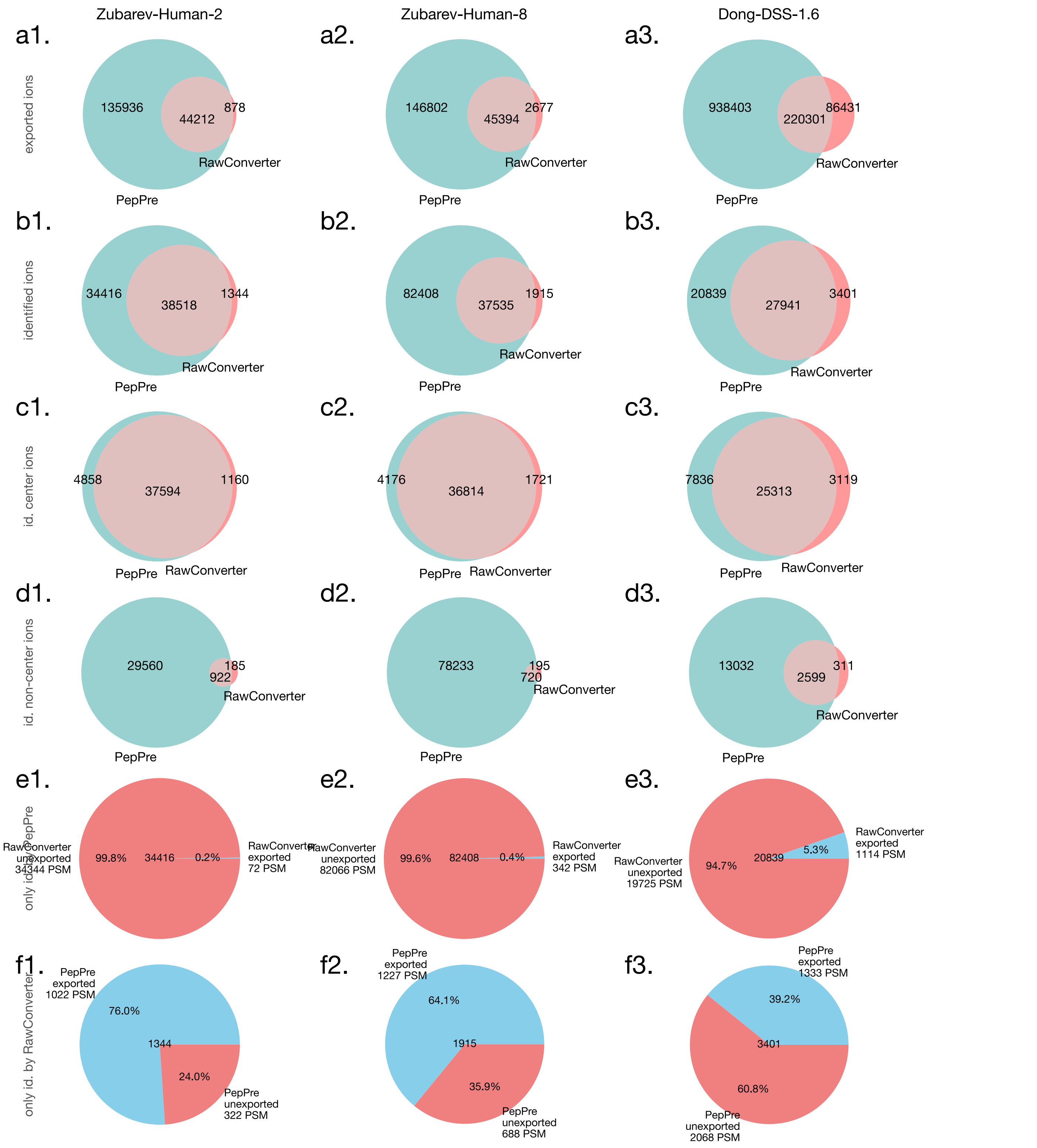


Figure S5: Comparison between PepPre and RawConverter.

### PepPre vs. Monocle

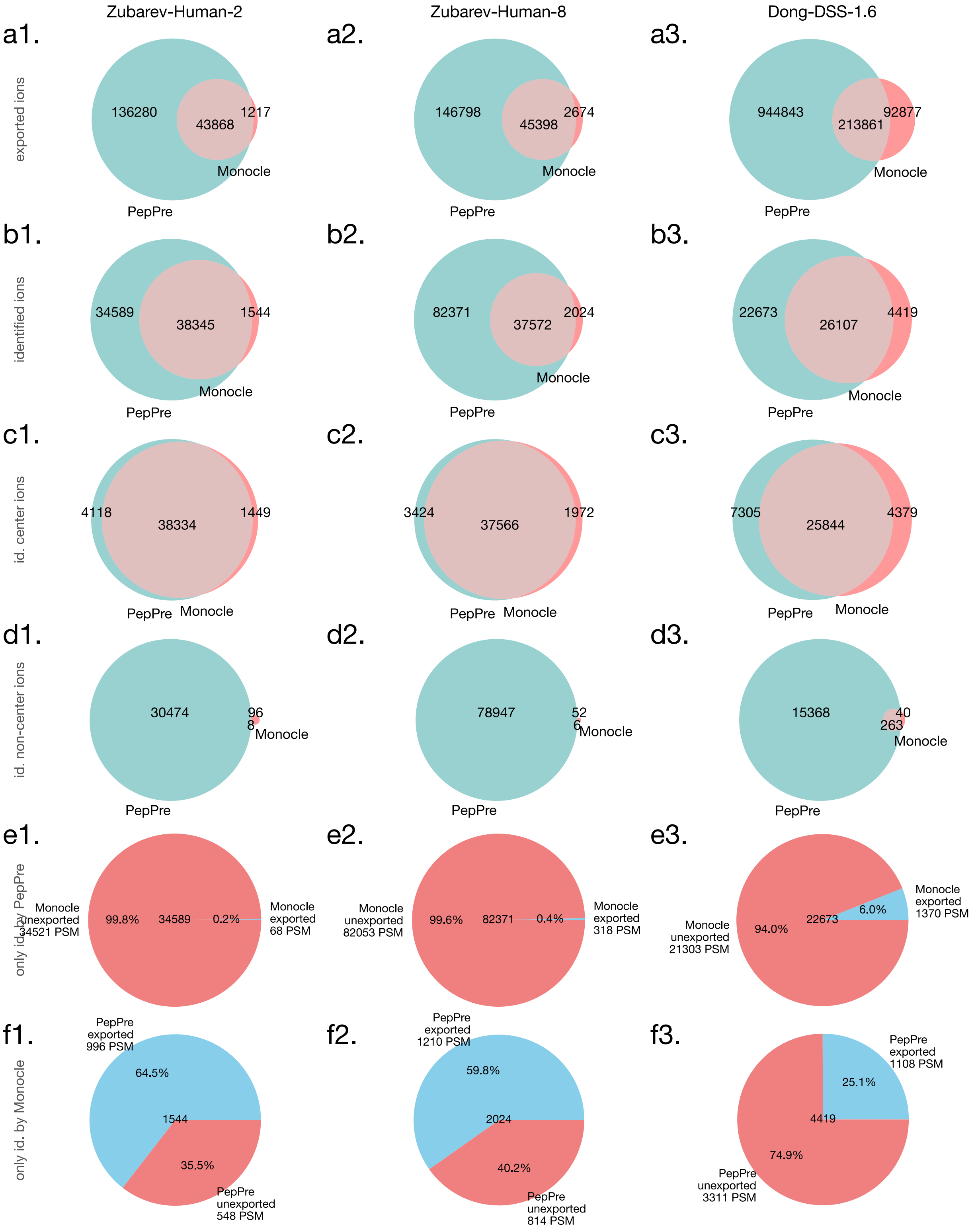


Figure S6: Comparison between PepPre and Monocle.

### PepPre vs. Decon2LS

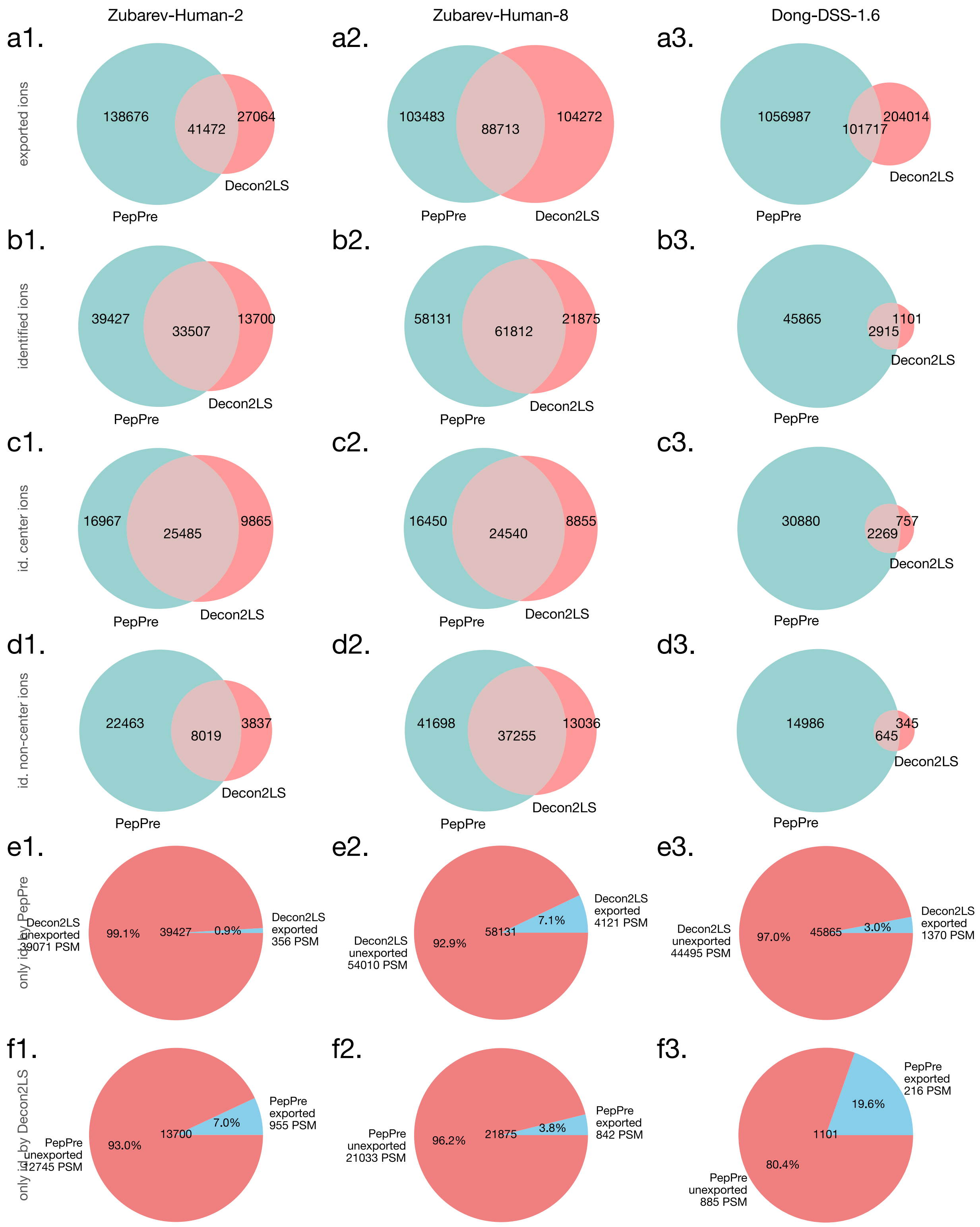


Figure S7: Comparison between PepPre and Decon2LS.

### PepPre vs. RAPID

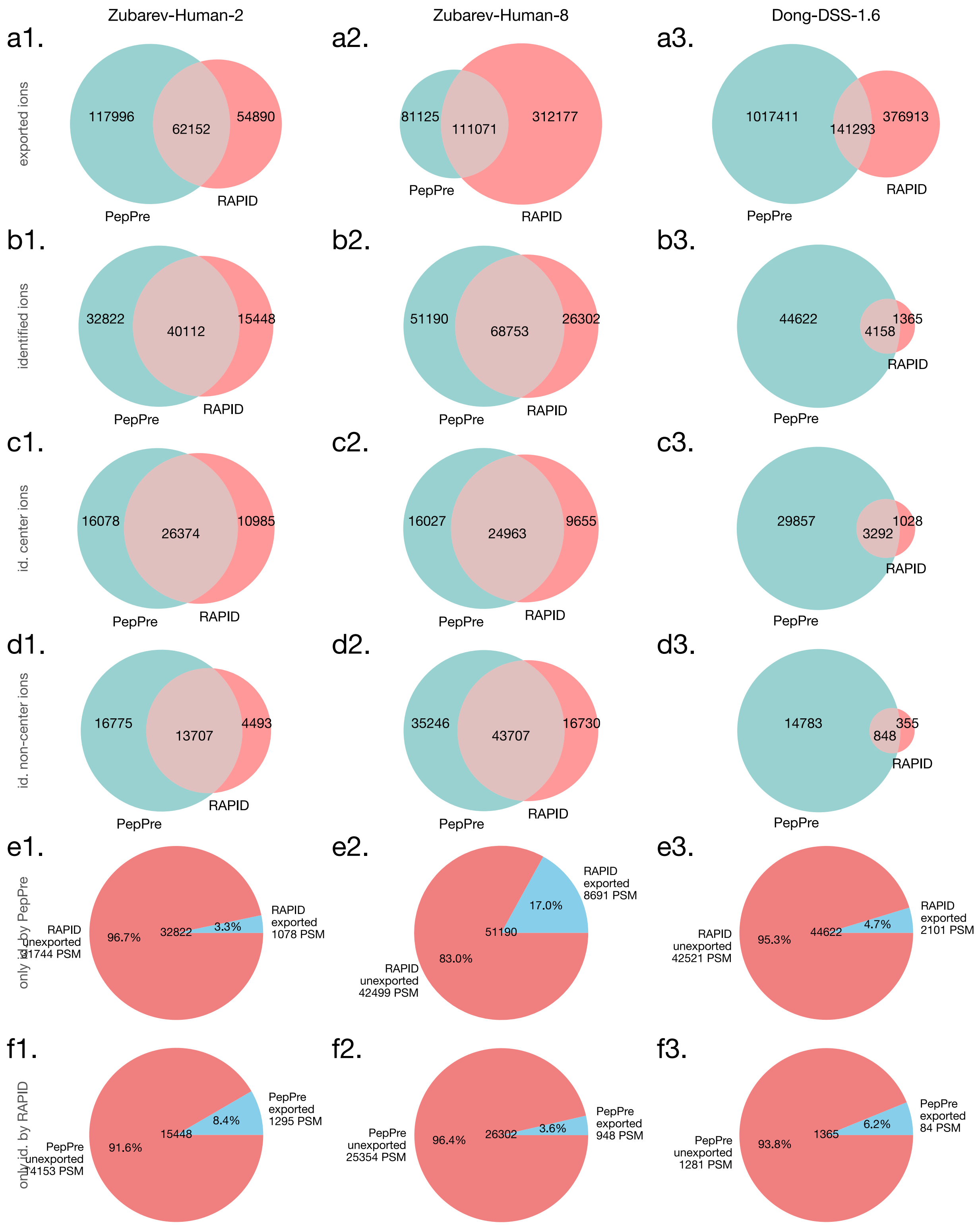


Figure S8: Comparison between PepPre and RAPID.

### PepPre vs. MaxQuant

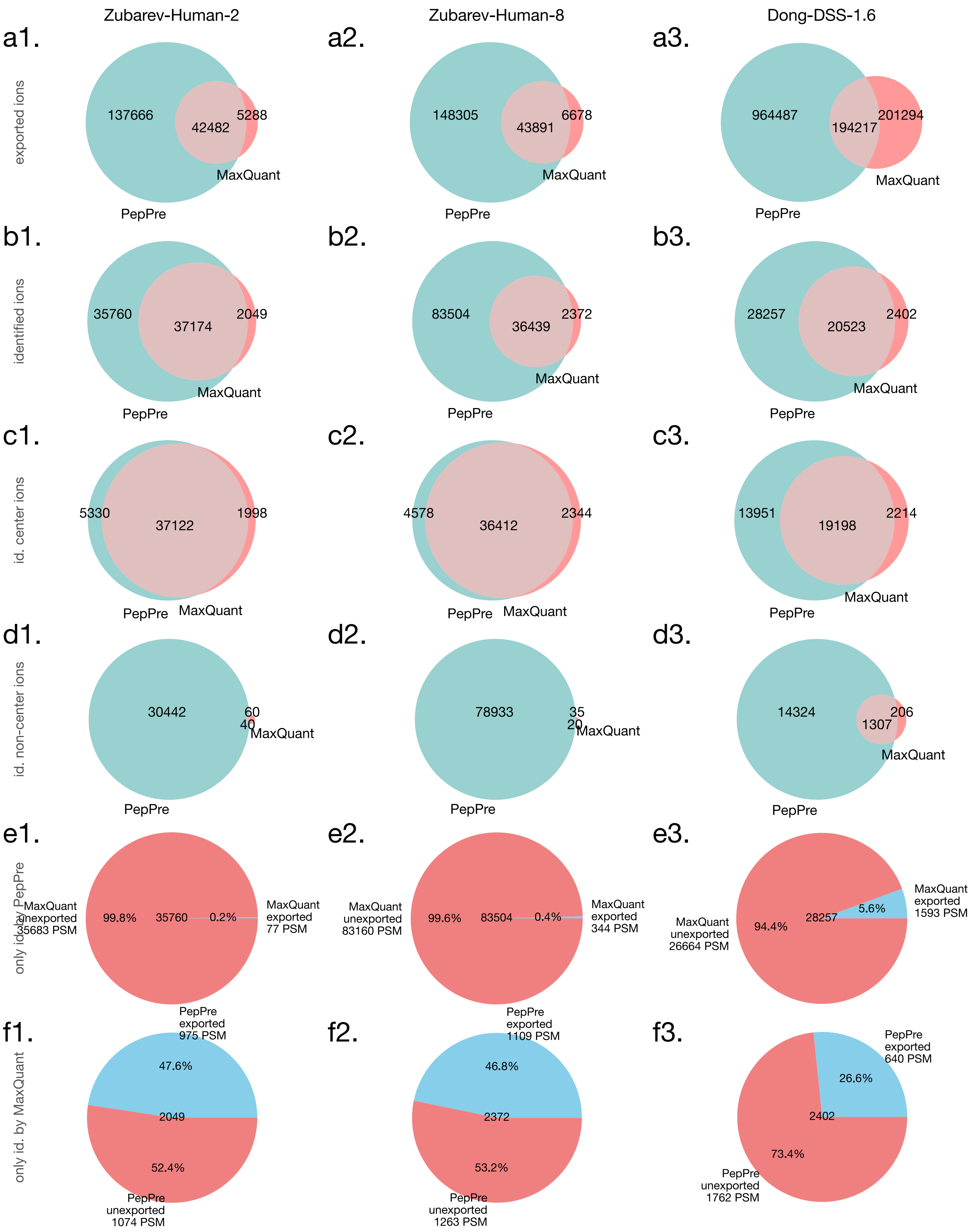


Figure S9: Comparison between PepPre and MaxQuant.

### PepPre vs. Dinosaur

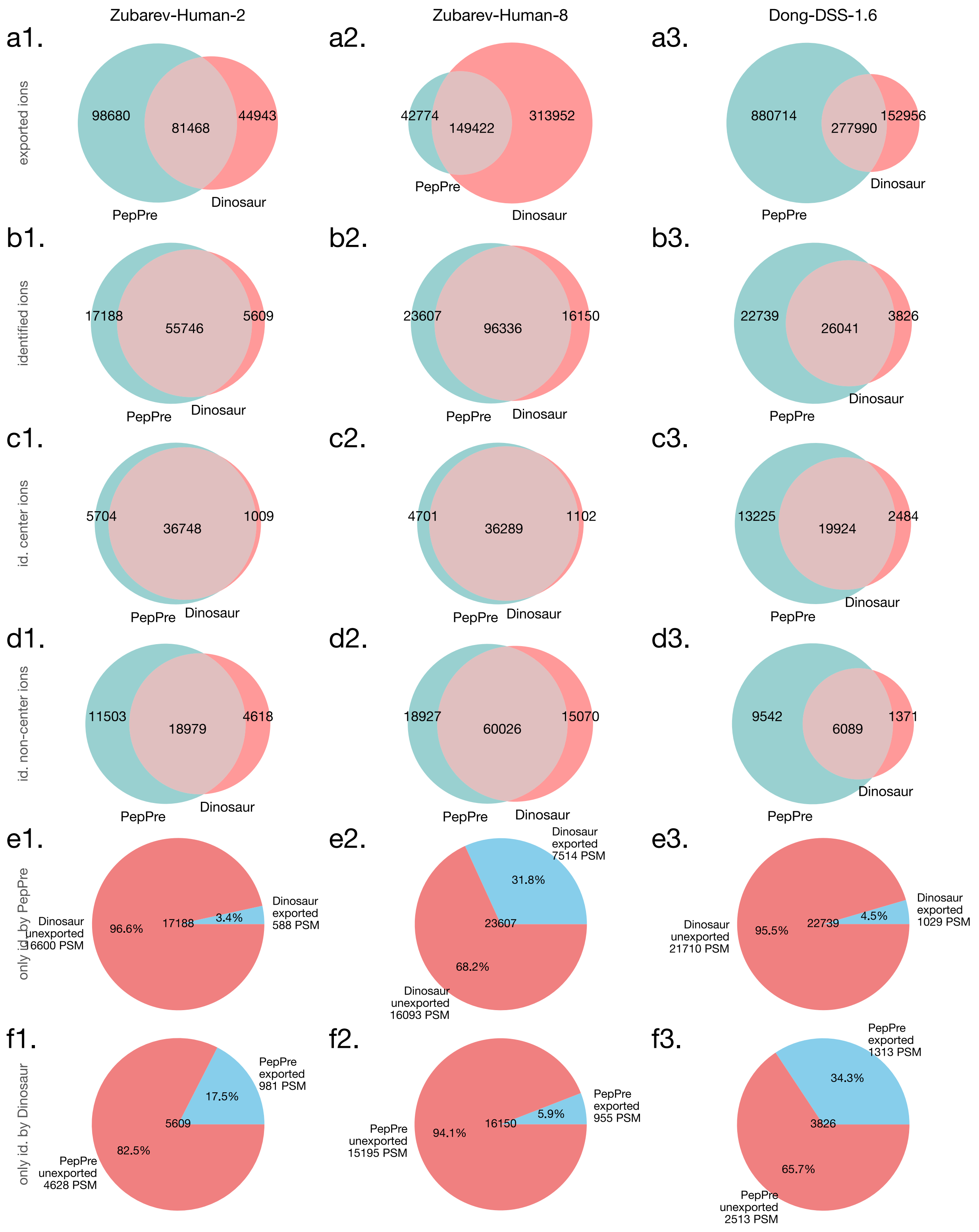


Figure S10: Comparison between PepPre and Dinosaur.

### PepPre vs. PointIso

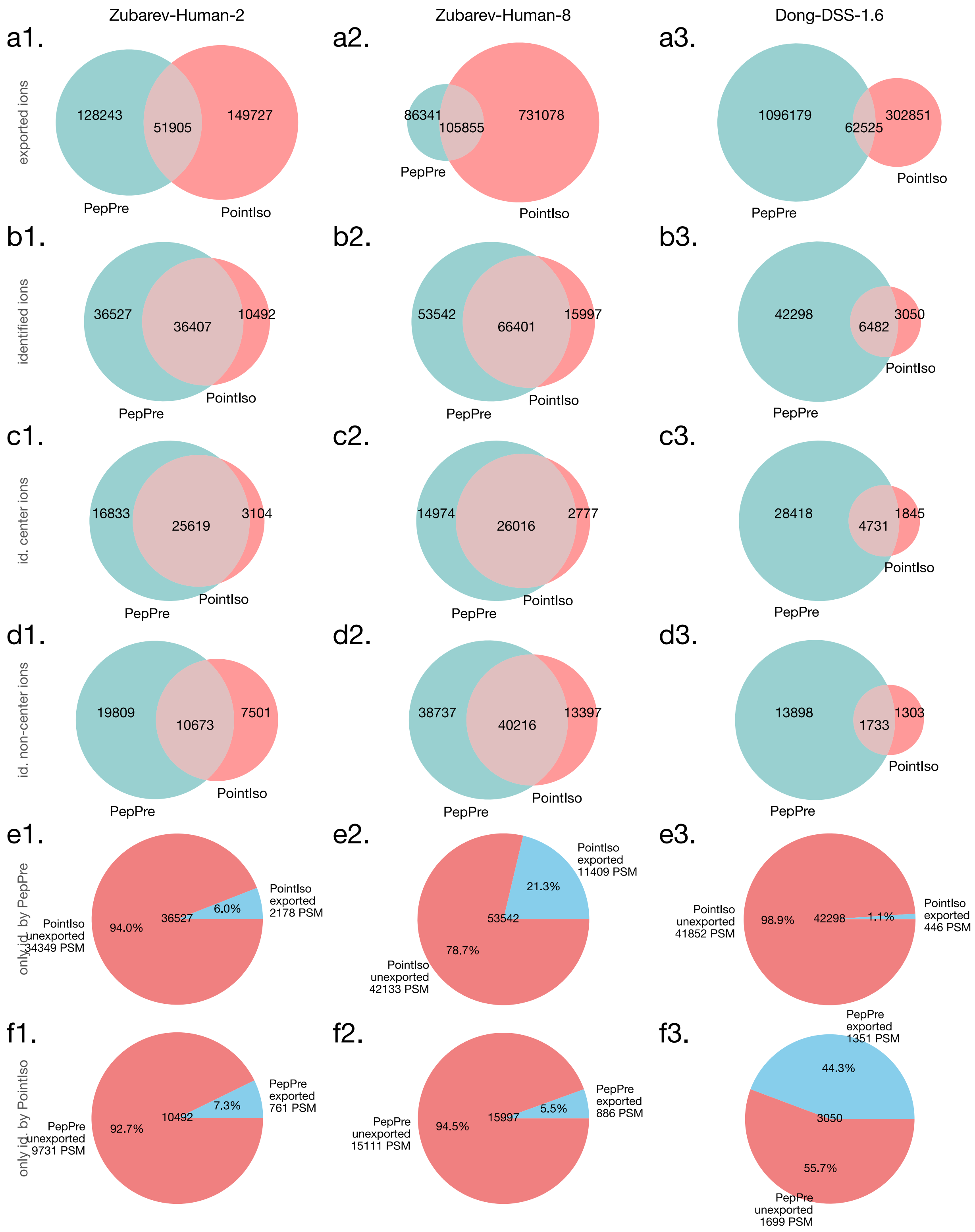


Figure S11: Comparison between PepPre and PointIso.

### PepPre vs. EnumInst

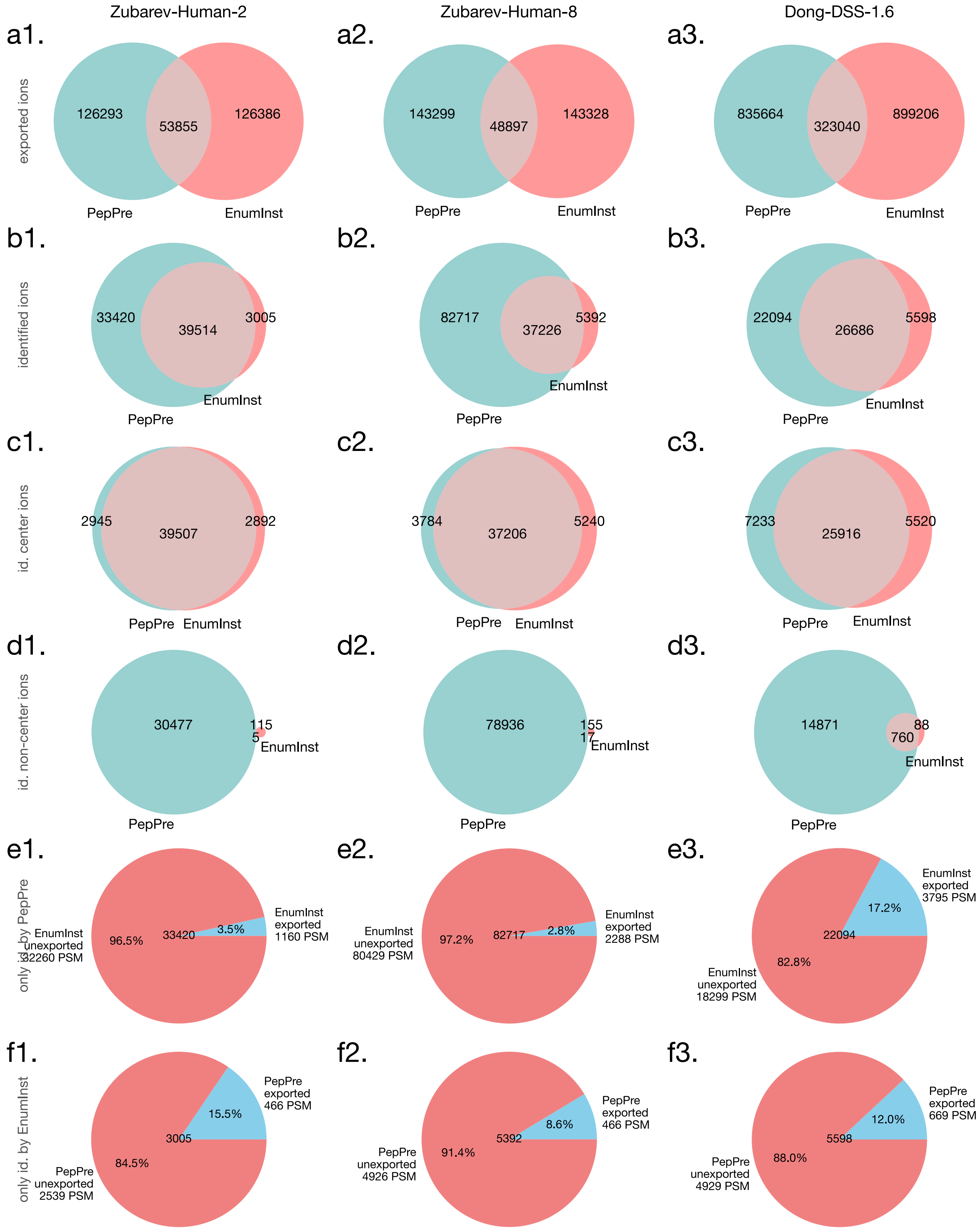


Figure S12: Comparison between PepPre and EnumInst.

### PepPre vs. EnumIW

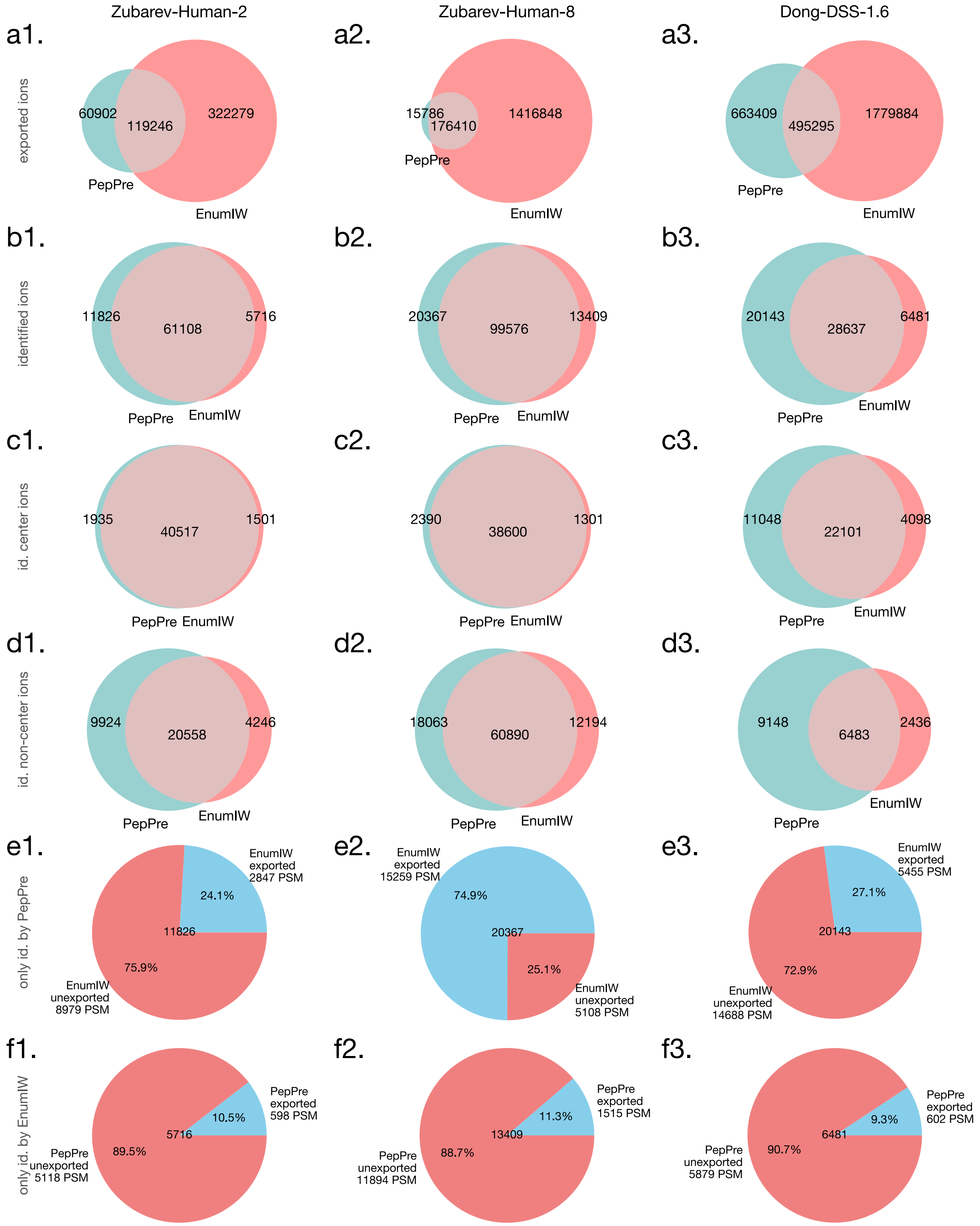


Figure S13: Comparison between PepPre and EnumIW.

### PepPre vs. EnumEx

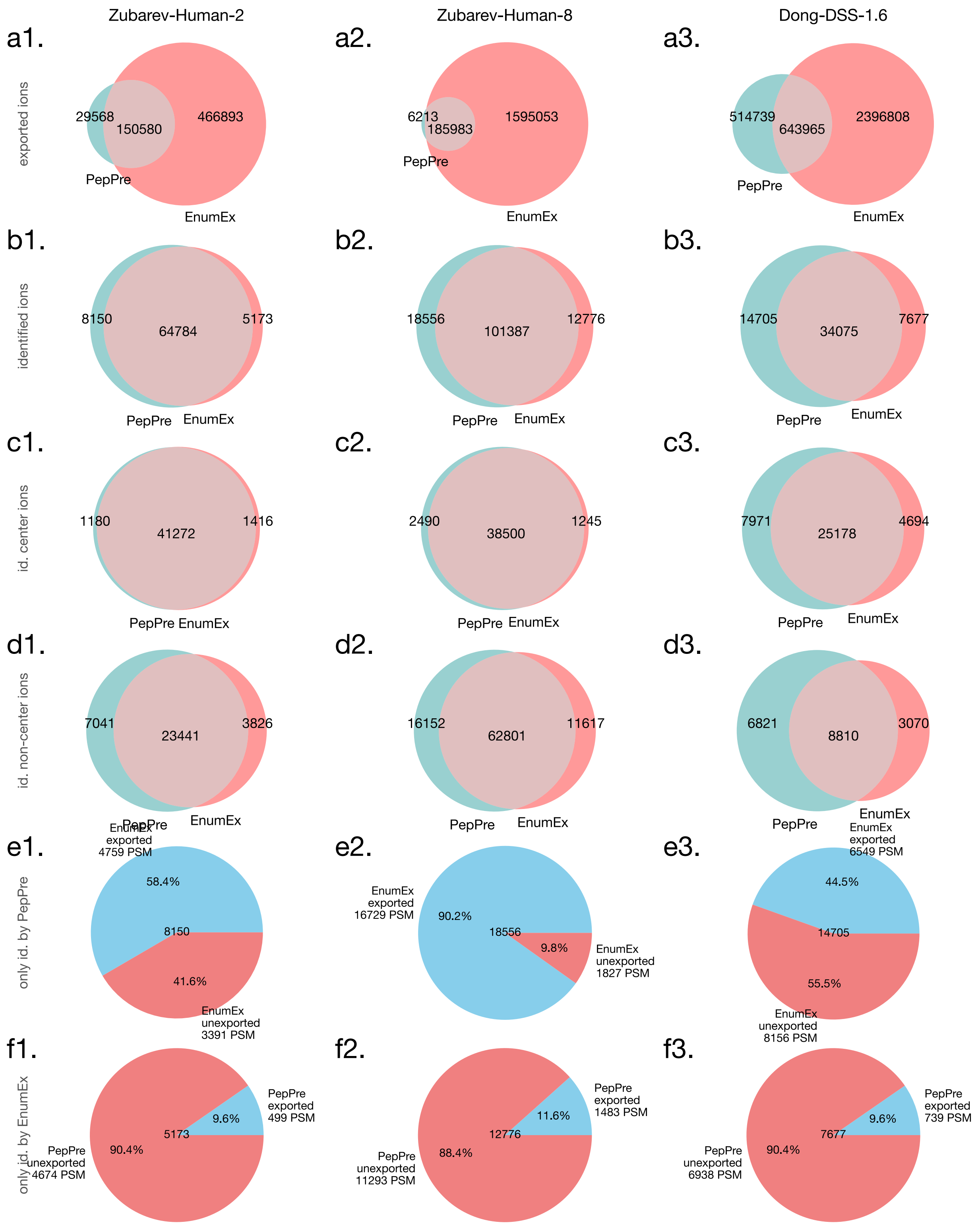


Figure S14: Comparison between PepPre and EnumEx.

### Charge State Distribution

Table S1: Zubarev-Human-2

| Charge | 1+ | 2+ | 3+ | 4+ | 5+ | $\geq$ 6+ |
|---|---|---|---|---|---|---|
| PepPre | 0 (0.00%) | 45741 (62.72%) | 23418 (32.11%) | 3259 (4.47%) | 468 (0.64%) | 48 (0.07%) |
| pParse | 0 (0.00%) | 44703 (63.97%) | 21768 (31.15%) | 2944 (4.21%) | 423 (0.61%) | 39 (0.06%) |
| RawConverter | 0 (0.00%) | 24088 (60.44%) | 13430 (33.70%) | 2009 (5.04%) | 301 (0.76%) | 28 (0.07%) |
| Monocle | 0 (0.00%) | 24184 (60.64%) | 13367 (33.52%) | 2002 (5.02%) | 300 (0.75%) | 30 (0.08%) |
| Decon2LS | 182 (0.39%) | 29411 (62.32%) | 15203 (32.21%) | 2067 (4.38%) | 306 (0.65%) | 27 (0.06%) |
| RAPID | 315 (0.57%) | 34990 (62.99%) | 17424 (31.37%) | 2386 (4.30%) | 369 (0.66%) | 64 (0.12%) |
| MaxQuant | 31 (0.08%) | 23772 (60.62%) | 13191 (33.64%) | 1913 (4.88%) | 265 (0.68%) | 43 (0.11%) |
| Dinosaur | 399 (0.65%) | 37346 (60.87%) | 20348 (33.16%) | 2825 (4.60%) | 400 (0.65%) | 36 (0.06%) |
| PointIso | 16 (0.03%) | 29727 (63.01%) | 17030 (36.10%) | 390 (0.83%) | 12 (0.03%) | 0 (0.00%) |
| EnumInst | 0 (0.00%) | 25956 (61.06%) | 14142 (33.27%) | 2073 (4.88%) | 307 (0.72%) | 32 (0.08%) |
| EnumIW | 0 (0.00%) | 41707 (62.43%) | 21465 (32.13%) | 3107 (4.65%) | 446 (0.67%) | 82 (0.12%) |
| EnumEx | 0 (0.00%) | 44115 (63.08%) | 22175 (31.71%) | 3128 (4.47%) | 445 (0.64%) | 76 (0.11%) |

Table S2: Zubarev-Human-8

| Charge | 1+ | 2+ | 3+ | 4+ | 5+ | $\geq$ 6+ |
|---|---|---|---|---|---|---|
| PepPre | 0 (0.00%) | 74938 (62.48%) | 38876 (32.41%) | 5369 (4.48%) | 706 (0.59%) | 54 (0.05%) |
| pParse | 0 (0.00%) | 67632 (62.21%) | 35629 (32.77%) | 4798 (4.41%) | 608 (0.56%) | 48 (0.04%) |
| RawConverter | 0 (0.00%) | 23846 (60.45%) | 13300 (33.72%) | 2015 (5.11%) | 270 (0.68%) | 17 (0.04%) |
| Monocle | 0 (0.00%) | 24097 (60.86%) | 13203 (33.35%) | 2006 (5.07%) | 265 (0.67%) | 24 (0.06%) |
| Decon2LS | 689 (0.82%) | 53326 (63.73%) | 25925 (30.98%) | 3261 (3.90%) | 436 (0.52%) | 39 (0.05%) |
| RAPID | 1228 (1.29%) | 61756 (64.98%) | 27931 (29.39%) | 3534 (3.72%) | 486 (0.51%) | 110 (0.12%) |
| MaxQuant | 17 (0.04%) | 23607 (60.83%) | 12994 (33.48%) | 1918 (4.94%) | 244 (0.63%) | 30 (0.08%) |
| Dinosaur | 1433 (1.27%) | 69192 (61.50%) | 36147 (32.13%) | 5073 (4.51%) | 600 (0.53%) | 65 (0.06%) |
| PointIso | 131 (0.16%) | 52531 (63.51%) | 29185 (35.29%) | 840 (1.02%) | 24 (0.03%) | 0 (0.00%) |
| EnumInst | 0 (0.00%) | 25169 (59.06%) | 14902 (34.97%) | 2221 (5.21%) | 302 (0.71%) | 23 (0.05%) |
| EnumIW | 0 (0.00%) | 71824 (63.57%) | 35409 (31.34%) | 4951 (4.38%) | 682 (0.60%) | 110 (0.10%) |
| EnumEx | 0 (0.00%) | 72743 (63.72%) | 35637 (31.22%) | 4982 (4.36%) | 682 (0.60%) | 110 (0.10%) |

Table S3: Dong-DSS-1.6

| Charge | 1+ | 2+ | 3+ | 4+ | 5+ | $\geq$ 6+ |
|---|---|---|---|---|---|---|
| PepPre | 0 (0.00%) | 93 (0.19%) | 22287 (45.69%) | 22085 (45.27%) | 3999 (8.20%) | 316 (0.65%) |
| pParse | 0 (0.00%) | 90 (0.22%) | 19463 (47.37%) | 18365 (44.70%) | 2907 (7.08%) | 261 (0.64%) |
| RawConverter | 0 (0.00%) | 0 (0.00%) | 15094 (48.16%) | 13953 (44.52%) | 2133 (6.81%) | 159 (0.51%) |
| Monocle | 0 (0.00%) | 0 (0.00%) | 15363 (50.34%) | 12887 (42.23%) | 2071 (6.79%) | 196 (0.64%) |
| Decon2LS | 0 (0.00%) | 1 (0.02%) | 2370 (56.92%) | 1604 (38.52%) | 183 (4.39%) | 6 (0.14%) |
| RAPID | 0 (0.00%) | 2 (0.04%) | 2973 (53.84%) | 2165 (39.21%) | 346 (6.27%) | 36 (0.65%) |
| MaxQuant | 0 (0.00%) | 52 (0.23%) | 11639 (50.78%) | 9647 (42.09%) | 1433 (6.25%) | 148 (0.65%) |
| Dinosaur | 0 (0.00%) | 73 (0.23%) | 14024 (43.91%) | 15286 (47.86%) | 2331 (7.30%) | 223 (0.70%) |
| PointIso | 0 (0.00%) | 82 (0.42%) | 17103 (87.09%) | 2454 (12.50%) | 0 (0.00%) | 0 (0.00%) |
| EnumInst | 0 (0.00%) | 0 (0.00%) | 16825 (52.13%) | 13331 (41.31%) | 1926 (5.97%) | 191 (0.59%) |
| EnumIW | 0 (0.00%) | 79 (0.22%) | 16410 (46.73%) | 16005 (45.58%) | 2375 (6.76%) | 248 (0.71%) |
| EnumEx | 0 (0.00%) | 95 (0.23%) | 19105 (45.76%) | 19076 (45.69%) | 3138 (7.52%) | 339 (0.81%) |

### Mass Distribution

Table S4: Zubarev-Human-2

| Mass (Da) | [0, 1000) | [1000, 1500) | [1500, 2000) | [2000, 2500) | [2500, 3000) | [3000, +∞) |
|---|---|---|---|---|---|---|
| PepPre | 14812 (20.31%) | 27684 (37.96%) | 16780 (23.01%) | 7751 (10.63%) | 3543 (4.86%) | 2364 (3.24%) |
| pParse | 14277 (20.43%) | 26820 (38.38%) | 15913 (22.77%) | 7283 (10.42%) | 3325 (4.76%) | 2259 (3.23%) |
| RawConverter | 6868 (17.23%) | 13651 (34.25%) | 9767 (24.51%) | 5100 (12.80%) | 2556 (6.41%) | 1914 (4.80%) |
| Monocle | 6986 (17.52%) | 13651 (34.23%) | 9722 (24.38%) | 5076 (12.73%) | 2541 (6.37%) | 1907 (4.78%) |
| Decon2LS | 9538 (20.21%) | 17438 (36.95%) | 10654 (22.57%) | 5055 (10.71%) | 2618 (5.55%) | 1893 (4.01%) |
| RAPID | 11936 (21.49%) | 20768 (37.39%) | 12086 (21.76%) | 5794 (10.43%) | 2883 (5.19%) | 2081 (3.75%) |
| MaxQuant | 6804 (17.35%) | 13507 (34.44%) | 9585 (24.44%) | 5004 (12.76%) | 2496 (6.36%) | 1819 (4.64%) |
| Dinosaur | 12034 (19.61%) | 22949 (37.40%) | 14462 (23.57%) | 6789 (11.07%) | 3046 (4.96%) | 2074 (3.38%) |
| PointIso | 9015 (19.11%) | 18866 (39.99%) | 11418 (24.20%) | 5285 (11.20%) | 1784 (3.78%) | 807 (1.71%) |
| EnumInst | 7384 (17.37%) | 14911 (35.08%) | 10274 (24.17%) | 5290 (12.44%) | 2678 (6.30%) | 1973 (4.64%) |
| EnumIW | 13457 (20.14%) | 25166 (37.67%) | 15370 (23.01%) | 7147 (10.70%) | 3331 (4.99%) | 2336 (3.50%) |
| EnumEx | 14144 (20.22%) | 26576 (38.00%) | 16051 (22.95%) | 7392 (10.57%) | 3425 (4.90%) | 2351 (3.36%) |

Table S5: Zubarev-Human-8

| Mass (Da) | [0, 1000) | [1000, 1500) | [1500, 2000) | [2000, 2500) | [2500, 3000) | [3000, +∞) |
|---|---|---|---|---|---|---|
| PepPre | 23065 (19.23%) | 45204 (37.69%) | 28549 (23.80%) | 12996 (10.84%) | 6163 (5.14%) | 3966 (3.31%) |
| pParse | 19509 (17.95%) | 41376 (38.06%) | 26640 (24.50%) | 12007 (11.04%) | 5614 (5.16%) | 3569 (3.28%) |
| RawConverter | 6434 (16.31%) | 13279 (33.66%) | 10065 (25.51%) | 5148 (13.05%) | 2595 (6.58%) | 1927 (4.88%) |
| Monocle | 6619 (16.72%) | 13371 (33.77%) | 10002 (25.26%) | 5104 (12.89%) | 2577 (6.51%) | 1922 (4.85%) |
| Decon2LS | 17114 (20.45%) | 33409 (39.93%) | 18584 (22.21%) | 7735 (9.24%) | 4063 (4.86%) | 2771 (3.31%) |
| RAPID | 19819 (20.85%) | 38359 (40.36%) | 20777 (21.86%) | 8688 (9.14%) | 4392 (4.62%) | 3010 (3.17%) |
| MaxQuant | 6407 (16.51%) | 13179 (33.96%) | 9852 (25.39%) | 5002 (12.89%) | 2517 (6.49%) | 1853 (4.77%) |
| Dinosaur | 20888 (18.57%) | 42863 (38.10%) | 27380 (24.34%) | 12335 (10.96%) | 5488 (4.88%) | 3556 (3.16%) |
| PointIso | 13965 (16.88%) | 34114 (41.24%) | 20855 (25.21%) | 9306 (11.25%) | 3182 (3.85%) | 1289 (1.56%) |
| EnumInst | 6054 (14.21%) | 14362 (33.70%) | 11234 (26.36%) | 5832 (13.68%) | 2993 (7.02%) | 2142 (5.03%) |
| EnumIW | 20792 (18.40%) | 43788 (38.76%) | 27065 (23.96%) | 11995 (10.62%) | 5641 (4.99%) | 3695 (3.27%) |
| EnumEx | 21087 (18.47%) | 44202 (38.72%) | 27319 (23.93%) | 12085 (10.59%) | 5736 (5.02%) | 3725 (3.26%) |

Table S6: Dong-DSS-1.6

| Mass (Da) | [0, 1000) | [1000, 1500) | [1500, 2000) | [2000, 2500) | [2500, 3000) | [3000, +∞) |
|---|---|---|---|---|---|---|
| PepPre | 0 (0.00%) | 1 (0.00%) | 6504 (13.33%) | 16485 (33.79%) | 19292 (39.55%) | 6498 (13.32%) |
| pParse | 0 (0.00%) | 1 (0.00%) | 5479 (13.34%) | 14104 (34.33%) | 16430 (39.99%) | 5072 (12.34%) |
| RawConverter | 0 (0.00%) | 1 (0.00%) | 4258 (13.59%) | 11338 (36.18%) | 12046 (38.44%) | 3696 (11.79%) |
| Monocle | 0 (0.00%) | 1 (0.00%) | 4114 (13.48%) | 10150 (33.26%) | 12213 (40.02%) | 4039 (13.24%) |
| Decon2LS | 0 (0.00%) | 0 (0.00%) | 1473 (35.37%) | 1512 (36.31%) | 1114 (26.75%) | 65 (1.56%) |
| RAPID | 0 (0.00%) | 0 (0.00%) | 1605 (29.07%) | 2058 (37.27%) | 1566 (28.36%) | 293 (5.31%) |
| MaxQuant | 0 (0.00%) | 0 (0.00%) | 3236 (14.12%) | 7969 (34.77%) | 8908 (38.87%) | 2806 (12.24%) |
| Dinosaur | 0 (0.00%) | 0 (0.00%) | 4492 (14.07%) | 11173 (34.98%) | 12338 (38.63%) | 3934 (12.32%) |
| PointIso | 0 (0.00%) | 0 (0.00%) | 6840 (34.83%) | 7477 (38.07%) | 4819 (24.54%) | 503 (2.56%) |
| EnumInst | 0 (0.00%) | 1 (0.00%) | 4450 (13.79%) | 10875 (33.70%) | 12799 (39.66%) | 4148 (12.85%) |
| EnumIW | 0 (0.00%) | 1 (0.00%) | 5191 (14.78%) | 11668 (33.23%) | 13837 (39.40%) | 4420 (12.59%) |
| EnumEx | 0 (0.00%) | 1 (0.00%) | 6006 (14.38%) | 13915 (33.33%) | 16307 (39.06%) | 5524 (13.23%) |

### Mixture Spectrum

Table S7: Zubarev-Human-2

| #PSM per MS2 | ≥ 1 | ≥ 2 | ≥ 3 | ≥ 4 | ≥ 5 | ≥ 6 |
|---|---|---|---|---|---|---|
| PepPre | 40751 (90.48%) | 21425 (47.57%) | 8060 (17.90%) | 2208 (4.90%) | 427 (0.95%) | 55 (0.12%) |
| pParse | 40831 (90.66%) | 20151 (44.74%) | 6989 (15.52%) | 1610 (3.57%) | 267 (0.59%) | 26 (0.06%) |
| RawConverter | 39856 (88.50%) | 0 (0.00%) | 0 (0.00%) | 0 (0.00%) | 0 (0.00%) | 0 (0.00%) |
| Monocle | 39883 (88.56%) | 0 (0.00%) | 0 (0.00%) | 0 (0.00%) | 0 (0.00%) | 0 (0.00%) |
| Decon2LS | 38115 (84.63%) | 8106 (18.00%) | 917 (2.04%) | 55 (0.12%) | 3 (0.01%) | 0 (0.00%) |
| RAPID | 39161 (86.95%) | 13275 (29.48%) | 2752 (6.11%) | 332 (0.74%) | 26 (0.06%) | 2 (0.00%) |
| MaxQuant | 39152 (86.93%) | 63 (0.14%) | 0 (0.00%) | 0 (0.00%) | 0 (0.00%) | 0 (0.00%) |
| Dinosaur | 39204 (87.05%) | 16557 (36.76%) | 4618 (10.25%) | 866 (1.92%) | 98 (0.22%) | 11 (0.02%) |
| PointIso | 32664 (72.53%) | 11258 (25.00%) | 2675 (5.94%) | 511 (1.13%) | 60 (0.13%) | 5 (0.01%) |
| EnumInst | 39493 (87.69%) | 2930 (6.51%) | 85 (0.19%) | 2 (0.00%) | 0 (0.00%) | 0 (0.00%) |
| EnumIW | 40654 (90.27%) | 18457 (40.98%) | 6014 (13.35%) | 1391 (3.09%) | 248 (0.55%) | 39 (0.09%) |
| EnumEx | 40671 (90.31%) | 19970 (44.34%) | 7083 (15.73%) | 1823 (4.05%) | 331 (0.73%) | 56 (0.12%) |

Table S8: Zubarev-Human-8

| #PSM per MS2 | ≥ 1 | ≥ 2 | ≥ 3 | ≥ 4 | ≥ 5 | ≥ 6 |
|---|---|---|---|---|---|---|
| PepPre | 44623 (92.87%) | 35406 (73.69%) | 22914 (47.69%) | 11452 (23.83%) | 4236 (8.82%) | 1073 (2.23%) |
| pParse | 44467 (92.55%) | 33543 (69.81%) | 19575 (40.74%) | 8197 (17.06%) | 2439 (5.08%) | 437 (0.91%) |
| RawConverter | 39448 (82.10%) | 0 (0.00%) | 0 (0.00%) | 0 (0.00%) | 0 (0.00%) | 0 (0.00%) |
| Monocle | 39595 (82.41%) | 0 (0.00%) | 0 (0.00%) | 0 (0.00%) | 0 (0.00%) | 0 (0.00%) |
| Decon2LS | 42528 (88.51%) | 25886 (53.87%) | 11123 (23.15%) | 3347 (6.97%) | 695 (1.45%) | 86 (0.18%) |
| RAPID | 43091 (89.68%) | 29215 (60.80%) | 14891 (30.99%) | 5774 (12.02%) | 1662 (3.46%) | 352 (0.73%) |
| MaxQuant | 38775 (80.70%) | 35 (0.07%) | 0 (0.00%) | 0 (0.00%) | 0 (0.00%) | 0 (0.00%) |
| Dinosaur | 43473 (90.48%) | 33358 (69.42%) | 20741 (43.17%) | 10059 (20.93%) | 3627 (7.55%) | 999 (2.08%) |
| PointIso | 38421 (79.96%) | 24682 (51.37%) | 12637 (26.30%) | 5046 (10.50%) | 1503 (3.13%) | 354 (0.74%) |
| EnumInst | 38554 (80.24%) | 3955 (8.23%) | 106 (0.22%) | 2 (0.00%) | 0 (0.00%) | 0 (0.00%) |
| EnumIW | 44042 (91.66%) | 33301 (69.31%) | 20584 (42.84%) | 10034 (20.88%) | 3670 (7.64%) | 1072 (2.23%) |
| EnumEx | 44109 (91.80%) | 33583 (69.89%) | 20966 (43.63%) | 10287 (21.41%) | 3811 (7.93%) | 1111 (2.31%) |

Table S9: Dong-DSS-1.6

| #PSM per MS2 | ≥ 1 | ≥ 2 | ≥ 3 | ≥ 4 | ≥ 5 | ≥ 6 |
|---|---|---|---|---|---|---|
| PepPre | 44225 (14.53%) | 4326 (1.42%) | 221 (0.07%) | 8 (0.00%) | 0 (0.00%) | 0 (0.00%) |
| pParse | 39246 (12.89%) | 1758 (0.58%) | 79 (0.03%) | 3 (0.00%) | 0 (0.00%) | 0 (0.00%) |
| RawConverter | 31339 (10.30%) | 0 (0.00%) | 0 (0.00%) | 0 (0.00%) | 0 (0.00%) | 0 (0.00%) |
| Monocle | 30517 (10.03%) | 0 (0.00%) | 0 (0.00%) | 0 (0.00%) | 0 (0.00%) | 0 (0.00%) |
| Decon2LS | 3925 (1.29%) | 236 (0.08%) | 3 (0.00%) | 0 (0.00%) | 0 (0.00%) | 0 (0.00%) |
| RAPID | 5500 (1.81%) | 22 (0.01%) | 0 (0.00%) | 0 (0.00%) | 0 (0.00%) | 0 (0.00%) |
| MaxQuant | 22919 (7.53%) | 0 (0.00%) | 0 (0.00%) | 0 (0.00%) | 0 (0.00%) | 0 (0.00%) |
| Dinosaur | 28922 (9.50%) | 2835 (0.93%) | 158 (0.05%) | 20 (0.01%) | 1 (0.00%) | 1 (0.00%) |
| PointIso | 8082 (2.65%) | 5123 (1.68%) | 3050 (1.00%) | 1789 (0.59%) | 913 (0.30%) | 413 (0.14%) |
| EnumInst | 31817 (10.45%) | 454 (0.15%) | 2 (0.00%) | 0 (0.00%) | 0 (0.00%) | 0 (0.00%) |
| EnumIW | 33566 (11.03%) | 1462 (0.48%) | 87 (0.03%) | 2 (0.00%) | 0 (0.00%) | 0 (0.00%) |
| EnumEx | 38924 (12.79%) | 2664 (0.88%) | 158 (0.05%) | 7 (0.00%) | 0 (0.00%) | 0 (0.00%) |

### Center Ion vs. Non-center Ion

Table S10

| | Zubarev-Human-2 | | Zubarev-Human-8 | | Dong-DSS-1.6 | |
|---|---|---|---|---|---|---|
| | Center | Non-center | Center | Non-center | Center | Non-center |
| PepPre | 42452 (58.21%) | 30482 (41.79%) | 40990 (34.17%) | 78953 (65.83%) | 33149 (67.96%) | 15631 (32.04%) |
| pParse | 43205 (61.83%) | 26672 (38.17%) | 40372 (37.14%) | 68343 (62.86%) | 32165 (78.29%) | 8921 (21.71%) |
| RawConverter | 38750 (97.23%) | 1106 (2.77%) | 38534 (97.68%) | 914 (2.32%) | 28429 (90.71%) | 2910 (9.29%) |
| Monocle | 39779 (99.74%) | 104 (0.26%) | 39537 (99.85%) | 58 (0.15%) | 30214 (99.01%) | 303 (0.99%) |
| Decon2LS | 35345 (74.89%) | 11851 (25.11%) | 33394 (39.91%) | 50282 (60.09%) | 3113 (74.76%) | 1051 (25.24%) |
| RAPID | 37355 (67.25%) | 18193 (32.75%) | 34617 (36.42%) | 60428 (63.58%) | 4319 (78.21%) | 1203 (21.79%) |
| MaxQuant | 39115 (99.74%) | 100 (0.26%) | 38755 (99.86%) | 55 (0.14%) | 21406 (93.40%) | 1513 (6.60%) |
| Dinosaur | 37766 (61.55%) | 23588 (38.45%) | 37400 (33.24%) | 75110 (66.76%) | 24049 (75.30%) | 7888 (24.70%) |
| PointIso | 28902 (61.27%) | 18273 (38.73%) | 28901 (34.94%) | 53810 (65.06%) | 13598 (69.24%) | 6041 (30.76%) |
| EnumInst | 42390 (99.72%) | 120 (0.28%) | 42445 (99.60%) | 172 (0.40%) | 31425 (97.37%) | 848 (2.63%) |
| EnumIW | 42013 (62.89%) | 24794 (37.11%) | 39900 (35.32%) | 73076 (64.68%) | 26197 (74.60%) | 8920 (25.40%) |
| EnumEx | 42682 (61.03%) | 27257 (38.97%) | 39744 (34.82%) | 74410 (65.18%) | 29871 (71.54%) | 11882 (28.46%) |

### Case #1

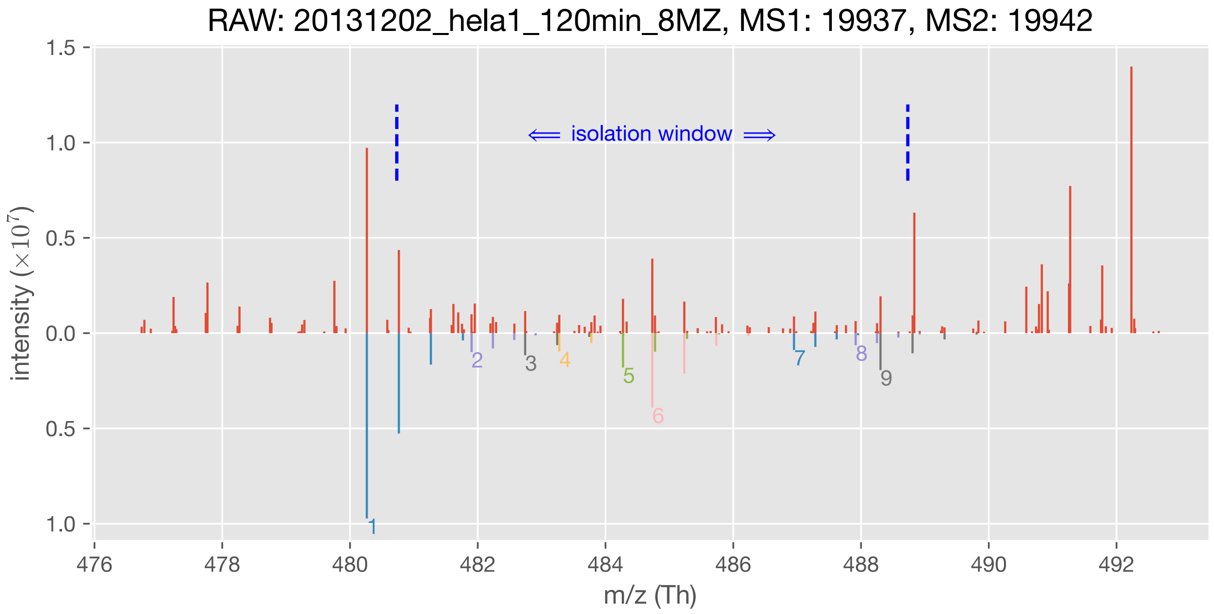


| | m/z (Th) | charge state | sequence | modification | q-value |
|---|---|---|---|---|---|
| #1 ion | 480.263305 | 2 | SLEVLENR | - | 0.08% |
| #2 ion | 481.903038 | 3 | SEPHSLSSEALMR | - | 0.11% |
| #3 ion | 482.742083 | 2 | FEEELAAR | - | 0.36% |
| #4 ion | 483.276243 | 2 | LLSQVSYR | - | 0.20% |
| #5 ion | 484.273943 | 2 | KEFFLQR | - | 0.46% |
| #6 ion | 484.731832 | 2 | NGDYLCVK | Carbamidomethyl[C]@6 | 0.02% |
| #7 ion | 486.950699 | 3 | KPLPDHVSLVEPK | - | 0.14% |
| #8 ion | 487.915765 | 3 | LSNRPAFMPSEGR | - | 0.45% |
| #9 ion | 488.305021 | 2 | KGVALNFVK | - | 0.11% |

Figure S15: Case #1

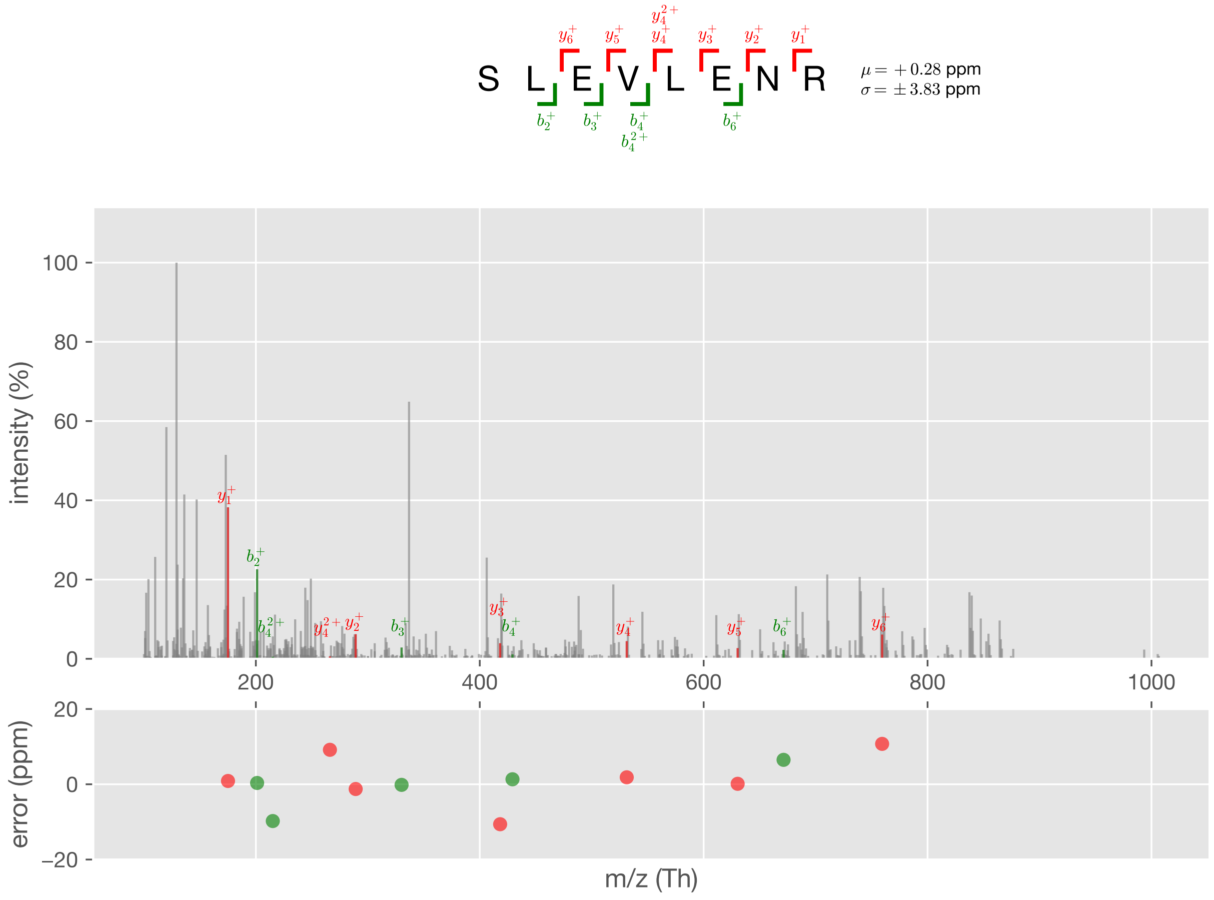


Figure S16: Case #1, PSM #1

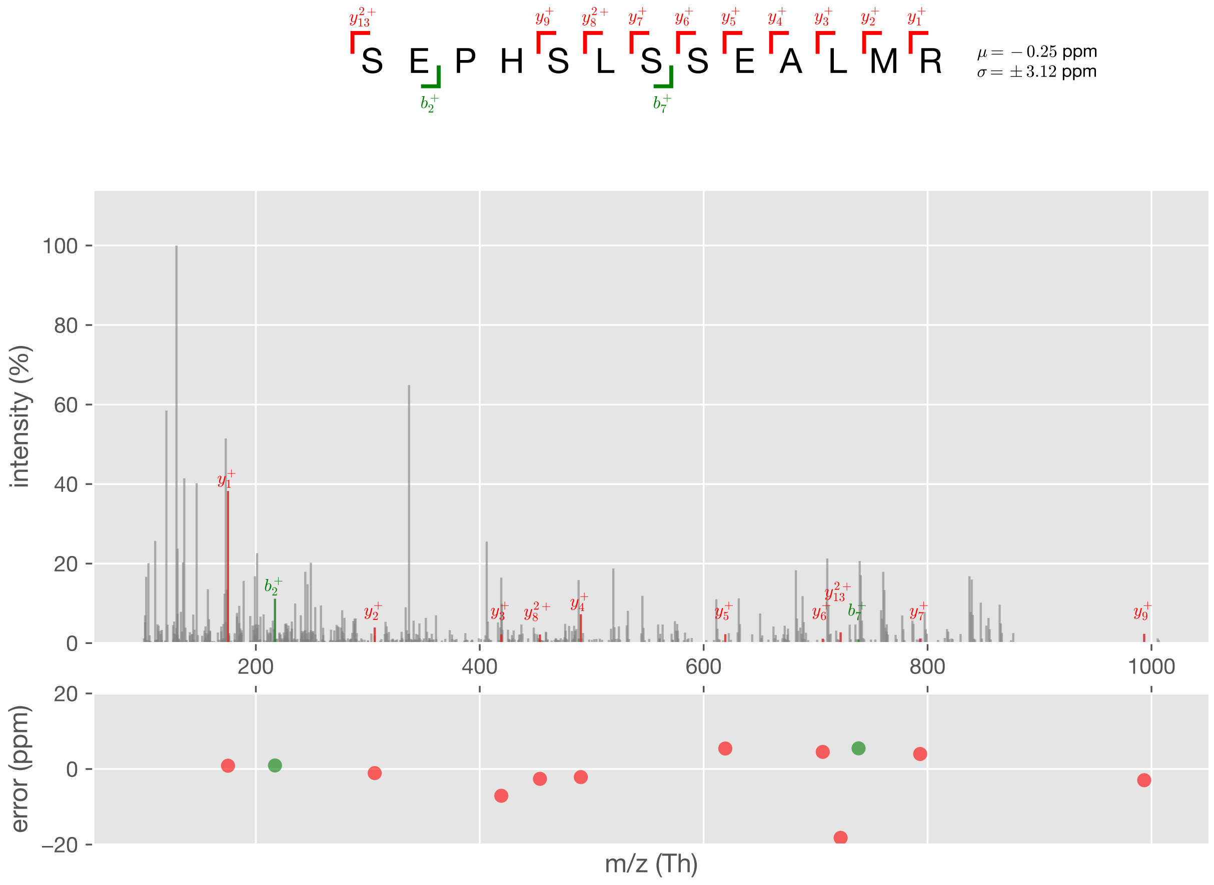


Figure S17: Case #1, PSM #2

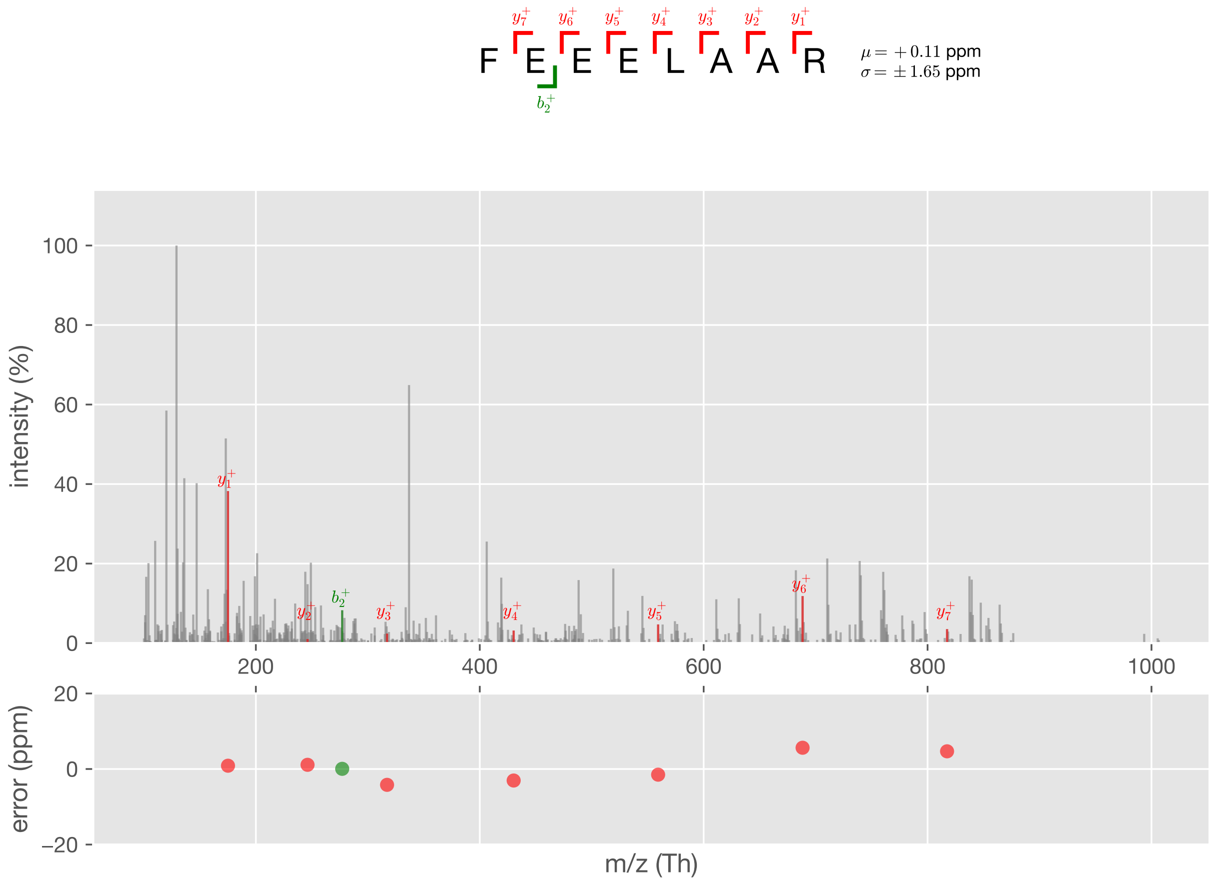


Figure S18: Case #1, PSM #3

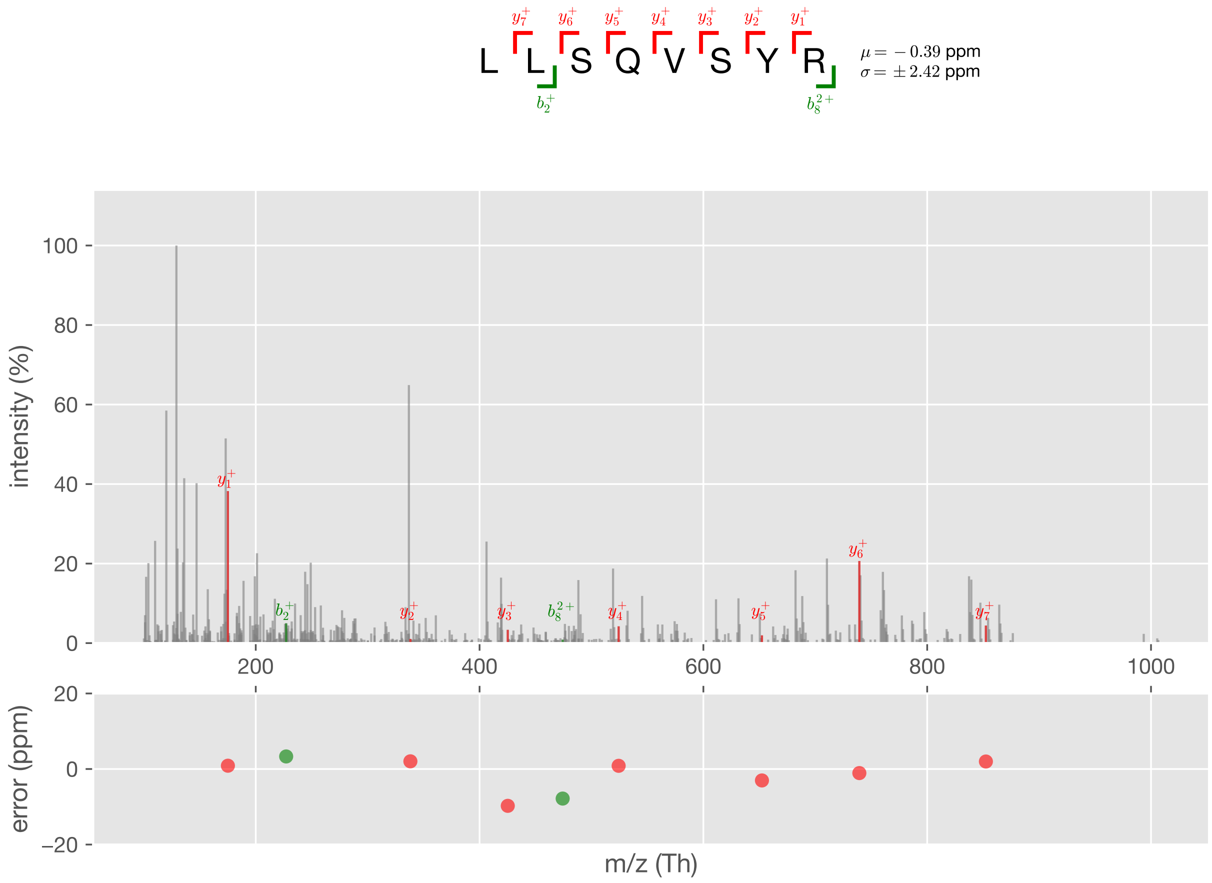


Figure S19: Case #1, PSM #4

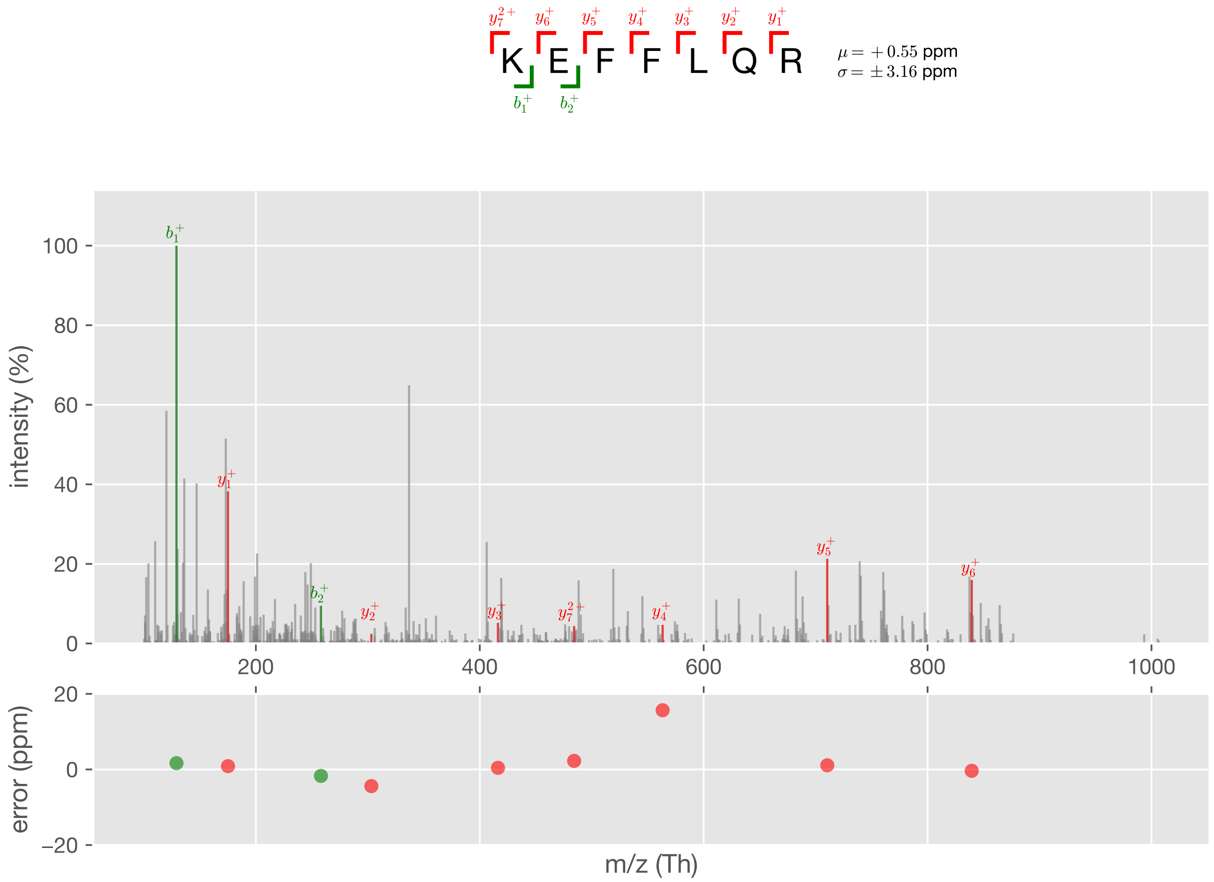


Figure S20: Case #1, PSM #5

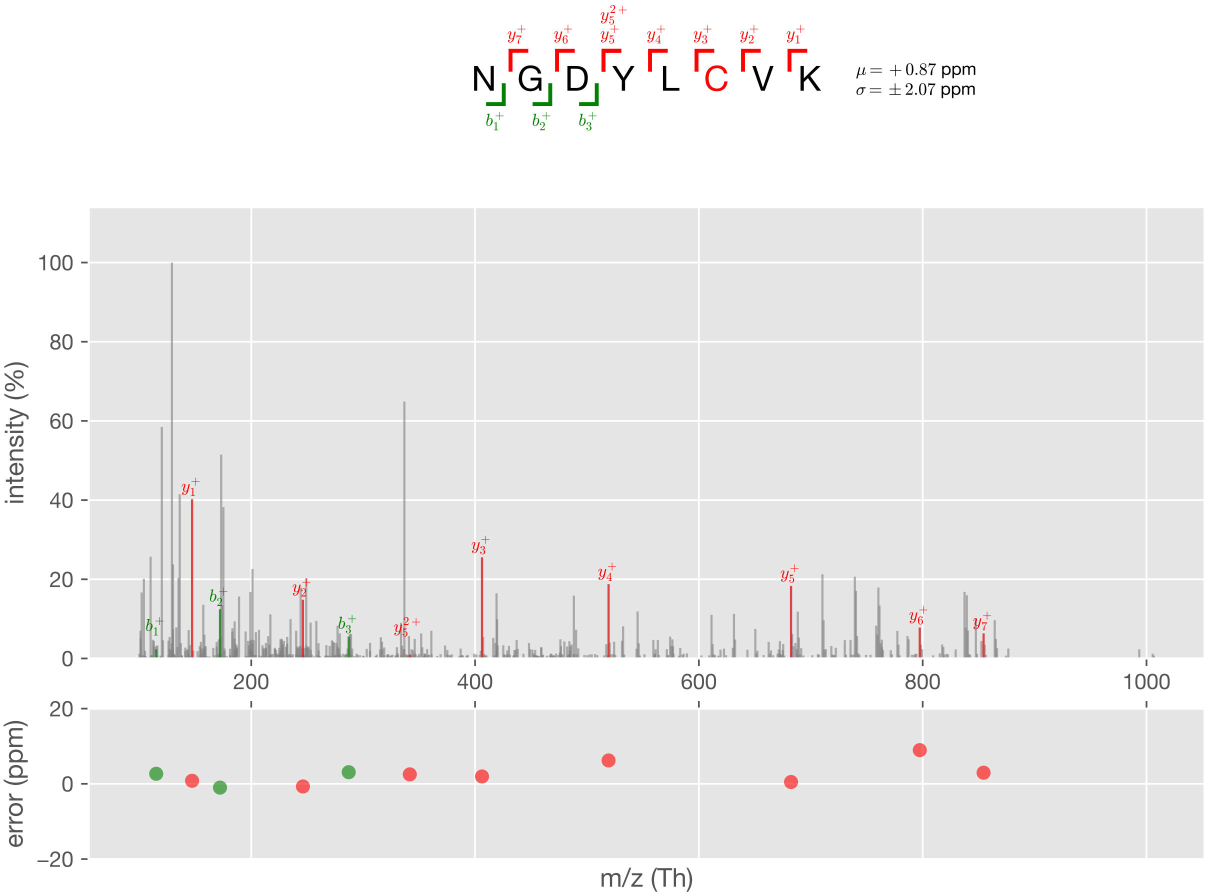


Figure S21: Case #1, PSM #6

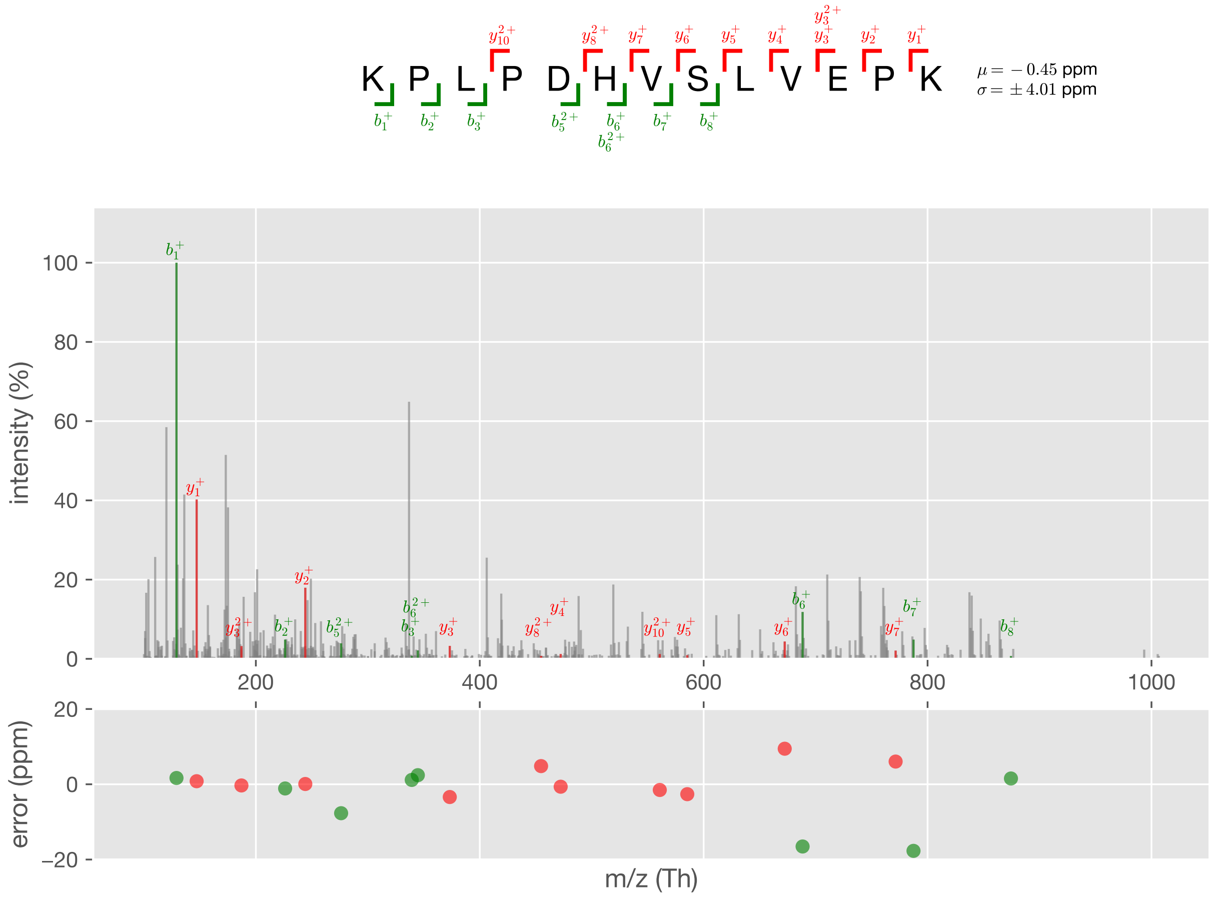


Figure S22: Case #1, PSM #7

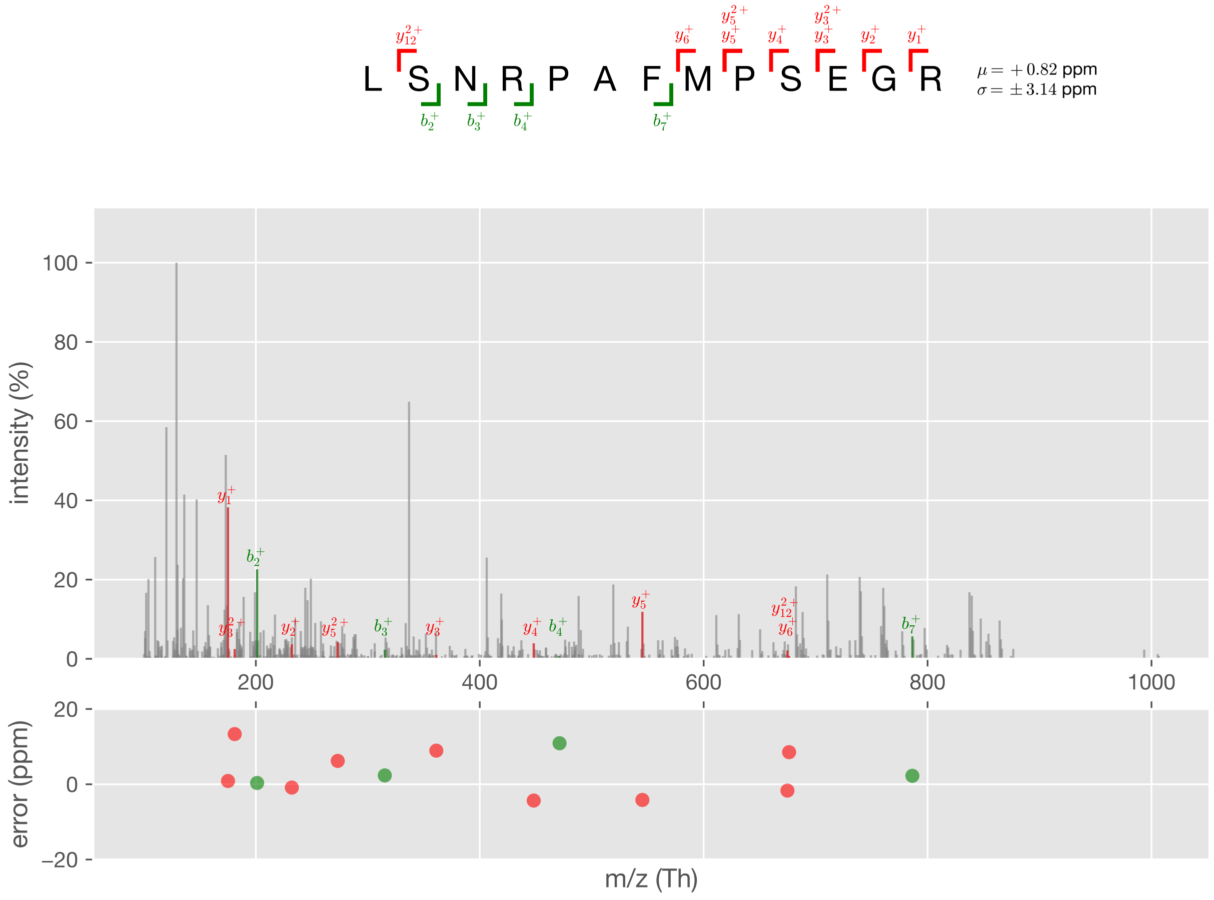


Figure S23: Case #1, PSM #8

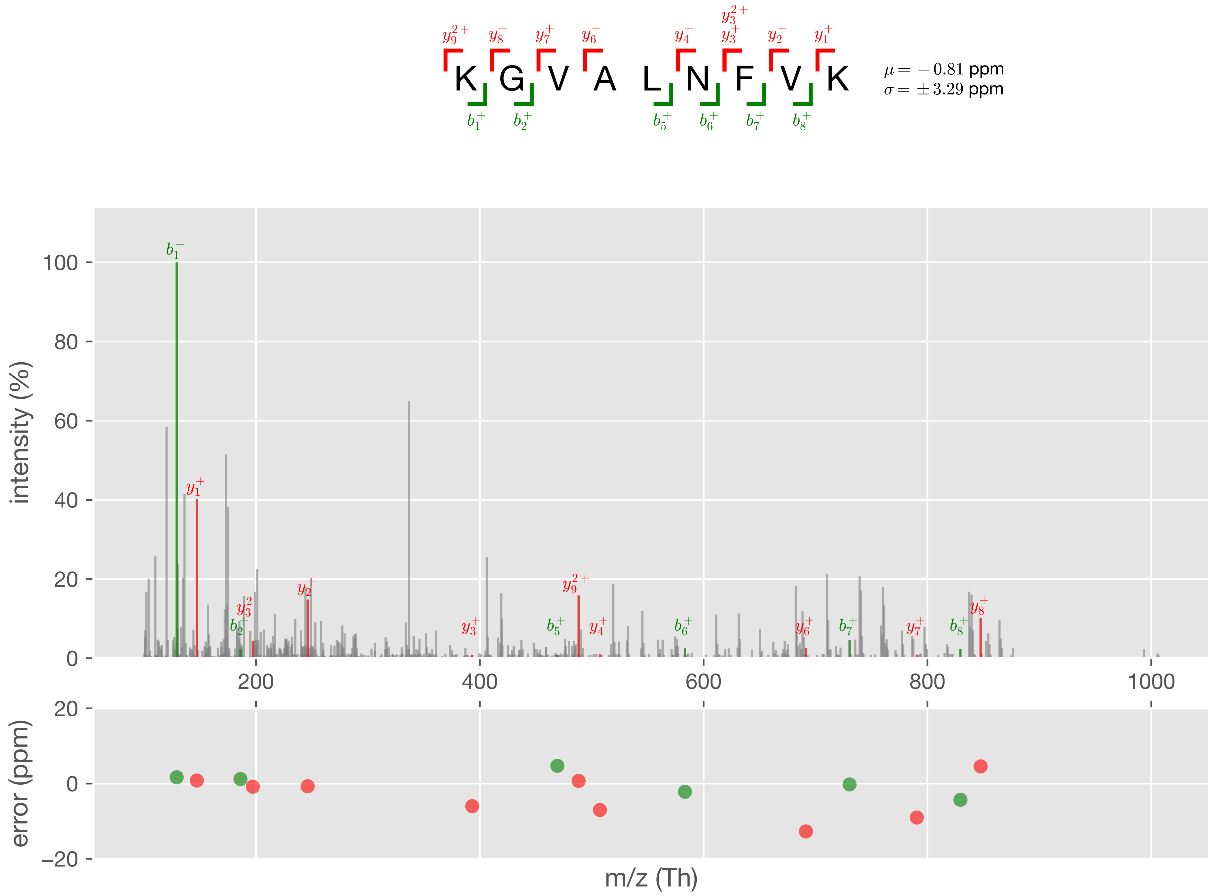


Figure S24: Case #1, PSM #9

### Case #2

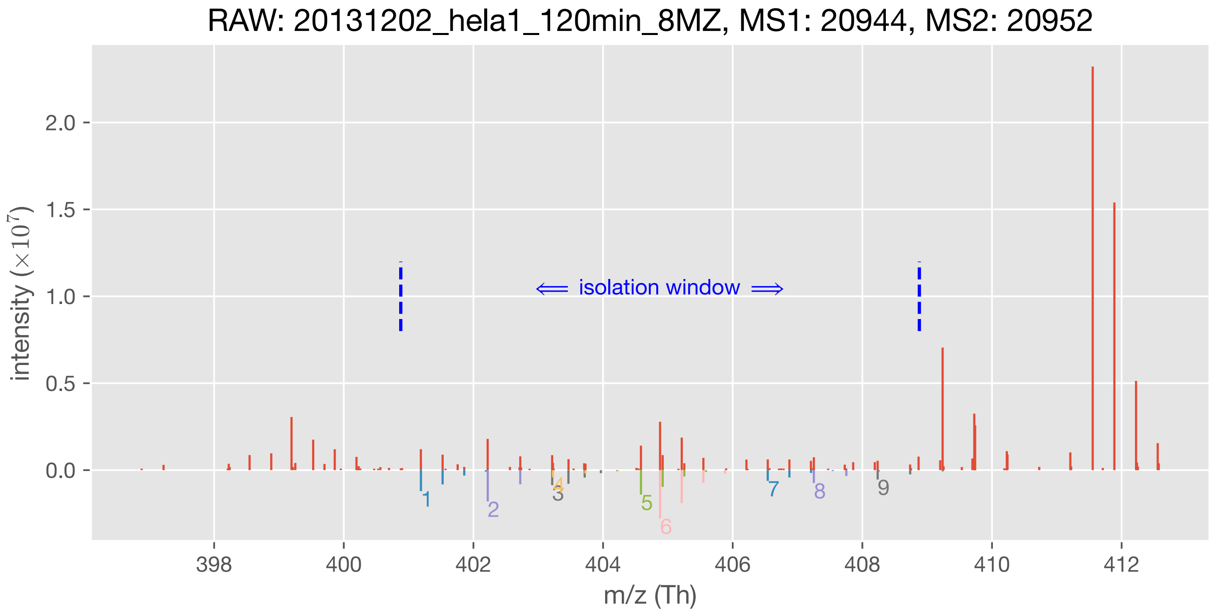


| | m/z (Th) | charge state | sequence | modification | q-value |
|---|---|---|---|---|---|
| #1 ion | 401.190066 | 3 | DMLDSHLENK | - | 0.37% |
| #2 ion | 402.220828 | 2 | LSDMPLK | - | 0.18% |
| #3 ion | 403.215074 | 4 | HLAPDHLLQLCHR | Carbamidomethyl[C]@11 | 0.01% |
| #4 ion | 403.227469 | 2 | FEVVPSK | - | 0.06% |
| #5 ion | 404.585073 | 3 | LSNLALVKPEK | - | 0.02% |
| #6 ion | 404.879165 | 3 | VTALCWHPTK | Carbamidomethyl[C]@5 | 0.00% |
| #7 ion | 406.540202 | 3 | VVDLMAHMASK | Oxidation[M]@5 | 0.05% |
| #8 ion | 407.251573 | 2 | NQLAVLR | - | 0.13% |
| #9 ion | 408.236325 | 2 | KGSGLPSK | Acetyl[AnyN-term]@1 | 0.23% |

Figure S25: Case #2

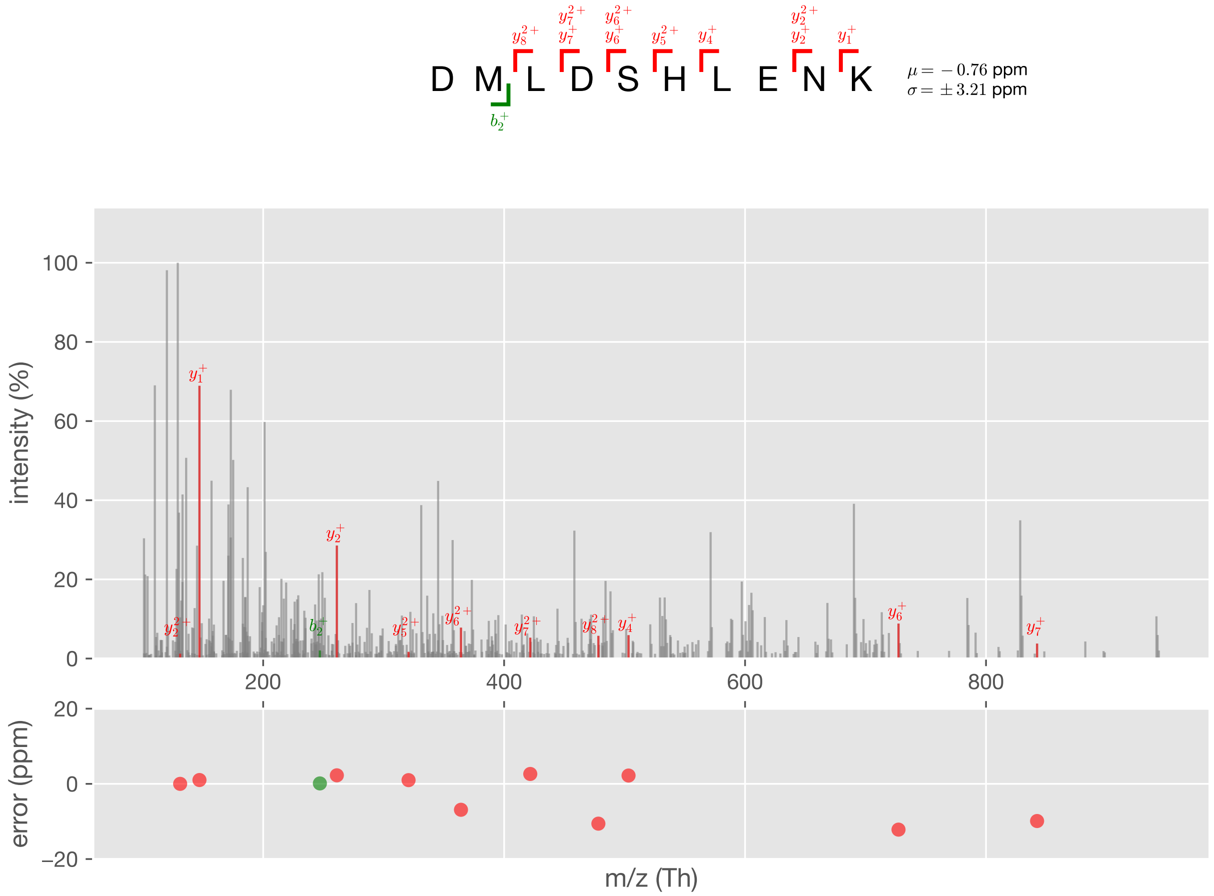


Figure S26: Case #2, PSM #1

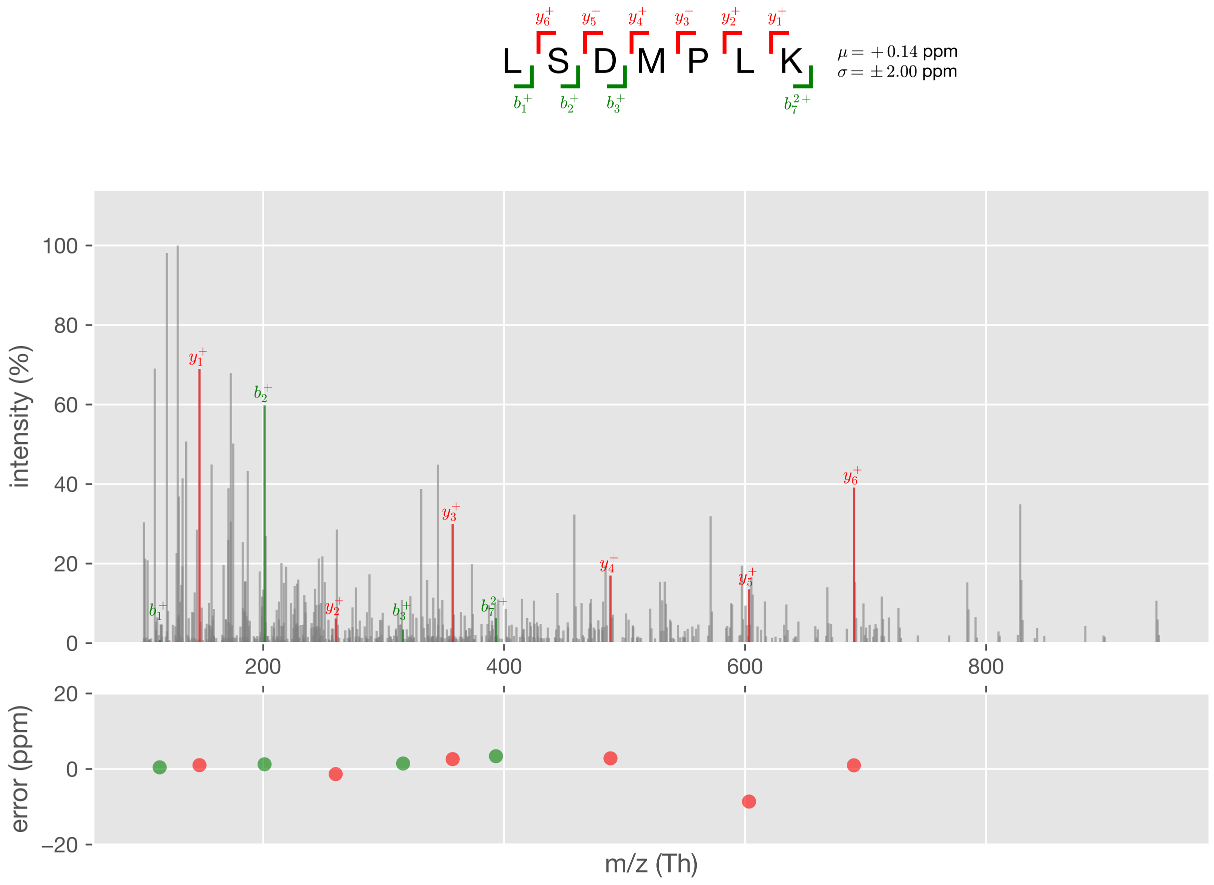


Figure S27: Case #2, PSM #2

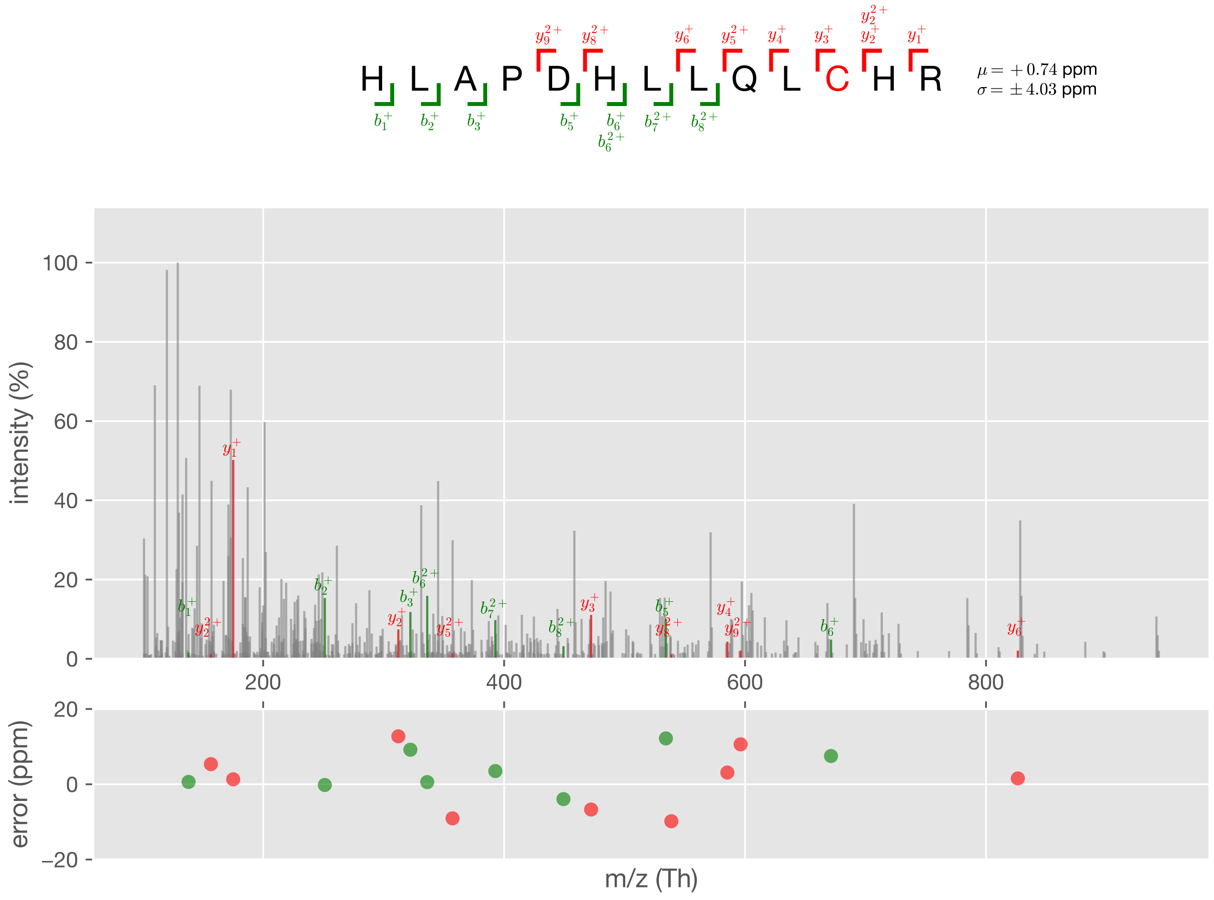


Figure S28: Case #2, PSM #3

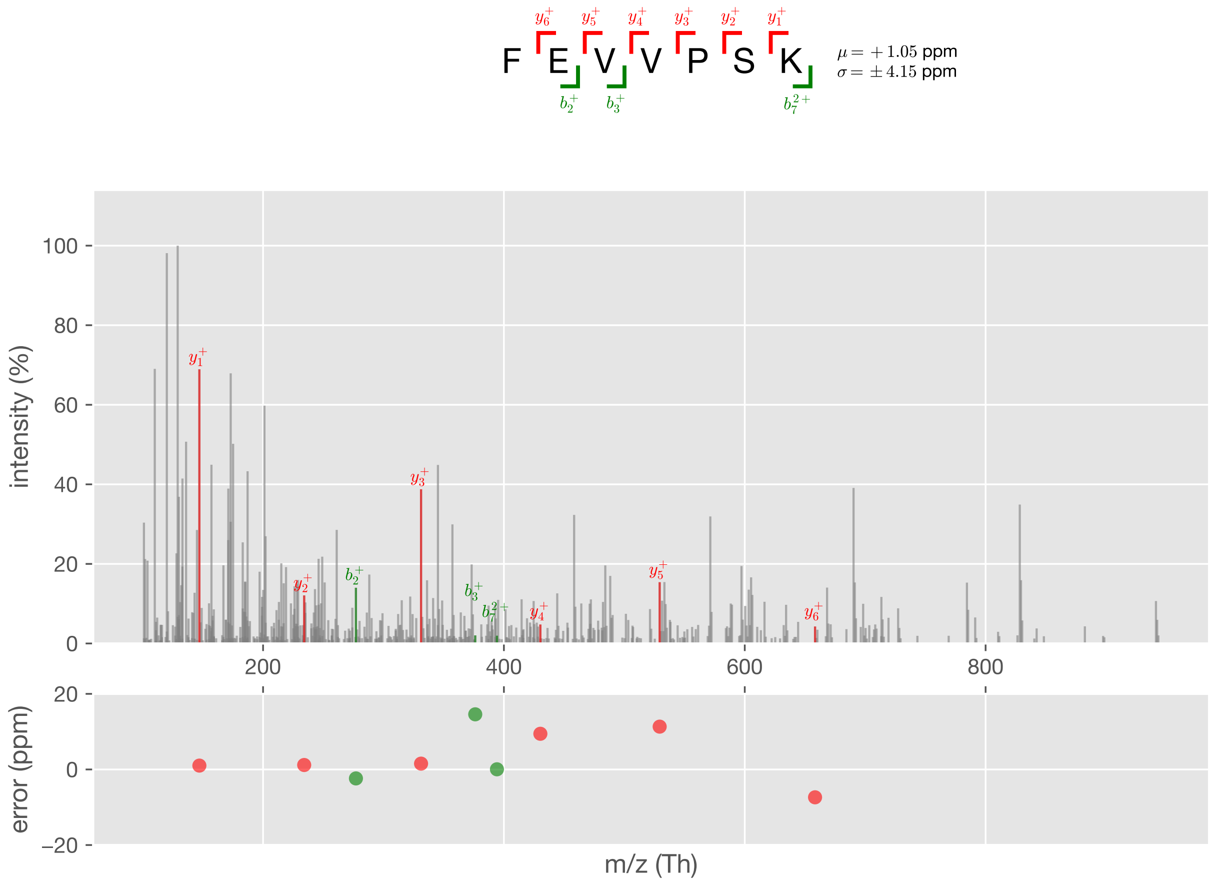


Figure S29: Case #2, PSM #4

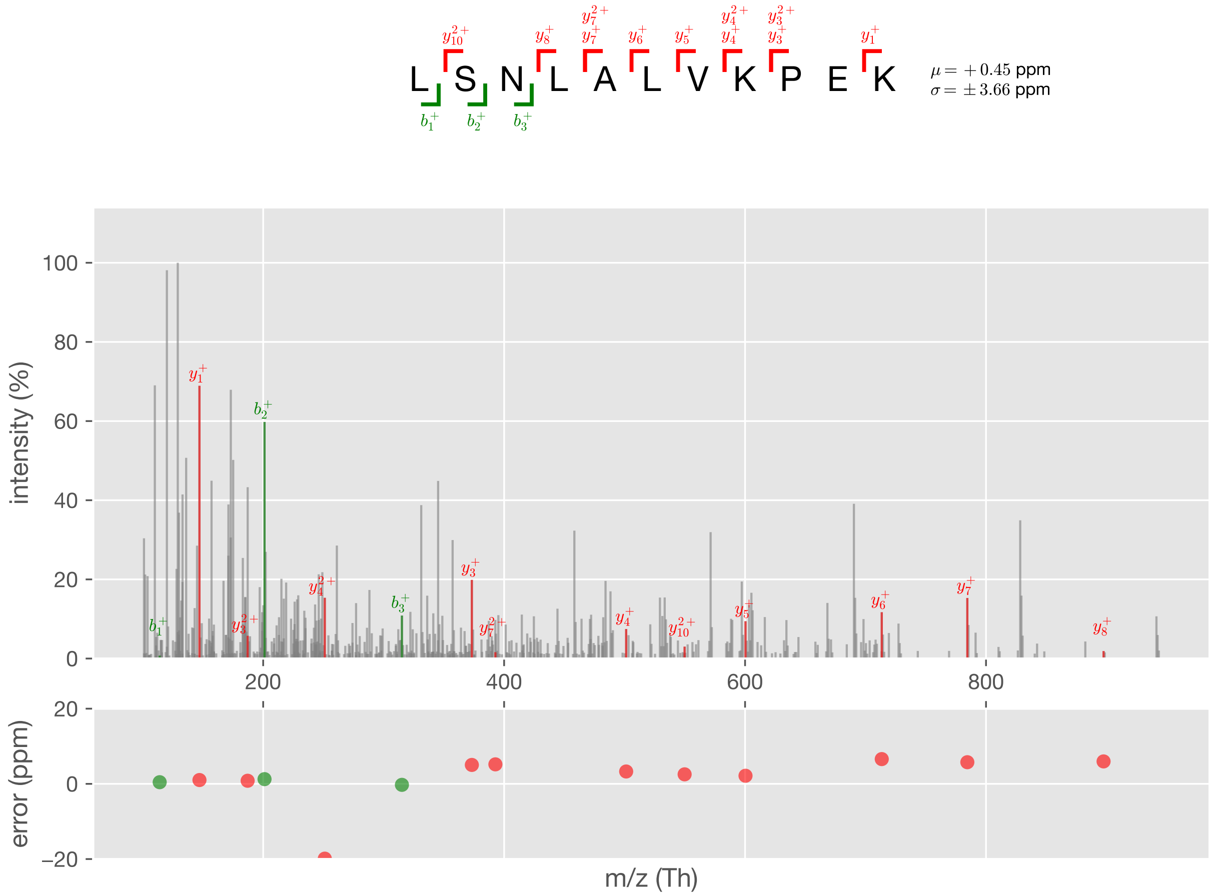


Figure S30: Case #2, PSM #5

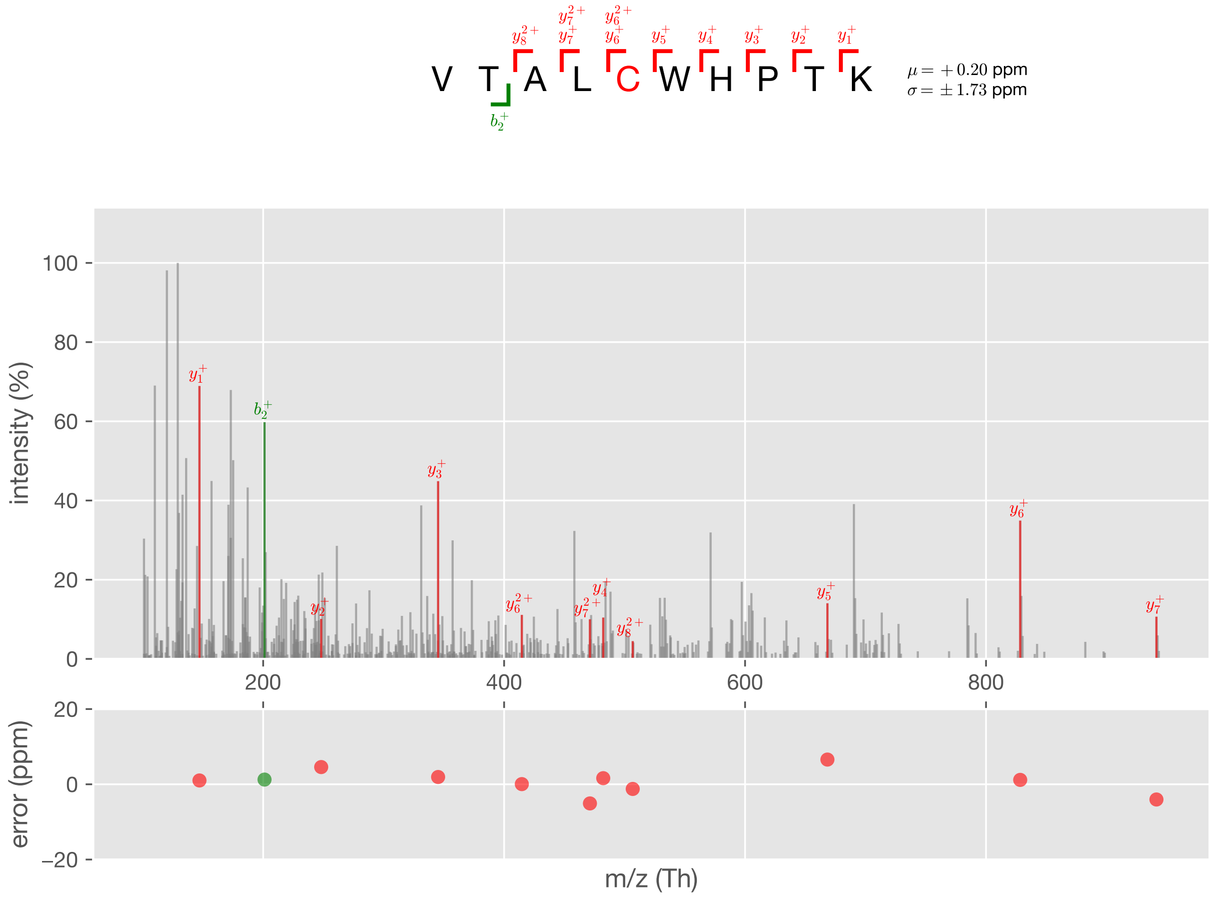


Figure S31: Case #2, PSM #6

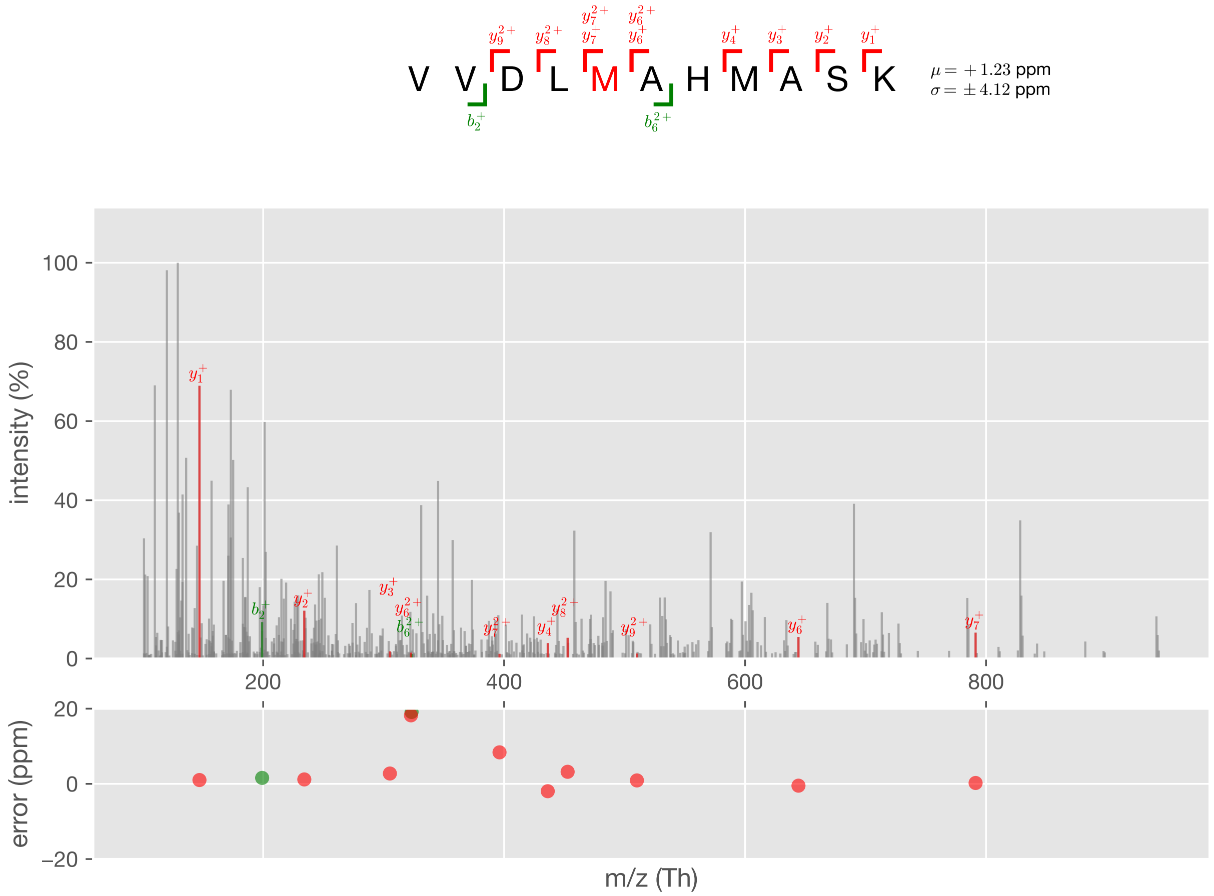


Figure S32: Case #2, PSM #7

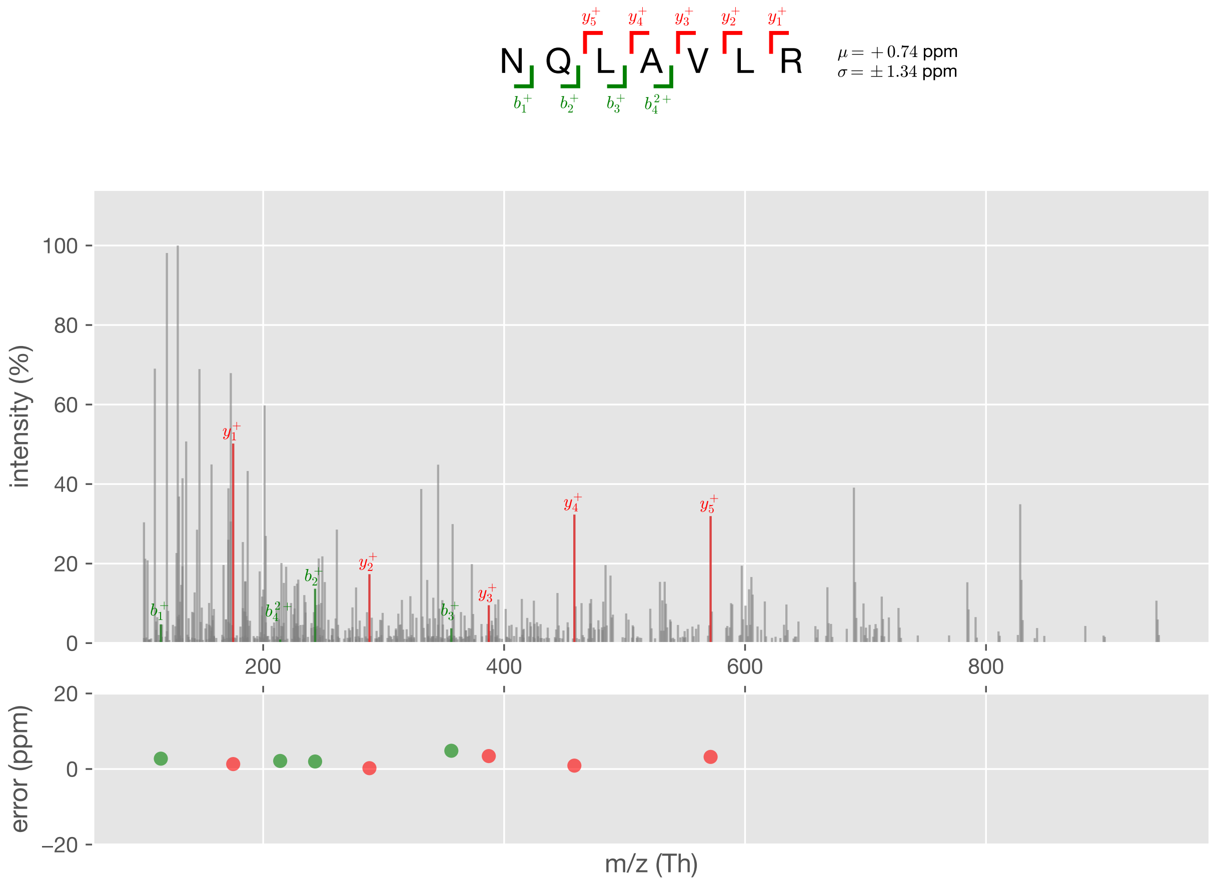


Figure S33: Case #2, PSM #8

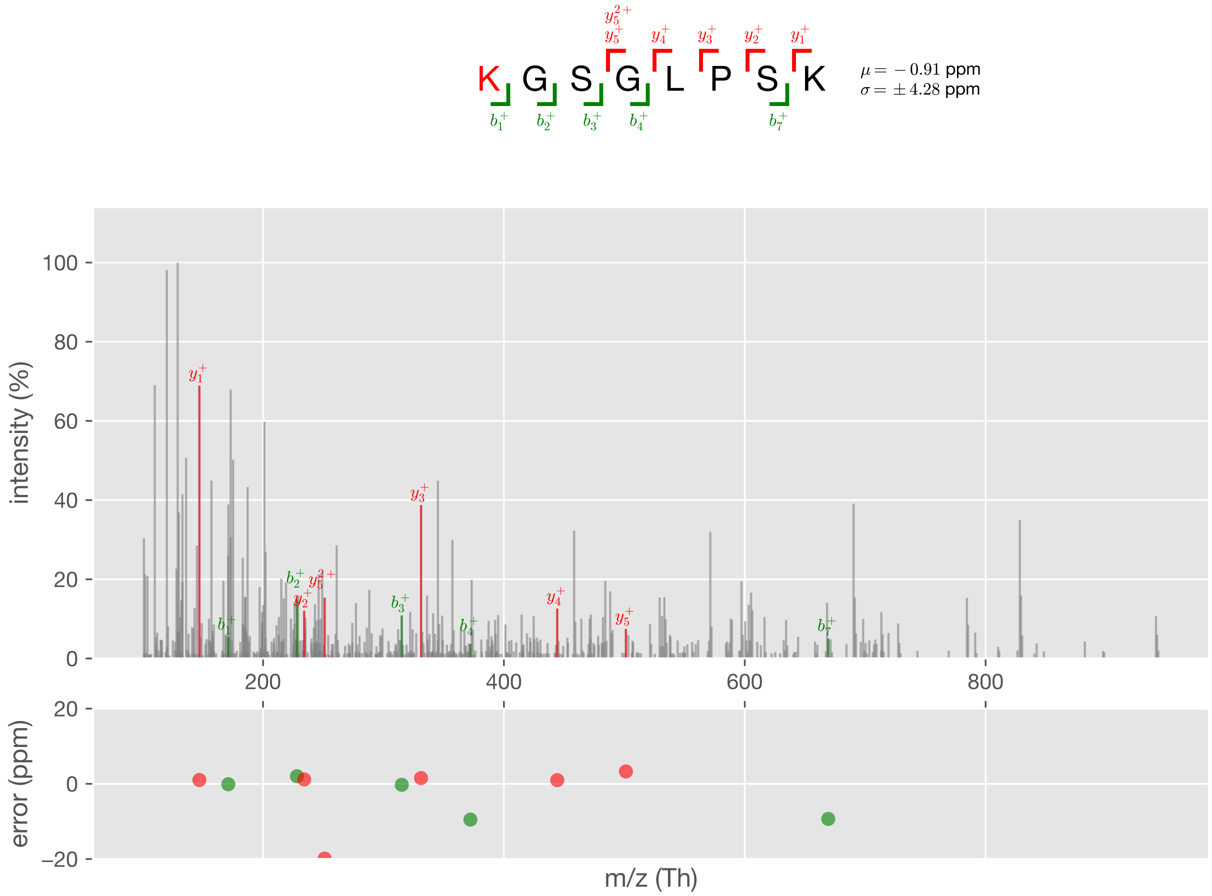


Figure S34: Case #2, PSM #9

### Case #3

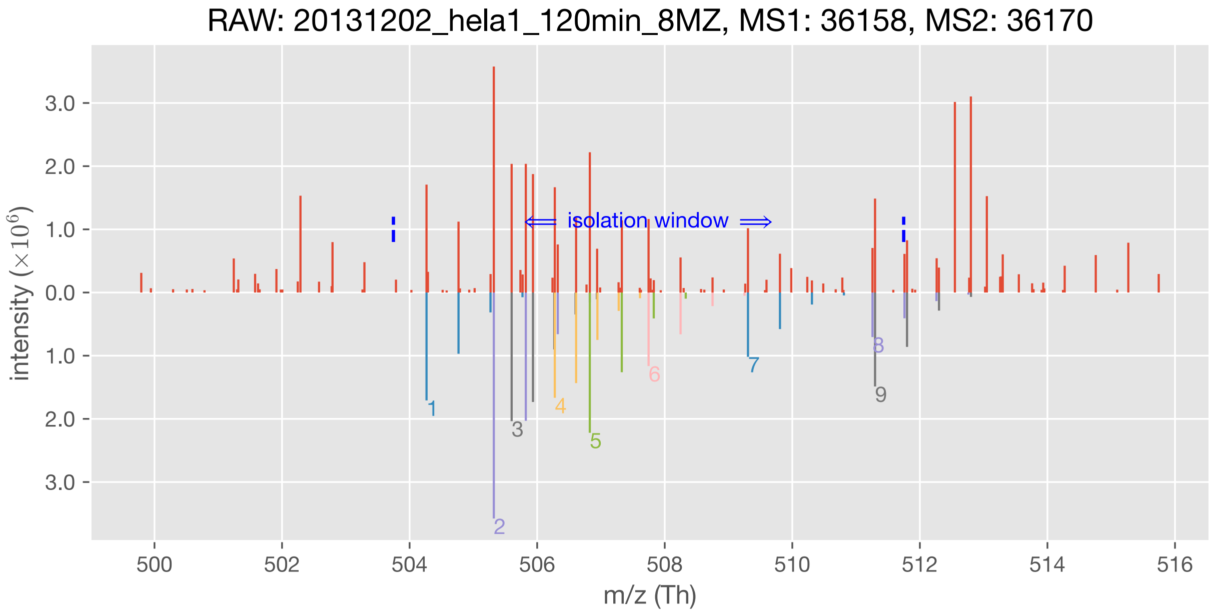


| | m/z (Th) | charge state | sequence | modification | q-value |
|---|---|---|---|---|---|
| #1 ion | 504.265343 | 2 | TWFSLPEK | - | 0.15% |
| #2 ion | 505.320387 | 2 | LLPLLTDPK | - | 0.02% |
| #3 ion | 505.599800 | 3 | NHPGLLLMDTTFR | - | 0.01% |
| #4 ion | 506.276203 | 3 | EVLLFPAMKPEDK | - | 0.05% |
| #5 ion | 506.825053 | 2 | VAAAVDLLLK | - | 0.01% |
| #6 ion | 507.746855 | 2 | DLTDYLMK | Oxidation[M]@7 | 0.44% |
| #7 ion | 509.305048 | 2 | VYLGDLPLK | - | 0.12% |
| #8 ion | 511.257180 | 2 | TEMDWVLK | - | 0.16% |
| #9 ion | 511.297306 | 2 | AGFVPVYLR | - | 0.95% |

Figure S35: Case #3

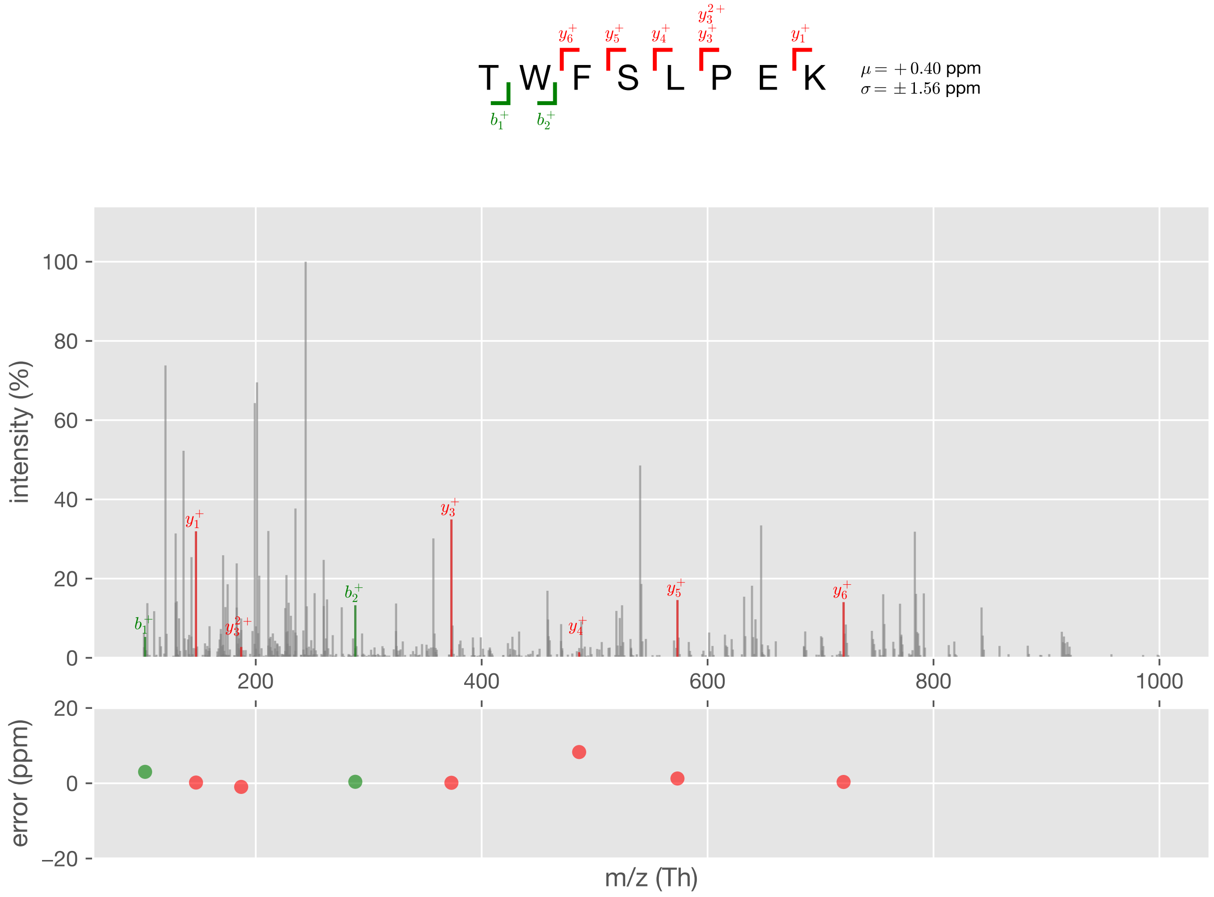


Figure S36: Case #3, PSM #1

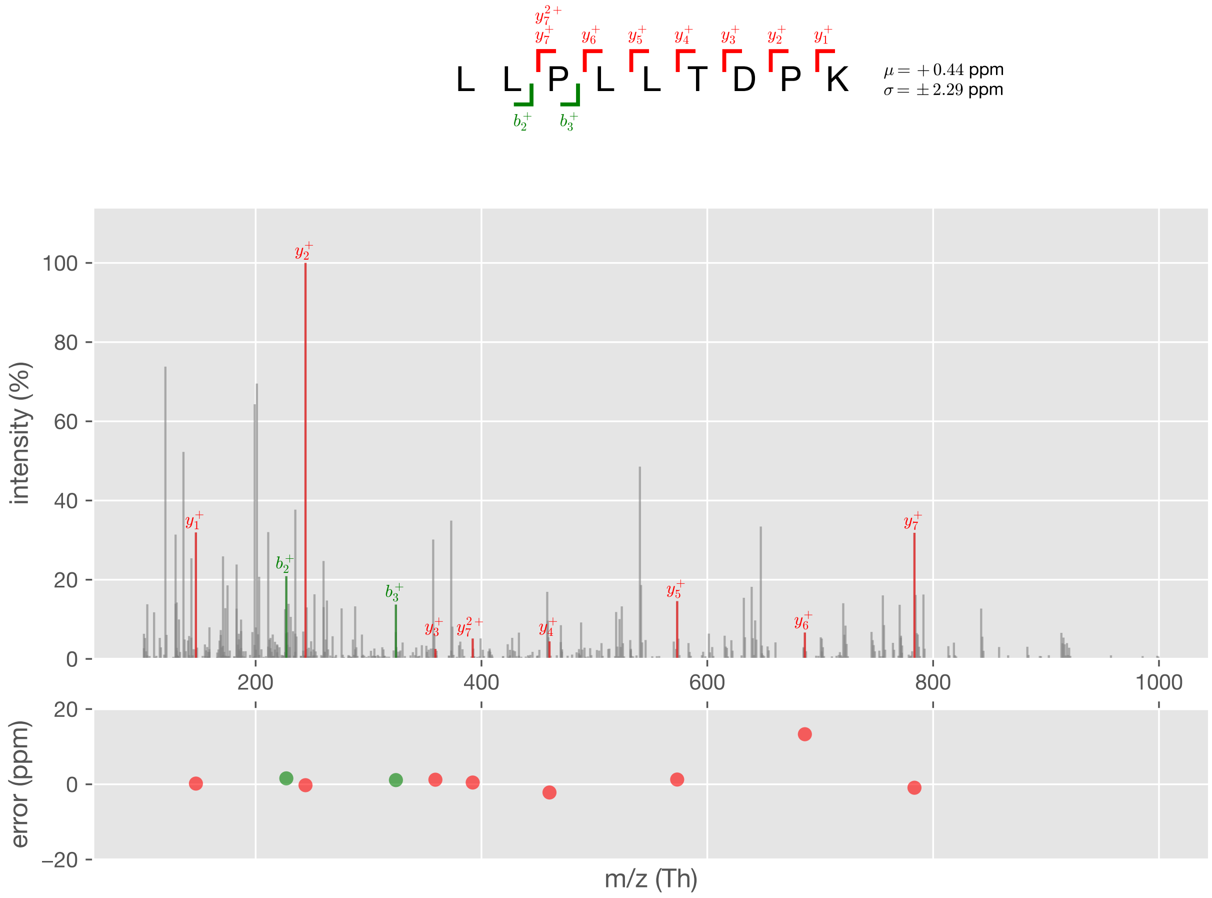


Figure S37: Case #3, PSM #2

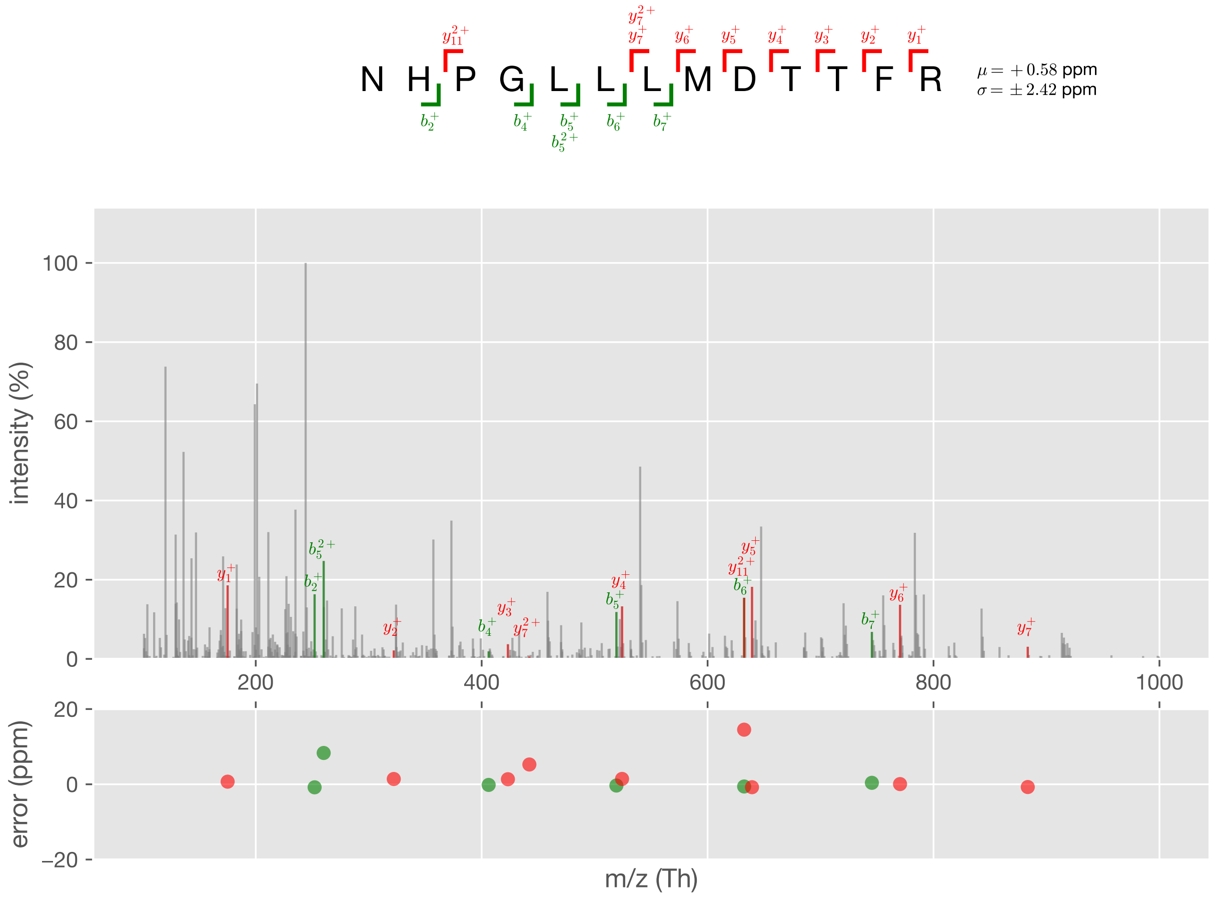


Figure S38: Case #3, PSM #3

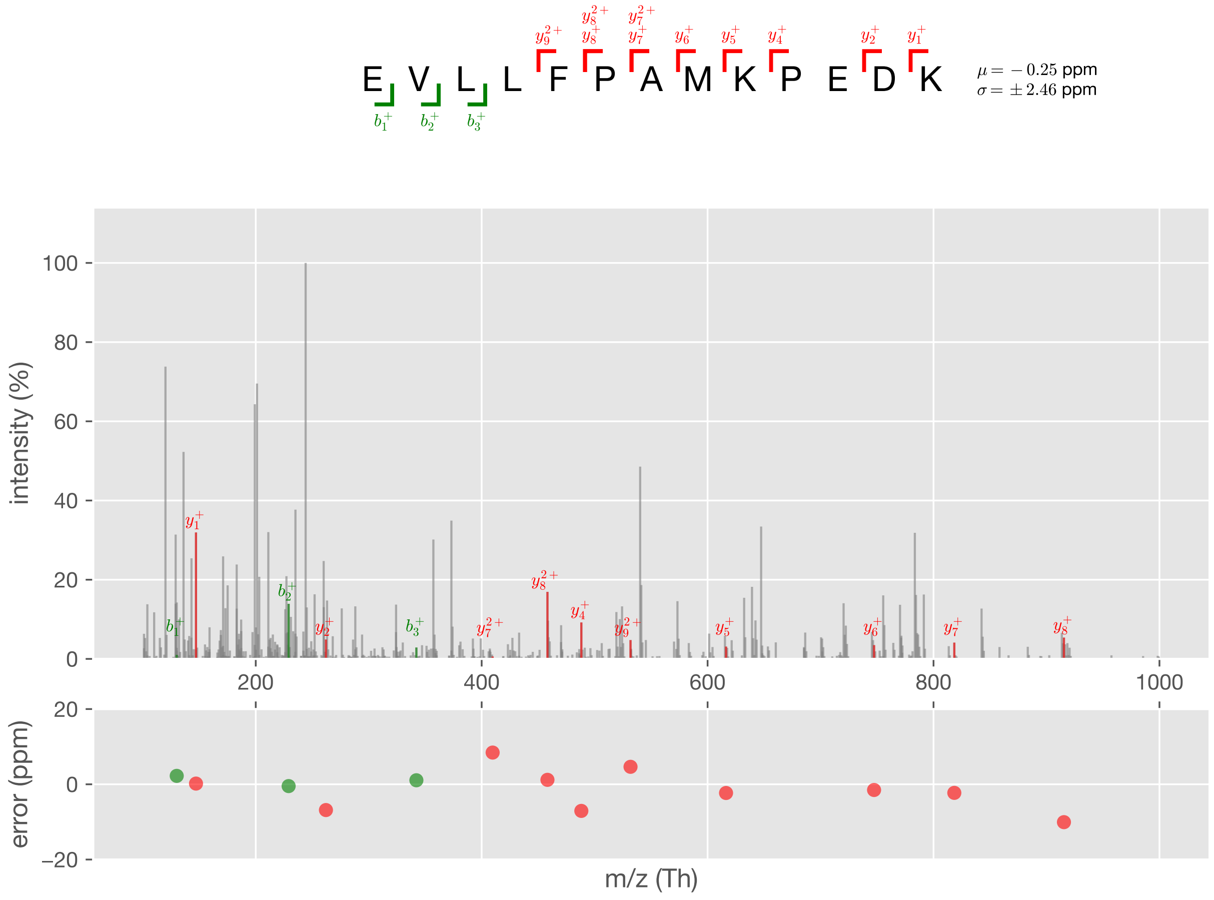


Figure S39: Case #3, PSM #4

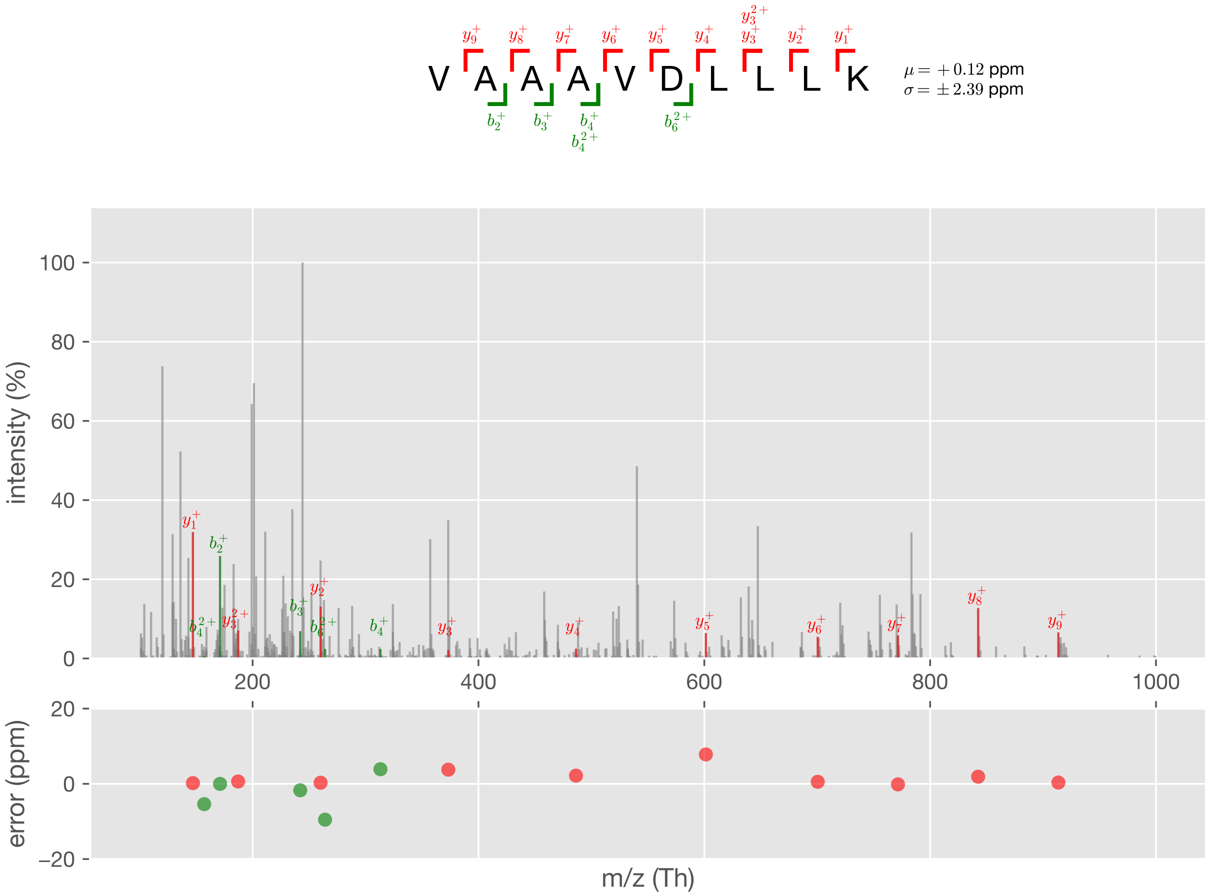


Figure S40: Case #3, PSM #5

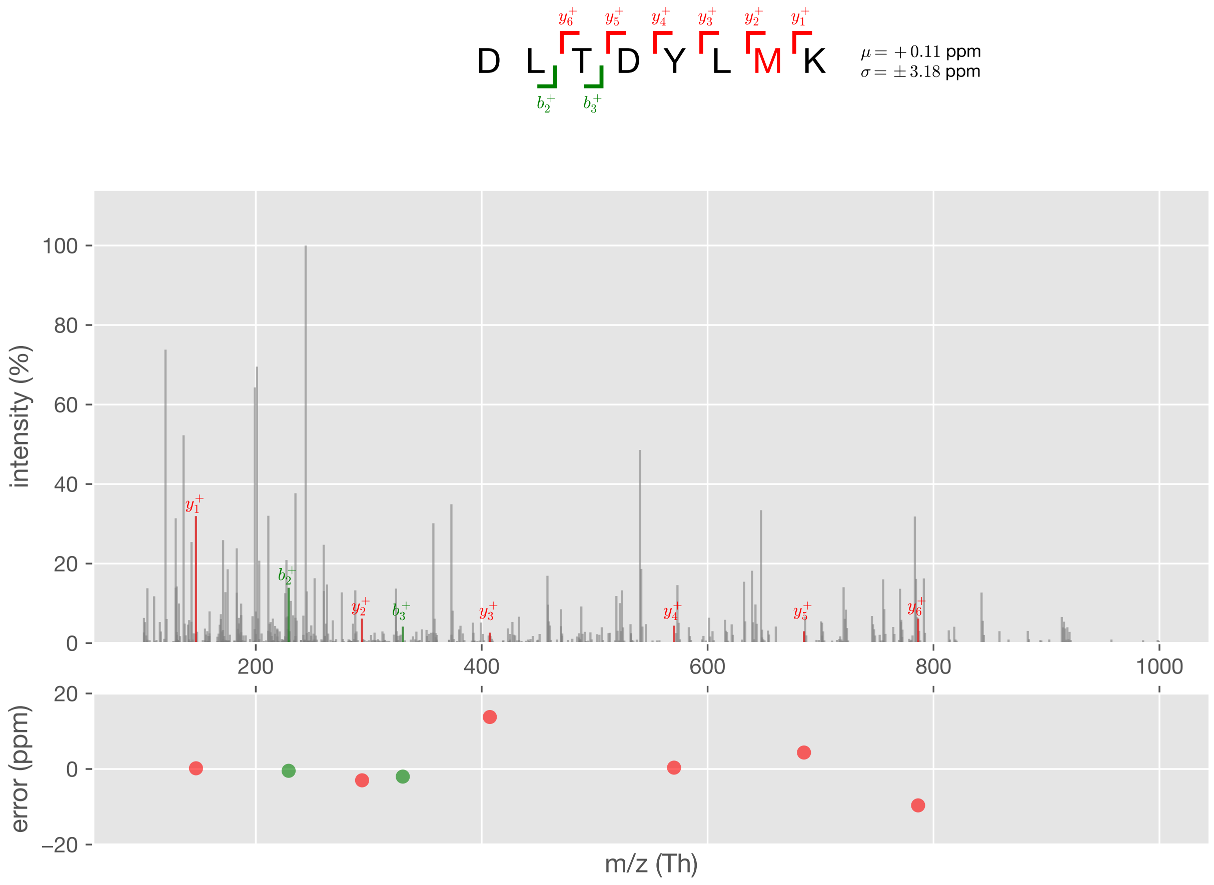


Figure S41: Case #3, PSM #6

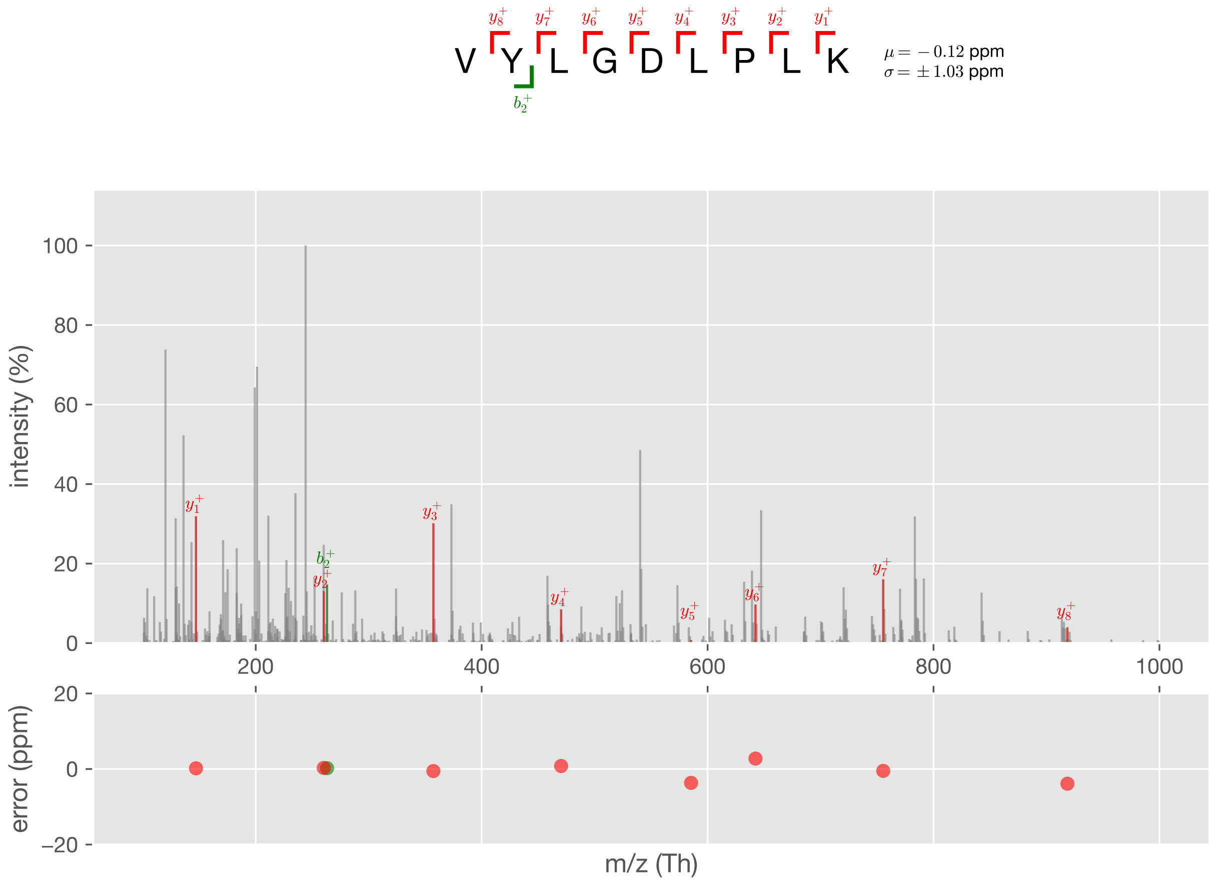


Figure S42: Case #3, PSM #7

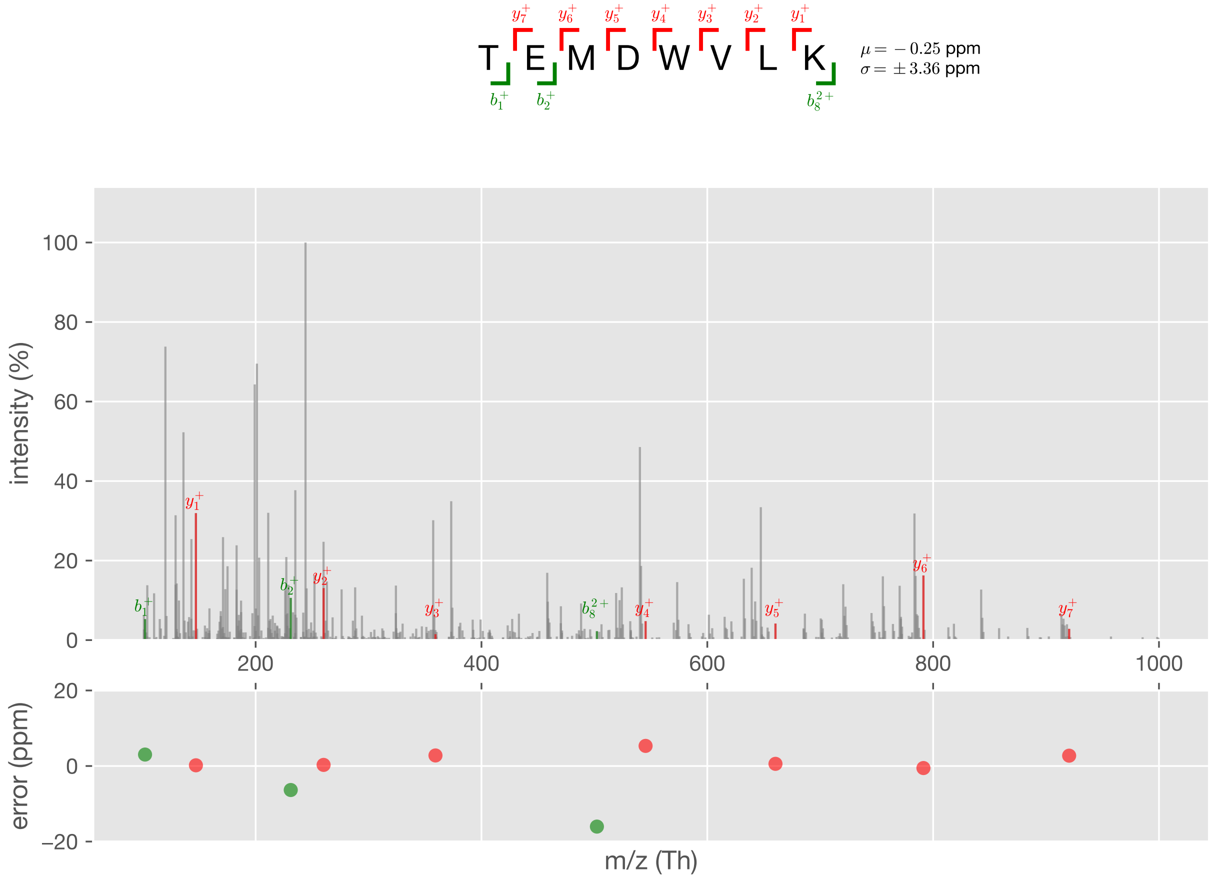


Figure S43: Case #3, PSM #8

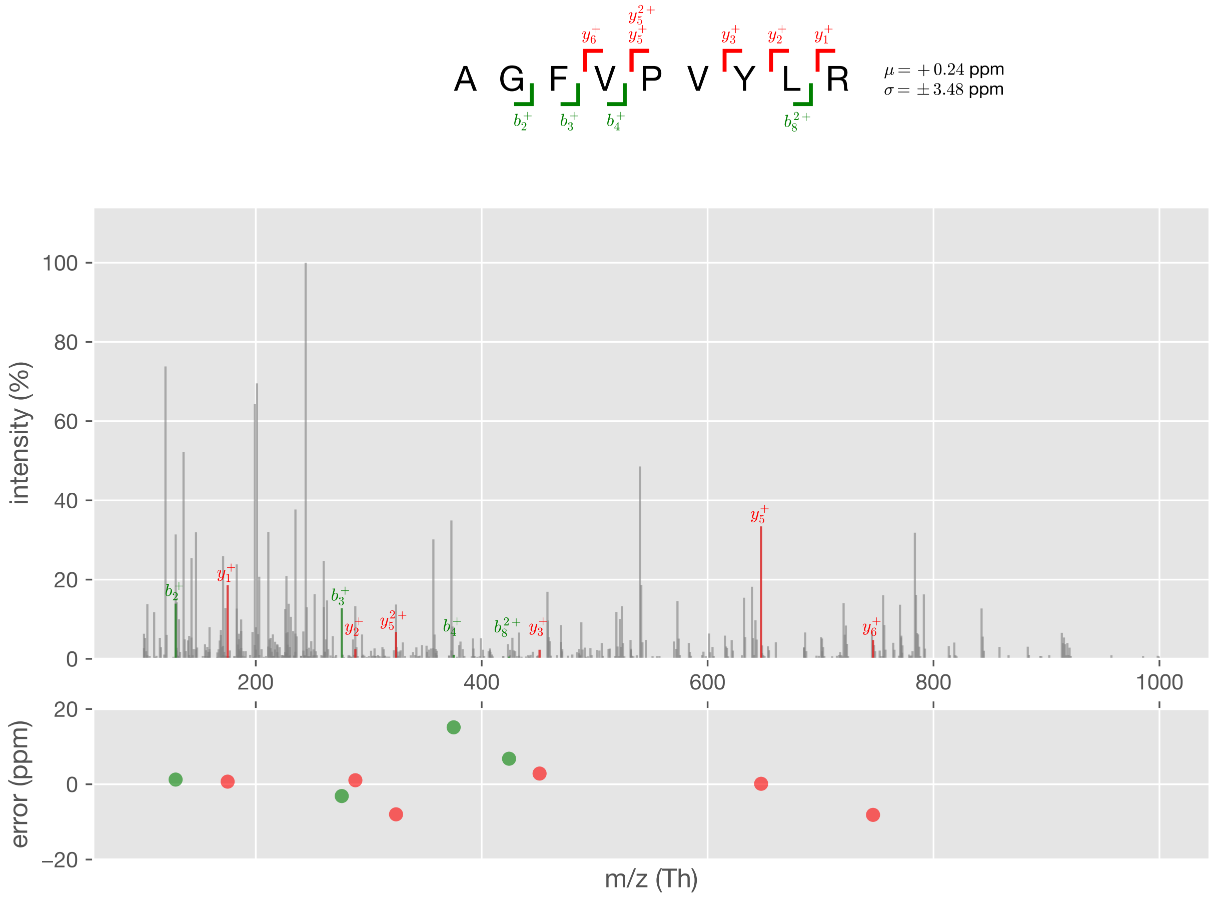


Figure S44: Case #3, PSM #9

### Case #4

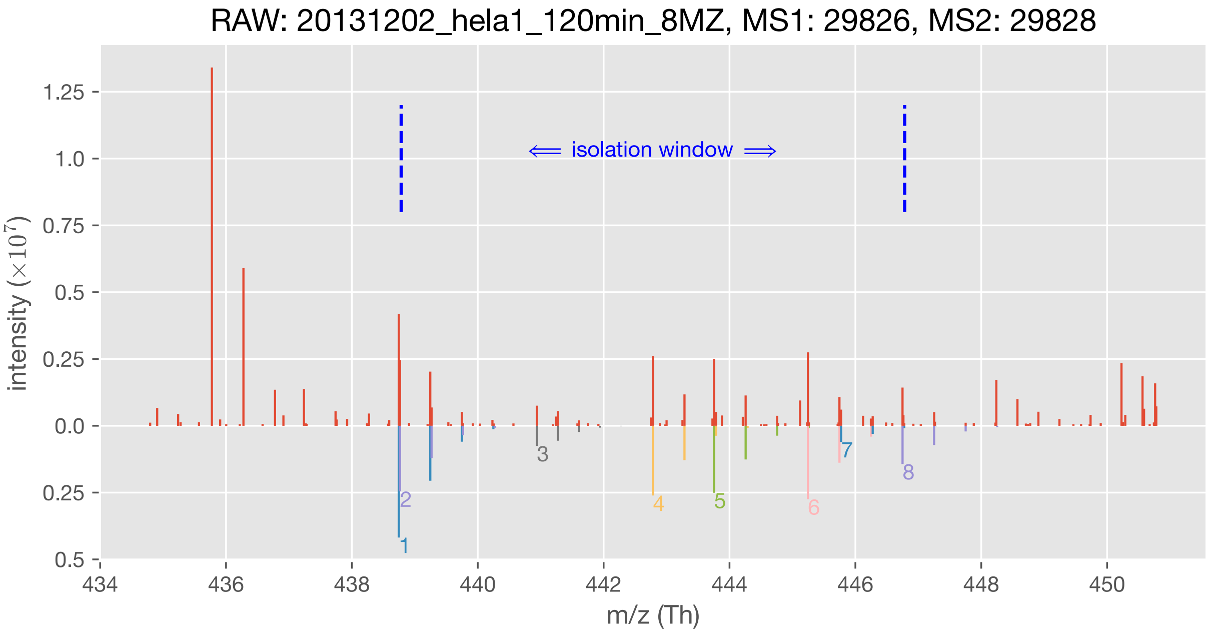


| | m/z (Th) | charge state | sequence | modification | q-value |
|---|---|---|---|---|---|
| #1 ion | 438.744359 | 2 | DLLSSVSR | - | 0.13% |
| #2 ion | 438.764244 | 2 | SLFSPVVK | - | 0.29% |
| #3 ion | 440.939563 | 3 | LLAHAGSLLNLAK | - | 0.08% |
| #4 ion | 442.783128 | 2 | LLLLAEGR | - | 0.00% |
| #5 ion | 443.754954 | 2 | VDAAELLR | - | 0.00% |
| #6 ion | 445.245714 | 2 | NGVDLLMK | - | 0.00% |
| #7 ion | 445.773660 | 2 | TLVLLMGK | Oxidation[M]@6 | 0.12% |
| #8 ion | 446.749400 | 2 | AYQELLR | - | 0.15% |

Figure S45: Case #4

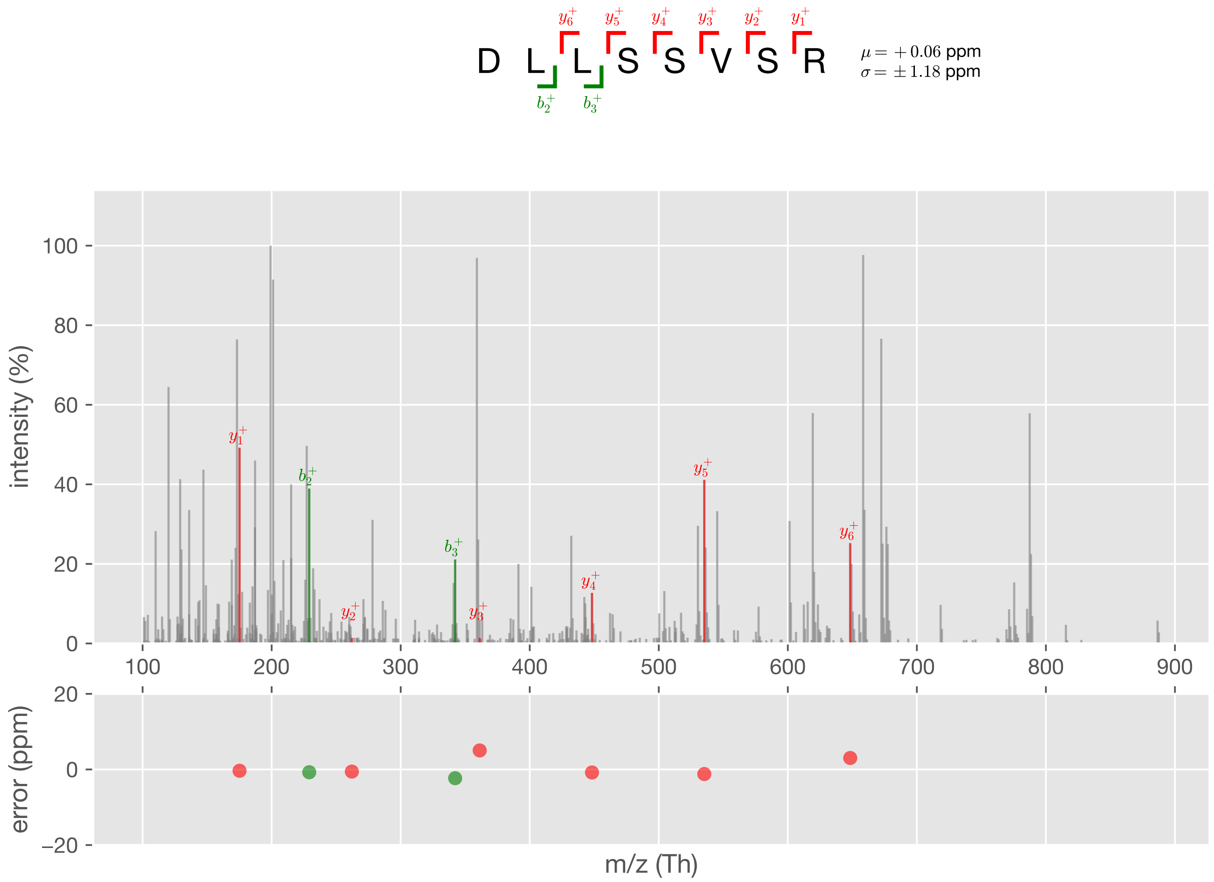


Figure S46: Case #4, PSM #1

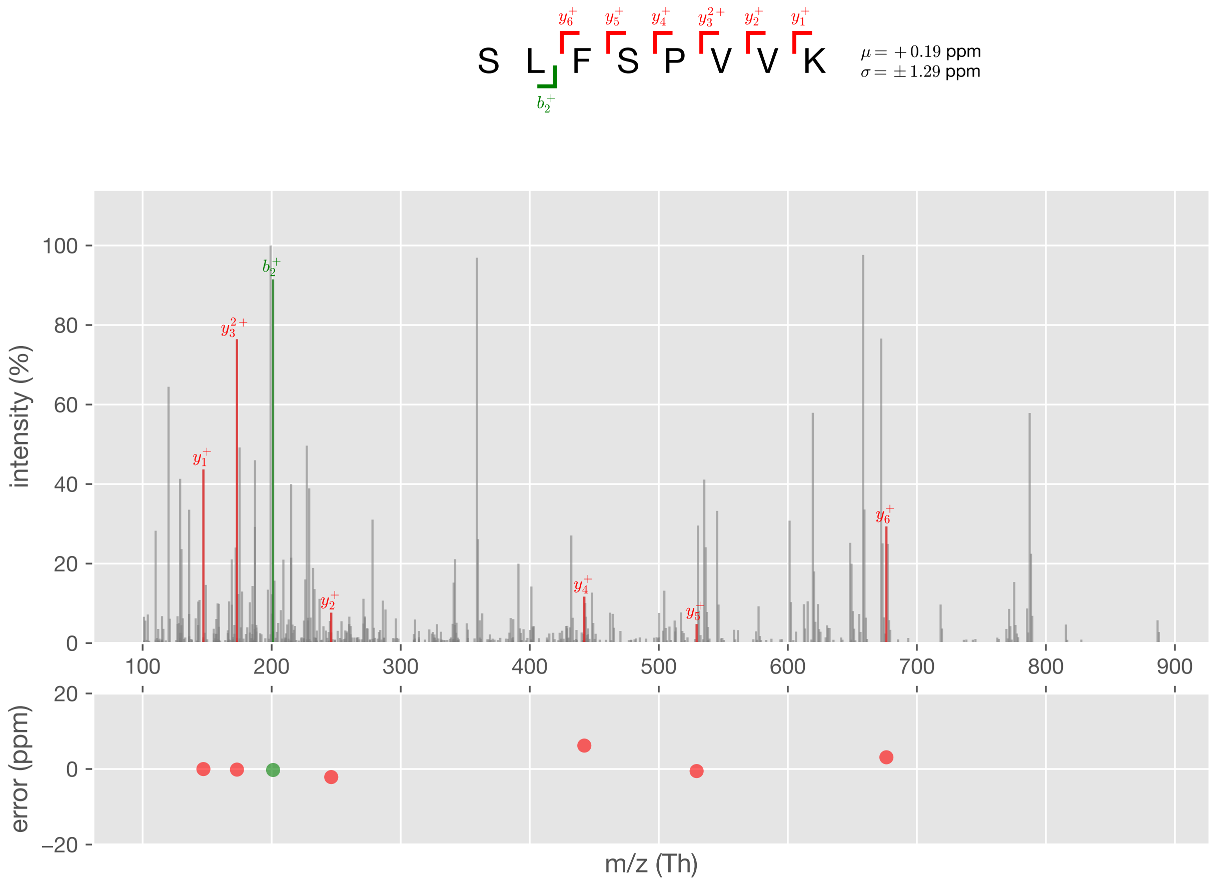


Figure S47: Case #4, PSM #2

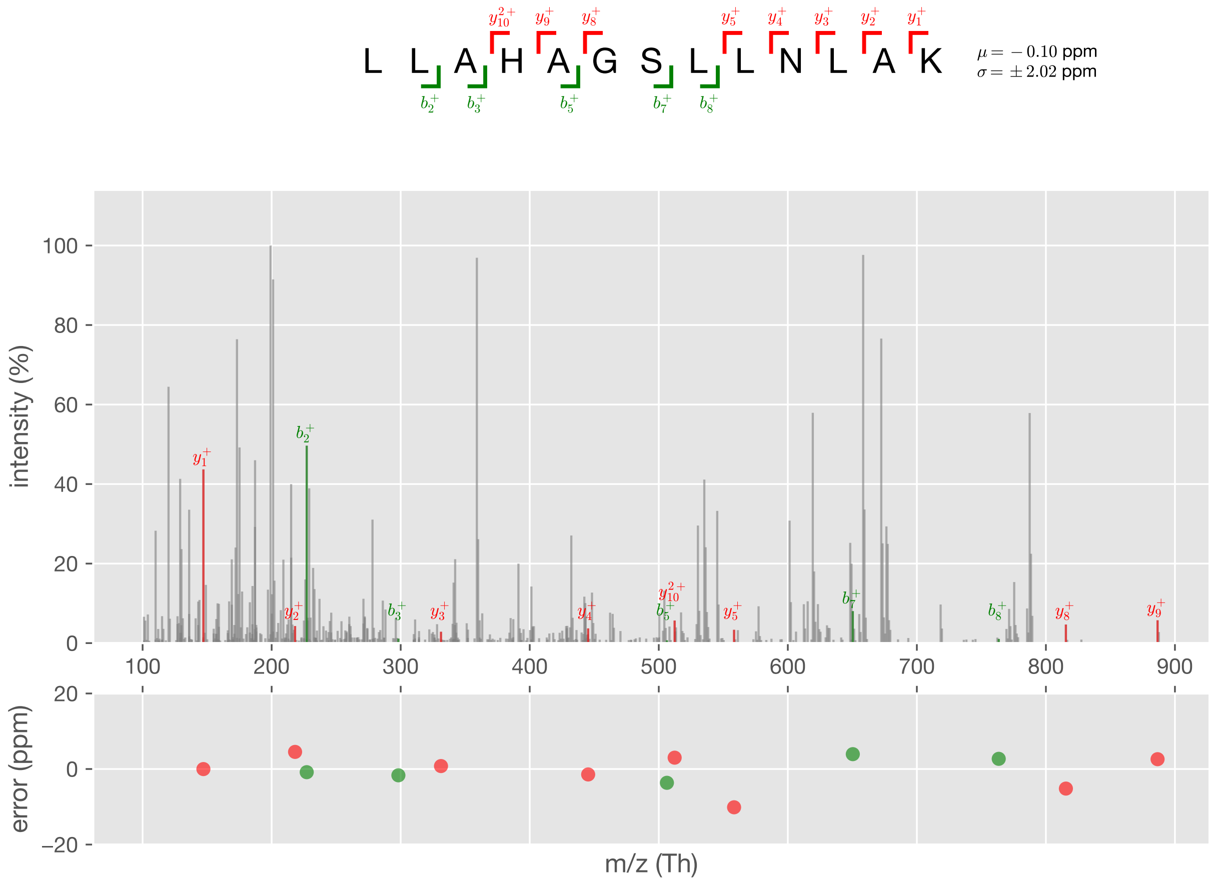


Figure S48: Case #4, PSM #3

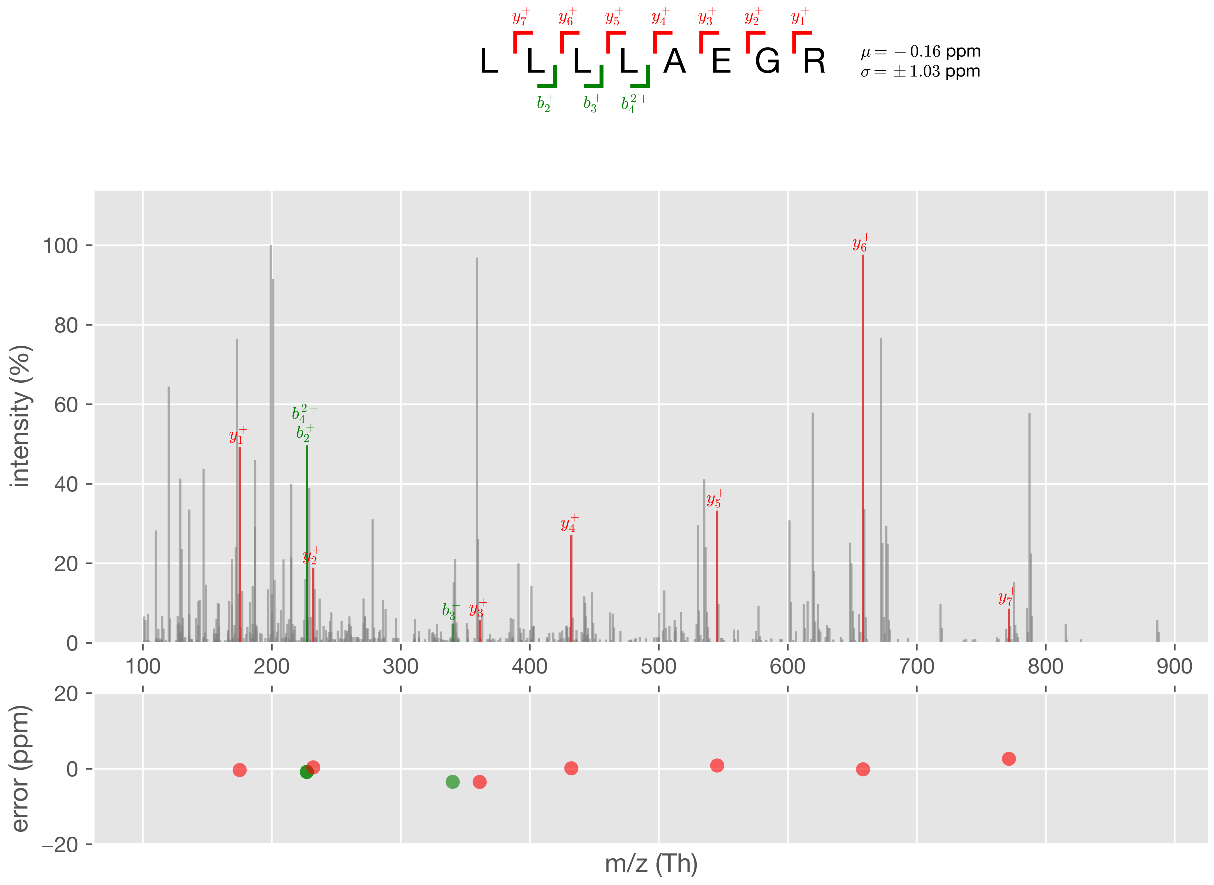


Figure S49: Case #4, PSM #4

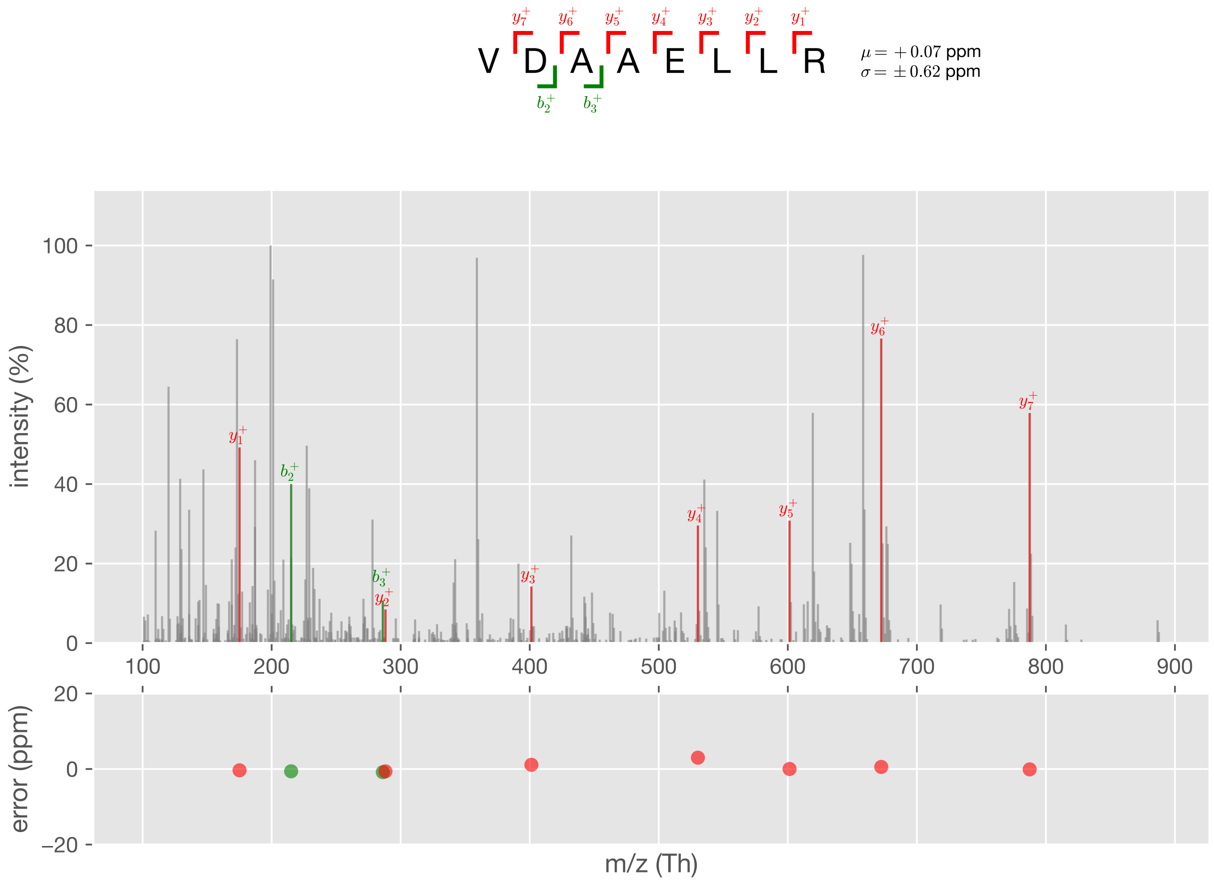


Figure S50: Case #4, PSM #5

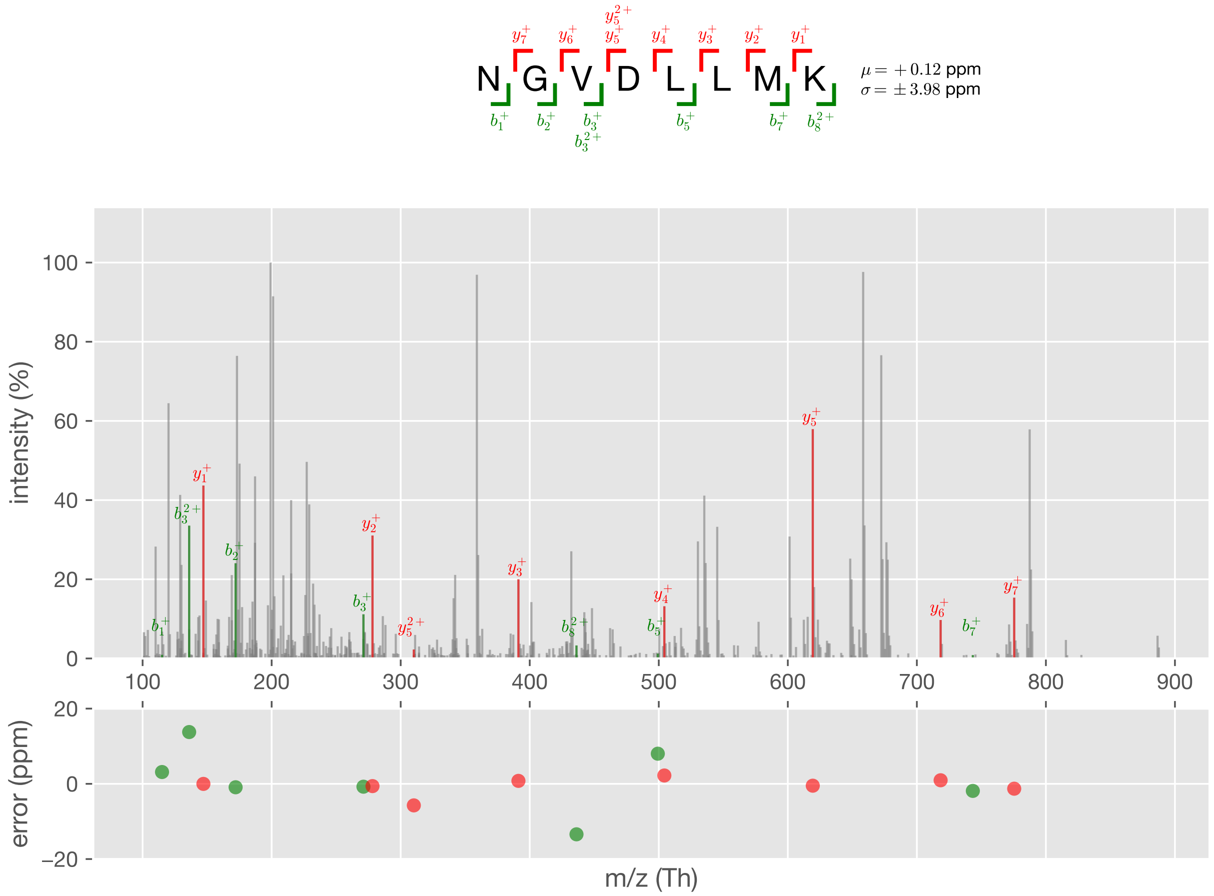


Figure S51: Case #4, PSM #6

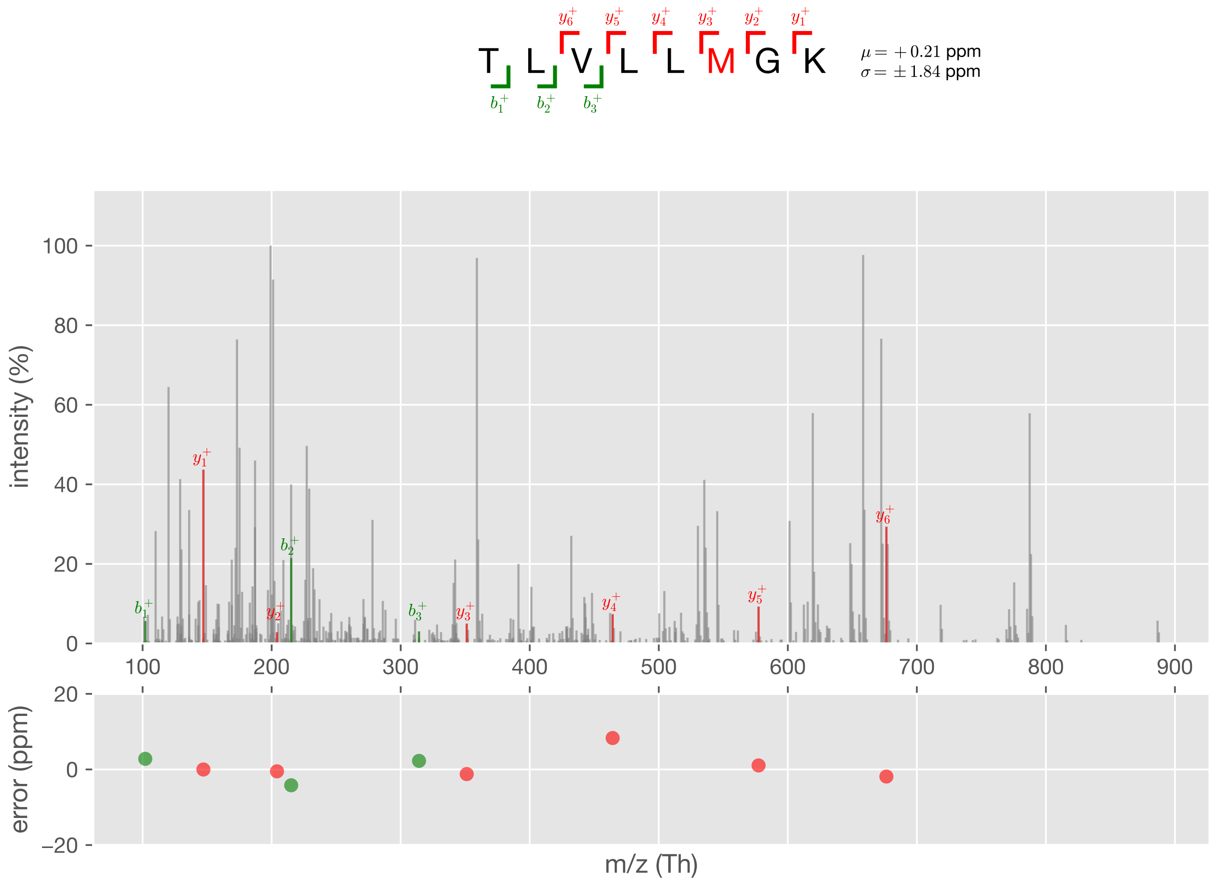


Figure S52: Case #4, PSM #7

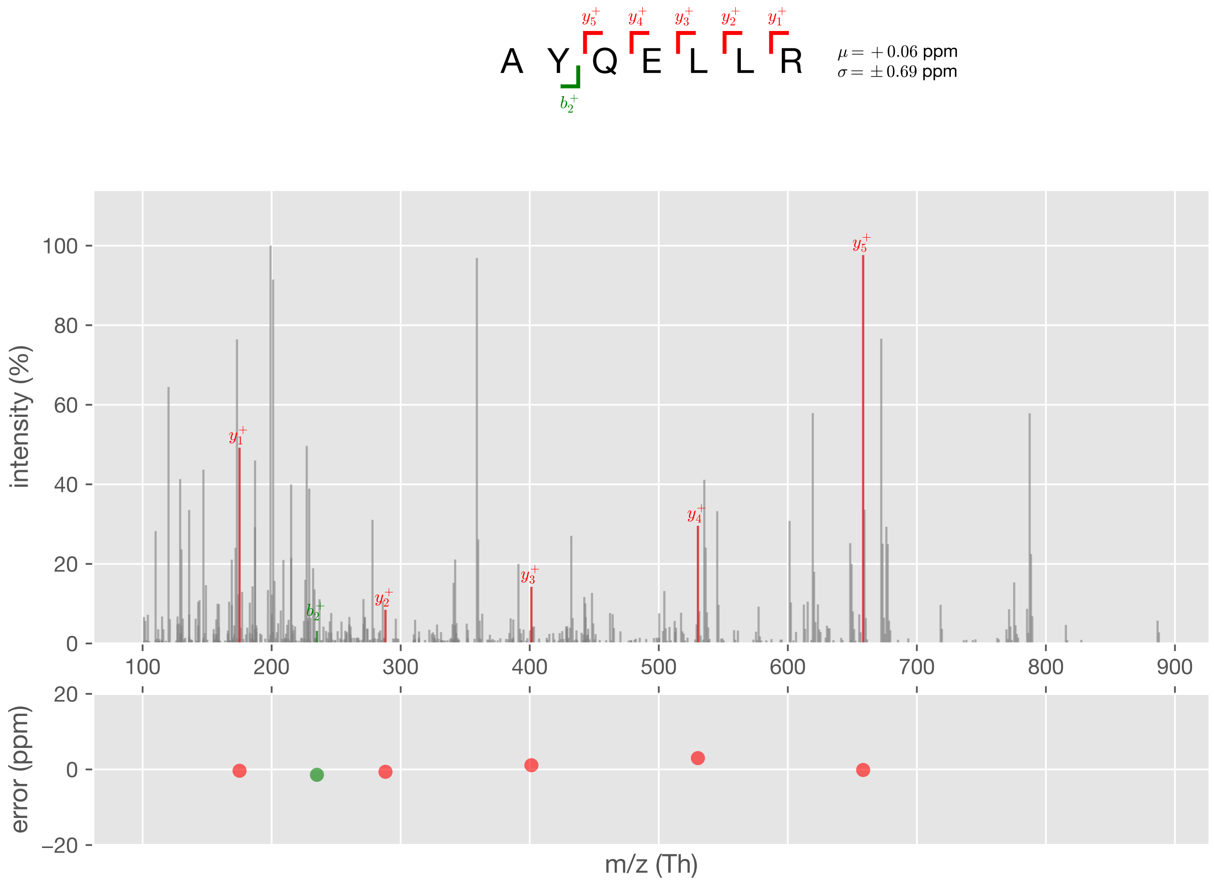


Figure S53: Case #4, PSM #8
