## Supplementary figures and images for "PepPre: Promote Peptide Identification Using Accurate and Comprehensive Precursors"

### Graphical User Interface of PepPre

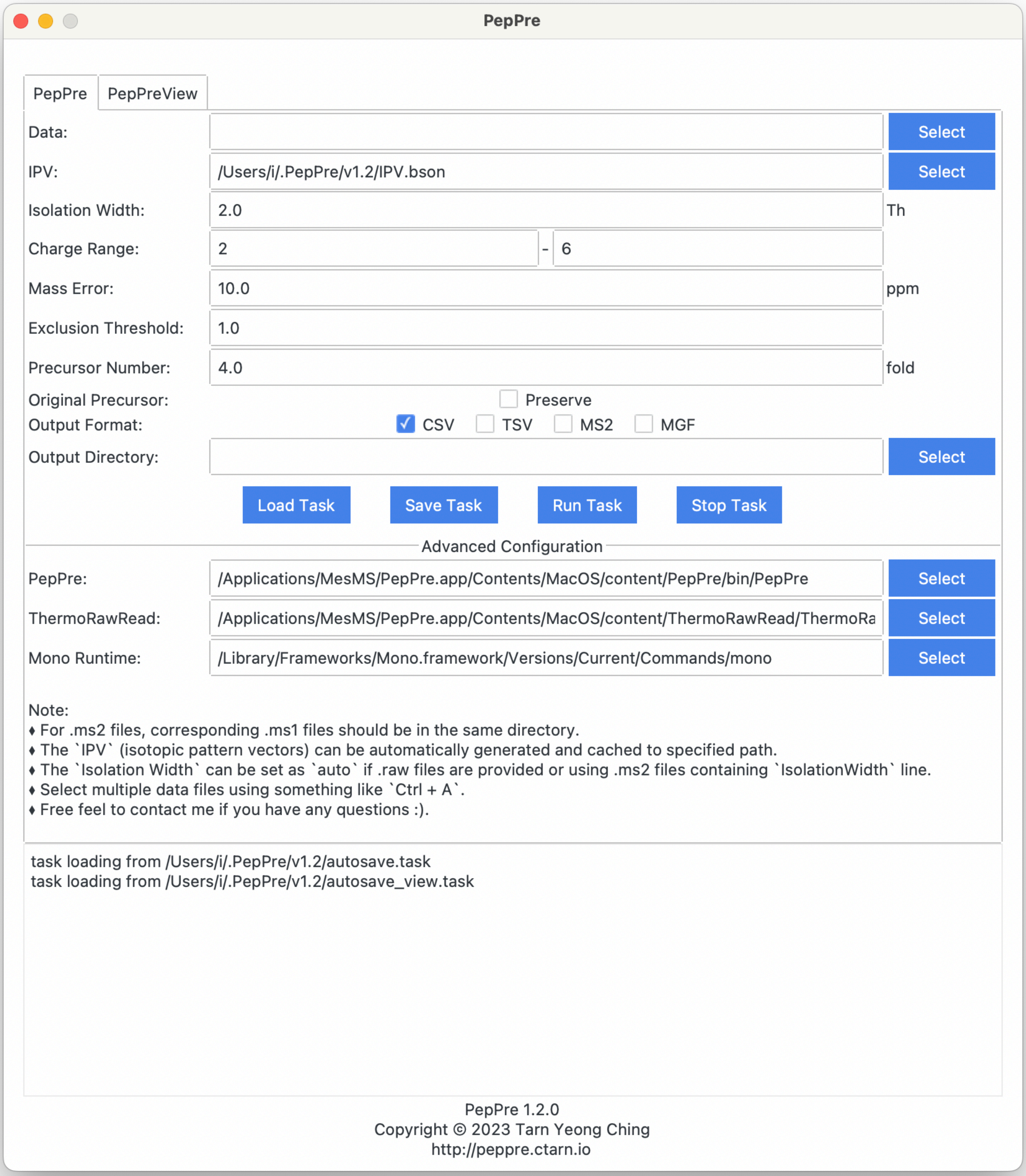
