## Supplementary material for "PepPre: Promote Peptide Identification Using Accurate and Comprehensive Precursors": Graphical User Interface of PepPreView

| RAW | Scan No. | Master Scan No. | Retention Time (s) | Activation Center (Th) | Isolation Width (Th) | #PSM |
|---|---|---|---|---|---|---|
| filter data... | | | | | | |
| 20131202_hela1_120min_8MZ | 19942 | 19937 | 3495.7587 | 484.73257446 | 8 | 9 |
| 20131202_hela1_120min_8MZ | 20952 | 20944 | 3617.9185 | 404.8793335 | 8 | 9 |
| 20131202_hela1_120min_8MZ | 36170 | 36158 | 5467.8134 | 507.74642944 | 8 | 9 |
| 20131202_hela1_120min_8MZ | 9124 | 9121 | 2174.7403 | 399.86538696 | 8 | 8 |
| 20131202_hela1_120min_8MZ | 9350 | 9345 | 2206.4353 | 407.21759033 | 8 | 8 |
| 20131202_hela1_120min_8MZ | 15630 | 15624 | 2954.0551 | 424.72940063 | 8 | 8 |
| 20131202_hela1_120min_8MZ | 15855 | 15850 | 2982.3767 | 480.2991333 | 8 | 8 |
| 20131202_hela1_120min_8MZ | 18535 | 18528 | 3320.5717 | 380.22128296 | 8 | 8 |
| 20131202_hela1_120min_8MZ | 19177 | 19167 | 3400.7005 | 379.2272644 | 8 | 8 |
| 20131202_hela1_120min_8MZ | 19542 | 19536 | 3445.5118 | 544.28375244 | 8 | 8 |

« ‹ 1 / 4805 › »

m/z error (ppm): 10

exclusion threshold: 1

score threshold: 0

charge state: 1+ 2+ 3+ 4+ 5+ 6+ 7+ 8+ 9+ 10+ 11+ 12+

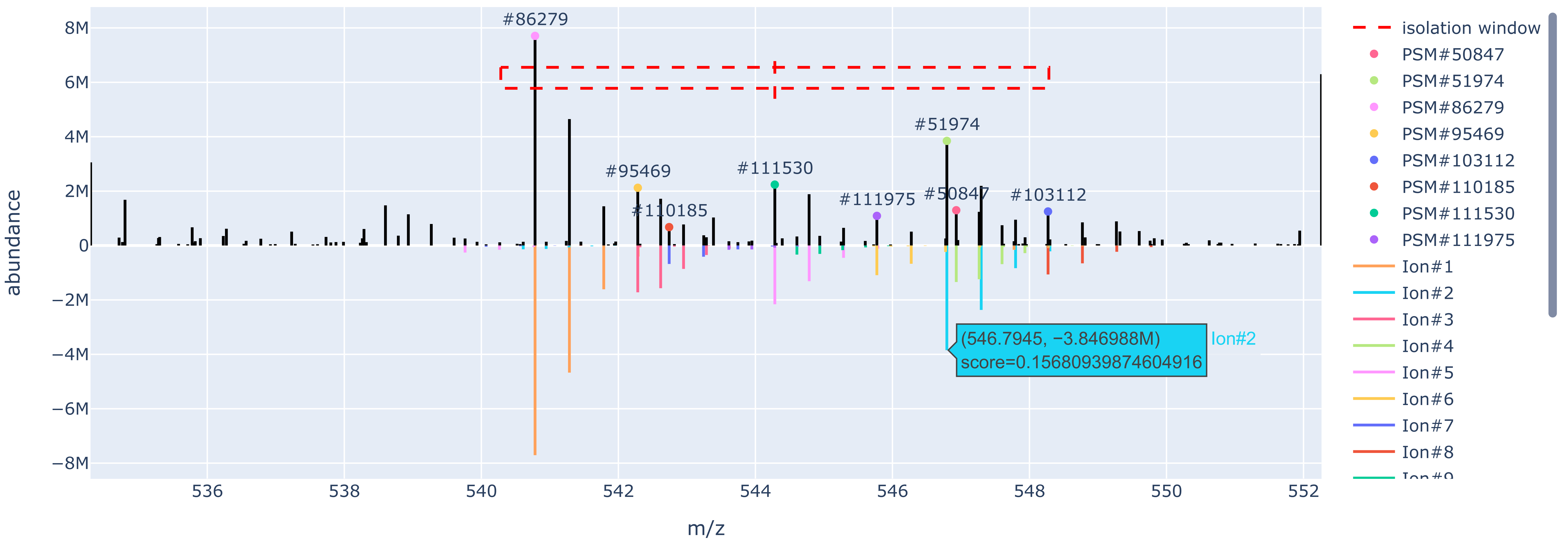


| PSM ID | M+H Mass (Da | m/z (Th) | Charge Stat | Sequence | Modification | Protein | Precursor Mass E |
|---|---|---|---|---|---|---|---|
| filter data. | | | | | | | |
| 50847 | 1638.779814 | 546.931455644414 | 3 | FNLTYVSHDGDDKK | | sp\|P26639\|SYTC_HUMAN/ | 2.47806 |
| 51974 | 1092.581525 | 546.7944007333106 | 2 | VHLELGPDGR | | sp\|P55795\|HNRH2_HUMAN/sp\|P52597… | 1.778356 |
| 86279 | 1080.56042 | 540.7838482333106 | 2 | YLAEVATGEK | | sp\|P61981\|1433G_HUMAN/ | 3.05674 |
| 95469 | 1624.836621 | 542.283724644414 | 3 | GSLTEGALAHSGNPVSK | | sp\|Q9UPQ0\|LIMC1_HUMAN/ | 4.78448 |
| 103112 | 1095.534819 | 548.2710477333105 | 2 | LTTDFGNAEK | | sp\|P02786\|TFR1_HUMAN/ | 2.91090 |
| 110185 | 1084.475373 | 542.7413247333105 | 2 | GYVSDLDCR | Carbamidomethyl[C]@8 | sp\|P48506\|GSH1_HUMAN/ | 2.42882 |
| 111530 | 1087.558686 | 544.2829812333106 | 2 | AAPSDGFKPR | Acetyl[AnyN-term]@1 | sp\|Q92979\|NEP1_HUMAN/ | 5.19512 |
| 111975 | 1090.539628 | 545.7734522333105 | 2 | SLETENAGLR | Deamidated[N]@6 | sp\|P02545\|LMNA_HUMAN/ | 2.0045 |

fragment m/z error (ppm): 20

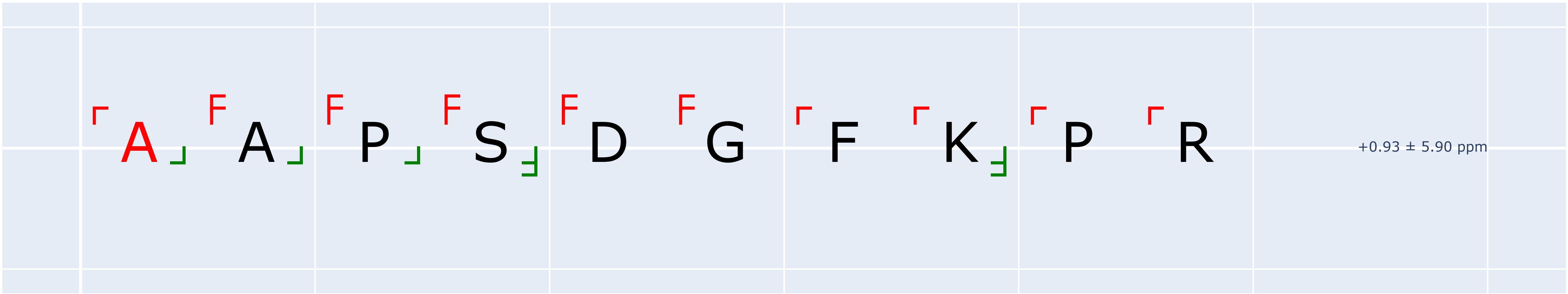


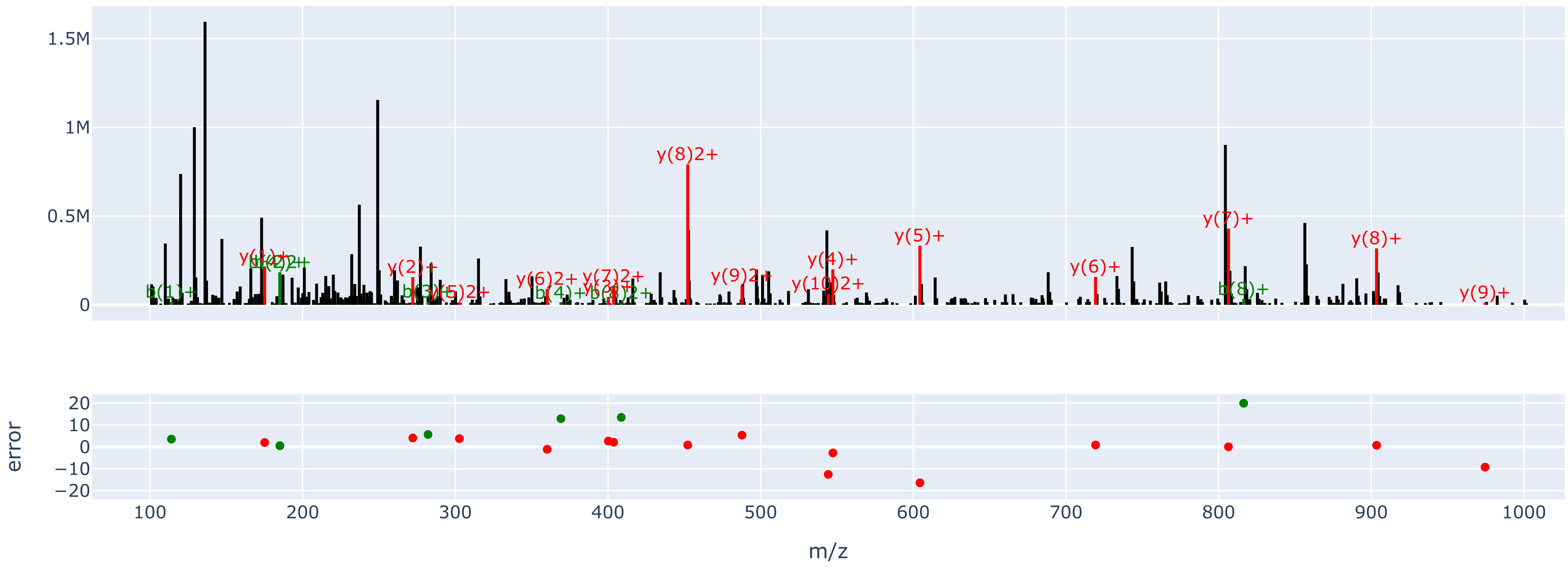
